# The abundances and occurrences of foliar microbes are poorly predicted by variation in plant traits and abiotic conditions

**DOI:** 10.1101/2022.05.20.492878

**Authors:** Joshua G. Harrison, C. Alex Buerkle

**Author notes:** Corresponding author : Joshua G. Harrison 1000 E. University Ave. Department of Botany, 3165 University of Wyoming Laramie, WY 82071, USA.

## Abstract

Much effort has been made to understand why foliar microbes live where they do. However, whether foliar microbiome composition can be predicted is unknown. Here, we determine the limits of prediction using metabarcoding data of both fungal and bacterial assemblages that occur within (endophytes) and without (epiphytes) leaves from 59 plant taxa. We built random forest models for prevalent taxa and quantified the combined predictive power of 24 plant traits, 12 abiotic conditions and 7 additional features. As response variables, we considered microbial relative and absolute abundances, and occurrences. Most microbial taxa were too rare to effectively model, but model performance was generally poor even for the most prevalent and abundant taxa (model *R*^2^ was typically <0.1). Fungi were more tractable for modeling than bacteria. Models of Shannon’s diversity were moderately successful but those for richness were not. Taxa responded idiosyncratically and non-linearly to variation in the foliar habitat. When prevalent microbes were included as features in models, performance improved. Our results suggest that easily measurable aspects of the phyllosphere habitat are poor predictors of microbiome composition. These results pose a challenge for the study of microbial biogeography and we discuss possible ways forward.

## Introduction

There is an astonishingly diverse array of microbiota living inside of leaves (henceforth endophytes) and on the surface of leaves (epiphytes; Arnold and Lutzoni 2007; Griffin et al. 2016; Lodge et al. 1996). Over the past decade, interest in these microbes has grown tremendously (Harrison and Griffin 2020), as motivated by the effects of endophytes on host plant traits, which can be quite dramatic (Doty 2011; Friesen et al. 2011) and likely scale up to influence entire ecosystems (Laforest-Lapointe et al. 2016).

A primary thrust of research has been to characterize the compositions of foliar microbiomes across various gradients—both biotic and abiotic—in an effort to determine drivers of community assembly and describe biogeographic patterns. Indeed, well over a thousand papers have linked variation in foliar microbiomes to a bewildering array of habitat characteristics, including rainfall (Lau et al. 2013), temperature (Oita et al. 2021), host taxon (Vincent et al. 2015), floral ‘neighborhood’ (Lajoie and Kembel 2021), the abundances of other microbes (Agler et al. 2016), insect herbivore activity (Humphrey and Whiteman 2020), the proximity of hosts to cities (Laforest-Lapointe et al. 2017), and even hail (Fernandes et al. 2011). These are but a smattering of examples—the list of interesting habitat-microbiome associations could fill the rest of this manuscript.

Most of these studies relied on composite response variables, meaning the response was neither the abundances nor occurrences of specific microbial taxa, but rather assemblage-wide richness, diversity, or estimates of divergence among samples (i.e., ordination-based analyses). Moreover, by logistical necessity, all studies have considered only a small subset of the numerous dimensions that together compose the foliar habitat. This has precluded accurate accounting of the relative importance of various plant traits and abiotic conditions for most foliar taxa. Indeed, whether or not the abundances of most taxa can be predicted by a consistent subset of habitat characteristics is unknown. A possible next step is to exhaustively characterize foliar habitat variation across host plant taxa growing in different conditions and to link habitat characteristics to the abundances of particular microbial taxa.

### Why the focus on prediction?

Predictive modeling of population dynamics has long-been a driving focus of applied population ecology, but community and microbial ecologists have tended to focus on pattern description (e.g., in species richness or diversity) and on ranking the importance of various processes—for example, those that mediate community assembly (Vellend 2010). Understanding process is akin to prediction, but not quite the same. Even if the relative importance of processes behind community assembly are understood, there may be stochastic forces at work that have largely unpredictable outcomes or chaotic, deterministic forces that are unpredictable without knowledge of antecedent conditions (May 2019). A further challenge is that various ecological forces may leave indistinguishable imprints in natural communities, precluding accurate quantification of process outside of a manipulative context. This suggests that accurate prediction is likely harder than accurately understanding process—a daunting prospect indeed.

A countering argument could be made that assessing the relative importance and contributions of confounded processes may be impossible, but prediction of various, useful attributes of ecological communities could be within reach. Indeed, at the macro-ecological scale this has proven true as many patterns can be reliably observed and thus predicted, but the relative importance of the myriad processes underlying them often remain obscure—the latitudinal gradient in species richness is a good example (Mittelbach et al. 2007; Pianka 1966). Similarly, patterns in assemblage diversity, richness, and perhaps even the relative abundances of specific taxa could be predictable by various measurable phenomena, be they causal or merely correlated. Indeed, the literature is built upon studies that explain some variation in community composition in this way (see the examples cited above), though the variance explained in response variables often is quite low. Under the predictive paradigm, the question becomes: what are the limits of predictive modeling, how can models be optimized, and what biology can be learned? We do not mean to suggest that prediction and the interrogation of process are necessarily at odds. Indeed, once the limits of prediction are known a better accounting of underlying process may be possible.

To illustrate the possible benefits of the predictive paradigm, consider two possibilities for the analyses presented here. It may be that through an comprehensive measurement of plant traits and abiotic characteristics we can account for a large proportion of the variation in the abundances of many microbial taxa—say 50% or more of the variation in the abundance of the 500 most common microbes (while acknowledging the vagueness of the word ‘common’). In this case, while parsing deterministic and stochastic forces may still be challenging, it suggests there is a firm basis for applied research and for future experimentation to discover causal mechanisms. On the other hand, if it is not possible to predict more than a few percent of the variation in abundances of common microbes then either the hypothesized deterministic forces that were measured are neither causal nor covary with causal phenomena or that stochastic and chaotic forces drive the observed variation. This situation would also suggest possible ways forward, through measuring different habitat dimensions that could be causal, or by developing better measurement tools, experiments, or theory to determine rates of stochastic community divergence.

### Modeling microbiota: tools and challenges

Most studies linking foliar microbiota to habitat conditions rely on linear modeling or some analysis of the variation within a distance (or dissimilarity) matrix made from taxon counts within samples (Bowman and Arnold 2021; Gomes et al. 2018; González-Teuber et al. 2020; Kembel et al. 2014; Kembel and Mueller 2014; Oita et al. 2021; Vincent et al. 2015). Linear modeling has many benefits, including intuitive interpretation of model coefficients and a resistance to overfitting. However, it may not perform well when predicting non-linear phenomena. Analyzing variation within distance matrices, typically via some combination of PERMANOVA and ordination, also has benefits, including the ease with which intuitive visualizations can be made of model results. But this approach is limited because it relies on describing differences in centroids of points that lie in a space with few dimensions (usually two or three). Thus, covariances among many thousands of organisms are decomposed into a few dimensions (i.e., eigenvectors) that may only explain a small percentage of the overall variation within the matrix and then those dimensions linked to habitat variation (sometimes qualitatively, via visual inspection of ordination plots). This technique provides an estimate of ‘community structure’ (i.e. patterns of among-sample similarity in assemblage composition) but provides no taxon-specific insights. Moreover, decisions must be made regarding the choice of matrix decomposition, which determines the relative weight of rare and abundant taxa on the analysis and that shapes inferences (Legendre and Gallagher 2001).

In comparison, machine learning methods, as a broadly-defined suite of approaches, include many algorithms that are optimized for prediction, can readily handle non-linear relationships, and that do not necessarily rely on distributional assumptions. As such, the application of machine learning methods to foliar microbiome data should provide novel insights. Here, we use the random forest algorithm because it has interpretable outputs (i.e., the ranking of feature importance), is easy to implement, and provides very strong predictive performance (Breiman 2001; Cutler et al. 2007). Random forests are a collection of decision trees that split the observations (e.g., taxon abundances) into sets according to values of covariates that are each made using a subset of the available data and covariates (henceforth referred to as features, as is typical within the machine learning literature). The entire ensemble of trees constitutes the forest and prediction is accomplished through aggregating and averaging the individual outputs of each tree. Feature importance is typically determined via a post-hoc perturbation test, where a feature is permuted, the model retrained, and the change in performance recorded.

Regardless of the analytical approach employed, microbiomes present two hurdles that complicate predictive modeling. First, most microbes are rare, meaning they may be represented by only a few sequence reads within several samples. Since rare taxa are so infrequently observed it is typically impossible to model their relationship with covariates— unless, the unlikely scenario occurs of a rare taxon being associated with some similarly unusual, but measurable, aspect of the foliar habitat and a large enough sample size has been obtained such that the associated phenomenon can be observed multiple times. Second, it is harder to accurately count microbial taxa than it is to count macroscopic organisms. To be counted, microbial taxa have to be cultured and colony-forming units tallied, a valuable but often logistically challenging endeavor (Carini 2019); or viewed within a sample via microscopy or flow cytometry, tools that have not yet been used at scale to ask questions pertaining to microbial biogeography; or characterized via sequencing of DNA. For the latter approach, the sequences of marker loci that vary among taxa are characterized. Different sequences are assigned to different taxa and referred to as operational taxonomic units (OTUs). The number of counts output by the sequencing machine for a particular sequence describes that taxon’s relative abundance within the sample. While this approach to characterizing microbiomes is appealingly cost and time-effective, various laboratory biases must be considered (Nilsson et al. 2018) and, even when all is well in the lab, the resulting data suffer from the limitation that they describe only relative abundances. That is, sequence count data are compositional in nature. Compositional data are interdependent, as sequence counts for one taxon increases those for another taxon must decrease (Jackson 1997; Tsilimigras and Fodor 2016). This limitation is imposed by the instrumentation because sequencing machines output a finite number of reads.

To circumvent this challenge, internal standards (ISDs) can be added to samples prior to sequencing. Since the amount of ISD is standardized among samples, division of sequence count data by the ISD places the data on the same scale, which is proportional to abundance (henceforth referred to as “absolute” abundance; Tourlousse et al. 2017). The benefits of ISDs can be undercut (as reviewed by Harrison et al. 2021c), but their use can lead to novel inference.

### The limits of prediction for foliar microbiota

Here, we determine to what extent the abundances and occurrences of foliar taxa are predictable and identify the most influential characteristics of the foliar habitat for fungal (ITS) and bacterial (16s) endophytes and epiphytes. We sequenced DNA from 1241 individual plants collected from the mountain ranges of Wyoming, U.S.A. (Fig. 1). When building models, we considered 24 plant traits, 12 abiotic site characteristics (e.g., rainfall, elevation, etc.), aspects of the vegetation matrix surrounding the site, and those metrics pertinent to sampling, such as the mass of sampled leaves (Table 1). We ask to what extent model performance is shaped by the choice of dataset: either relative or absolute abundance of taxa or presence and absence data for occurrence. Finally, we measured the limits of prediction for common derived variables, including Shannon’s diversity and richness.

**Figure 1:**
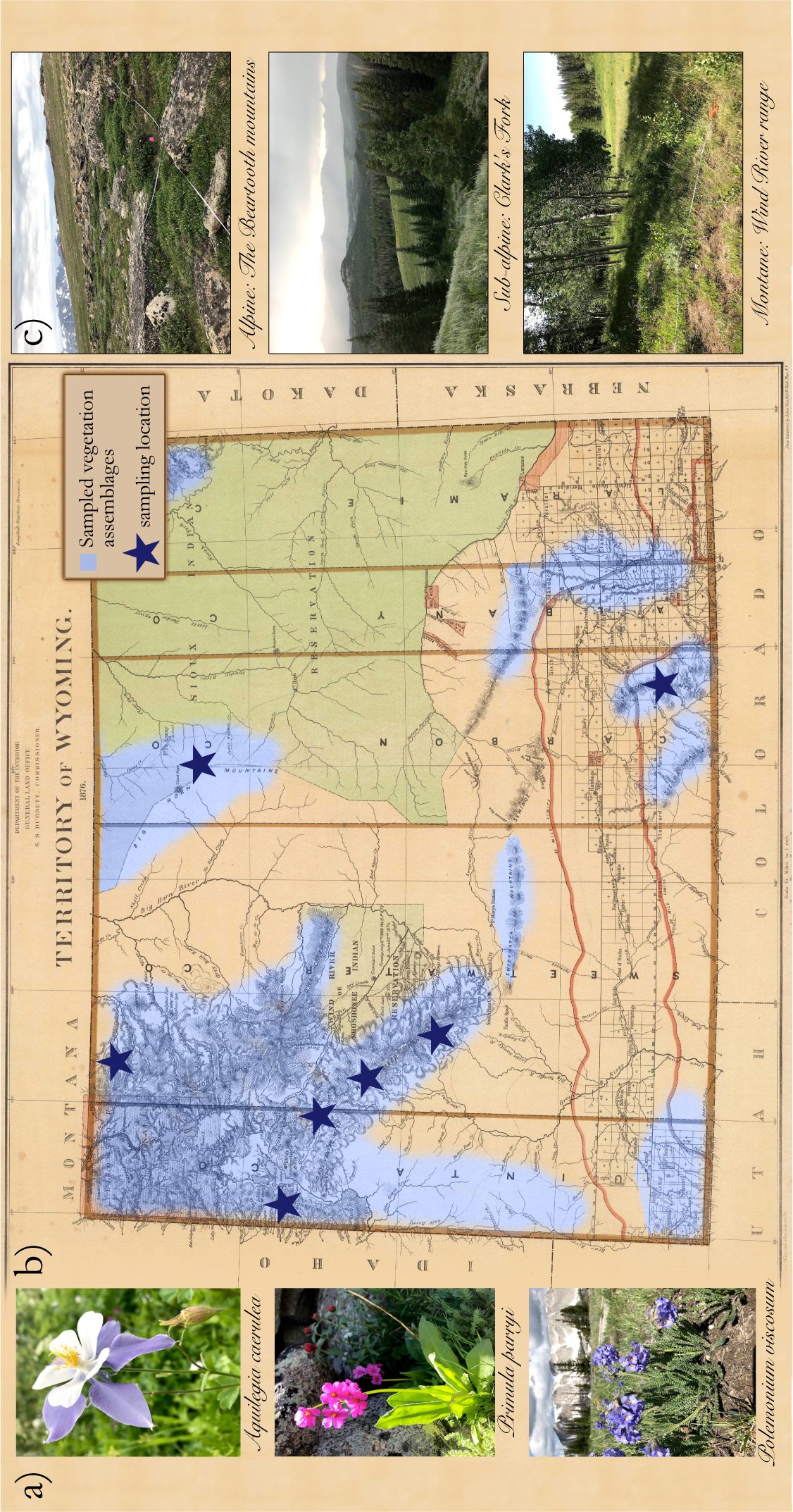
Example plant taxa sampled (a) and sampling locations in Wyoming, U.S.A (b). Each star denotes a sampling region—at each region, sampling took place at three different elevations. Sites were chosen within the alpine, sub-alpine and montane vegetation zones (examples shown in c). The portion of the map shaded blue approximately denotes the geographical extent of the vegetation types surveyed. Plant taxa sampled included dominant trees, shrubs, and forbs, along with interesting less common plants, with the goal of sampling ∼80% of above-ground biomass. The map shown is from 1876, several years before Wyoming became a state. Political jurisdictions of the day are shown along with the portion of the state occupied by the Union-Pacific railroad (delineated in pink lines at the bottom25of the map). All photographs by J. G. Harrison; map is part of the public domain.

**Table 1:** Predictor variables (features), including plant traits and abiotic conditions, measured for this study. Details of measurement and citations explaining the method or supporting the possible influence of the feature are shown. We apologize to the many scientists whose work was omitted due to space limitations.

| Predictor variable | Description | Citations |
| --- | --- | --- |
| Absorbance of leaves at 940 nm | Measured using the PhotosynQ multispeQ handheld spectrophotometer. | (Carvalho and Castillo 2018; Sanchez-Azofeifa et al. 2012) |
| Ambient humidity | Relative humidity (%) | (Aung et al. 2018) |
| Area | Leaf area measured in cm<sup>2</sup> | (Kinkel et al. 1987) |
| Circumference of stem | Circumference of trees 1.5 m from the ground | (Yu et al. 2021) |
| Compartment | Endophyte (EN) or epiphyte (EP). Putative endophytes were obtained from washed leaves, while epiphytes were obtained from the liquid used to wash leaves | (Coleman-Derr et al. 2016) |
| Date | Julian date of sampling, starting from Jan 1 | (Arnold and Herre 2003) |
| Dead & down | The number of dead trees lying on the ground at the site | Decaying wood could be an inoculum source. |
| Densiometer | Densiometer measurements above each of the four floral composition quadrants, ranges from 0–4, average taken for each site | An index of canopy closure at the site. |
| Electrochromic shift (ECS) initial | Measurement of ATP synthase activity. Measured using the PhotosynQ multispeQ handheld spectrophotometer. | (Cruz et al. 2001; Kramer et al. 1999; Sacksteder et al. 2000) |
| Elevation | Measured in meters | (Zimmerman and Vitousek 2012) |
| Fs | Light-adapted steady state fluorescence. Measured using the PhotosynQ multispeQ handheld spectrophotometer. | (Genty et al. 1989) |
| gH | Proton conductivity. Measured using the PhotosynQ multispeQ handheld spectrophotometer. | (Kanazawa and Kramer 2002) |
| Habit | Growth habit, either shrub, forb, graminoid, or tree | (Harrison and Griffin 2020) |
| Height of sample | Height in meters above ground of the sampling location on the plant | (Harrison et al. 2016) |
| Host taxon | Specific epithet of the host plant | The influence of host has been found in all papers of which we are familiar, though it can be quite weak (Vincent et al. 2015). |
| Latitude | Latitude of site | (Arnold and Lutzoni 2007) |
| Longitude | Longitude of site | (Wu et al. 2013) |
| Leaf temperature differential | Contactless temperature (leaf surface temperature) minus ambient temperature. Measured using the PhotosynQ multispeQ handheld spectrophotometer. | – |
| Leaves extracted | Number of leaves or leaflets from which DNA was extracted | (Arnold et al. 2000) |
| Linear electron flow (LEF) | Measurement of the movement of electrons from water to NADP+ during photosynthesis. Measured using the PhotosynQ multispeQ handheld spectrophotometer. | (Genty et al. 1989) |
| Light intensity PAR | Photosynthetically active radiation. Fraction of light (400–700nm) important for photosynthesis. Measured using the PhotosynQ multispeQ handheld spectrophotometer. | (Carvalho and Castillo 2018; Fahimipour et al. 2018) |
| Mass extracted | Grams of leaf material from which DNA was extracted. | (Kinkel et al. 1987) |
| Moran's eigenvector map (MEM) 1, 2 | Two MEMs created from latitude and longitude data of each site. | (Bowman and Arnold 2021) |
| Phenological status | Categorical. One of fruiting, vegetative, flowering. | (Cook et al. 2012) |
| Phi NO | Non-regulatory energy dissipation. Measured using the PhotosynQ multispeQ handheld spectrophotometer. | — |
| Phi NPQ | Measurement of non-photochemical quenching. Measured using the PhotosynQ multispeQ handheld spectrophotometer. | — |
| Plant volume | Volume of a box that could fit over the plant | (Rho et al. 2018) |
| Precipitation | Mean precip. during April (the month prior to the start of sampling) | (Zimmerman and Vitousek 2012) |
| qL | Photosystem II redox state from the “lake” model. Measured using the PhotosynQ multispeQ handheld spectrophotometer. | (Kramer et al. 2004) |
| Relative chlorophyll intensity | A parameter derived by the multispeQ describing absorbance intensity of chlorophyll. | (Sanchez-Azofeifa et al. 2012) |
| Relative chlorophyll | A parameter derived by the multispeQ from absorbance measurements at 650 and 940 nm that provides an estimate of chlorophyll concentration | (Sanchez-Azofeifa et al. 2012) |
| Shannon's veg. diversity | Vegetation diversity from Daubenmire plots within each site | (Griffin and Carson 2018; Lajoie and Kembel 2021) |
| Shrub richness | The number of shrub species at the site | (Griffin and Carson 2018; Lajoie and Kembel 2021) |
| Specific leaf area (SLA) | Leaf area divided by leaf mass | (Knoth et al. 2013; Liu et al. 2019) |
| Slope percentage | The slope of the site measured in percent | — |
| SPAD intensity at 420 | Special Products Analysis Division (SPAD) - estimation of relative chlorophyll content from absorbance measurements. SPAD is a constant times relative chlorophyll content. This constant is defined by the multispeQ and approximates a proprietary constant from a Konica-Minolta chlorophyll measurement device. Because of high correlation of the measurement for different wavelengths only 420 nm was used. | (Sanchez-Azofeifa et al. 2012) |
| Temperature | Ambient temperature in degrees Celsius. Measured using the PhotosynQ multispeQ handheld spectrophotometer. | (Zimmerman and Vitousek 2012) |
| Temperature one month prior | Average temperature in April, the month before sampling started | (Zimmerman and Vitousek 2012) |
| Time of day | Sampling time | (Hubbard et al. 2017) |
| Thickness | Leaf thickness in $\mu\text{m}$ | (González-Teuber et al. 2020) |
| Leaf toughness | Measured in g with Chatillon penetrometer | (Arnold and Herre 2003) |
| Tree richness | Number of tree species within plot | (Griffin and Carson 2018; Lajoie and Kembel 2021) |
| Water retention | Ability of a leaf to hold a water droplet as a function of leaf angle, measured in degrees. | (Doan et al. 2020) |

## Methods

### Sampling details

Sampling took place during the summer of 2018 in six mountain ranges in Wyoming, U.S.A. (Fig. 1). Within each locality 2–3 50m by 50m plots were selected. At each locality, plots were selected in the alpine, sub-alpine, and lower sub-alpine/montane forest. The elevation range of sampling locations spanned 2120–3419m. A total of twenty plots were sampled. A total of 59 plant species were sampled. We attempted to collect 10 individuals from each plant species at each site. We obtained leaves or leaflets from 1241 plants. For details regarding field protocols, including how plant traits were measured, see the Supplemental Material.

### Sample preparation, sequencing, and bioinformatics

To separate endophytes from epiphytes, leaves were placed in tubes and agitated in a solution of 1×PBS, pure water, and 0.15% Tween 20 for 20 minutes and then sonicated for 5 min. The solution was decanted, centrifuged, and lyophilized and constituted our epiphyte (EP) samples (see the Supplemental Material). Washed leaves were lyophilized and ground in a mixer mill.

DNA was extracted using Qiagen DNEasy plant kits. Library preparation followed the two-step PCR procedure described in Harrison et al. (2021a). To amplify the ITS region, the ITS1f-ITS2 primer pair was used and for the 16S locus the 505-806 pair was used (Wang and Qian 2009; White et al. 1990). We added an equimolar amount of a synthetic DNA internal standard (ISD) to each sample prior to PCR (Tourlousse et al. 2017). Negative controls, including for cross-contamination, and positive controls were employed during library preparation and sequencing. Psomagen, Inc. (Rockville, MD, USA) performed paired-end 2*×*250 sequencing using an Illumina NovaSeq machine.

For details of bioinformatics see the Supplemental Material. In brief, exact sequence variants (ESVs) were determined and clustered by 97% similarity into OTUs. Taxonomic hypotheses for OTUs were generated using the SINTAX algorithm (Edgar 2016) and the UNITE (for ITS; v7.2; Community 2017) and Greengenes (for 16S; v13.5; DeSantis et al. 2006) databases.

Prior to modeling, count data were either Hellinger standardized (for relative abundance datasets) or normalized using the internal standard, thus putting all taxa on an equivalent scale and avoiding many of the challenges posed by compositionality (Harrison et al. 2021c). For ordinations and cluster analyses, data were Hellinger standardized and the Euclidean distance was calculated (Legendre and Gallagher 2001). Richness was estimated using the breakaway R package (Willis and Bunge 2015), which uses non-linear regression of the ratios of taxon frequencies to estimate richness within samples.

### Predictive modeling approach

To determine the predictability of microbial abundances we used random forest models implemented using the ranger R package (v 0.13.1 Wright and Ziegler 2017). We modeled four sets of response variables. First, we modeled the absolute and relative abundances of prevalent microbial taxa, which were those that occurred in 100 or more individual plants (about 10% of samples). 172 fungal and 26 bacterial OTUs met this prevalence threshold. Second, we repeated this analysis using qualitative data for occupancy. Third, we asked if predictive ability shifted when examining the microbiome of a single host, again we used absolute and relative abundances and occupancy data. Fourth, we modeled Shannon’s diversity and estimated richness for both fungi and bacteria.

We used a rigorous approach to model fitting, tuning, and performance determination that was reliant upon the mlr3 R package (v 0.12.0; Lang et al. 2019).

Models of ISD transformed abundances were repeated while including the abundances of other prevalent microbes as features. These microbes were the same as those chosen for modeling (see above) and thus represented the most abundant and prevalent taxa. For models of bacteria, only co-occurring bacteria were used as features and the same was true for fungal models. The purpose of this analysis was to determine if microbial abundances could be predicted by the abundances of co-occurring taxa. We conducted this analysis using ISD transformed data only to avoid spurious results due to compositionality.

## Results

### Biodiversity and general sequencing results

After filtering reads, removing non-target taxa (e.g., the host plant), and removing quality control sequences, we retained 12,795,691 ITS reads and 5,733,638 16s microbial reads for analysis (notably we obtained ∼13 million additional plant reads from our 16S data that were discarded; Fig. S6). ESVs among these reads were identified and clustered (97% similarity threshold) into 3189 fungal OTUs and 2360 bacterial OTUs.

Most microbial taxa were observed in few plant hosts and were low abundance. Only 172 fungal taxa and 23 bacterial taxa were present in more than 100 of the individual plants we sampled (out of 1241 total plants). More abundant taxa tended to also be more prevalent for both bacteria (Pearson’s correlation of median abundance and prevalence, r = 0.49, p < 0.001) and fungi (r = 0.49, p < 0.001). The strength of this correlation increased when only considering those taxa that occurred in 100 or more samples (fungi: r = 0.5; bacteria: r = 0.76, p < 0.001 in both cases).

Fungi tended to be more abundant inside rather than outside of leaves, but the opposite was true for bacteria (Fig. 2). Many microbial taxa occurred as both epiphytes and endophytes; indeed, all fungi that occurred as epiphytes occurred as endophytes (Fig. 3d). However, there were many more microbial taxa that were observed solely in epiphyte samples, suggesting that our leaf washing protocol to capture epiphytes was successful. The leaf barrier seemed to shape bacterial assemblages more than it did for fungi as we observed greater correlation in microbiomes between plant compartments (EP versus EN) for the latter (Mantel test of Hellinger transformed distance matrices; for bacteria: r = 0.11, p < 0.01 and for fungi: r = 0.36, p < 0.01).

**Figure 2:**
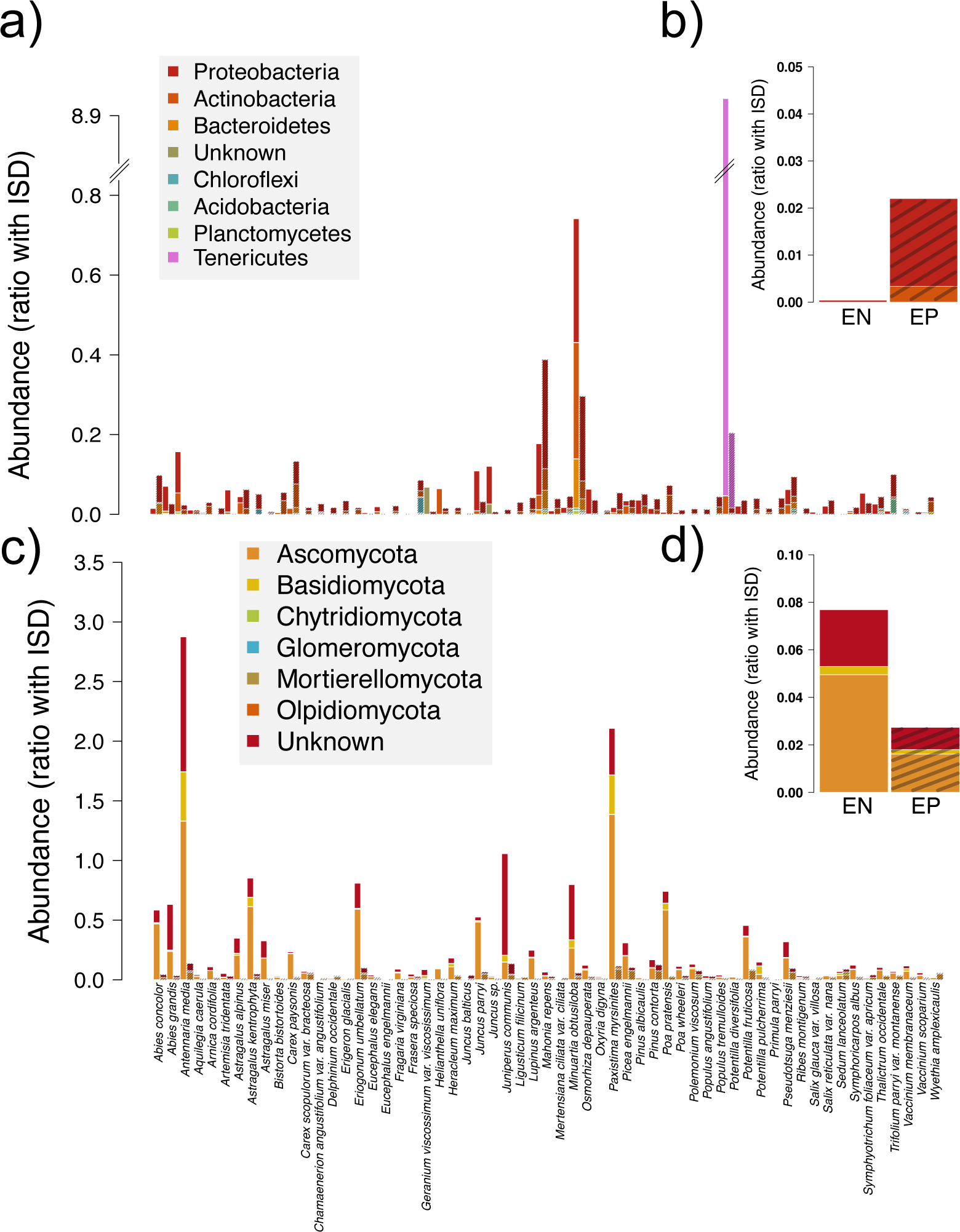
Biodiversity of phyllosphere microbes across 59 plant taxa collected from the mountain ranges of Wyoming, U.S.A. Hash marks are superimposed on epiphyte data. Data shown are median estimates of the relative abundances of phyla across samples. An internal standard (ISD) was included in each sample and used to place abundances on a standard scale (see main text). Panel a) shows abundance by phylum and host taxon for bacteria. Panel b) shows abundance by phylum and compartment for bacteria. Panels c) and d) repeat this motif for fungi.

**Figure 3:**
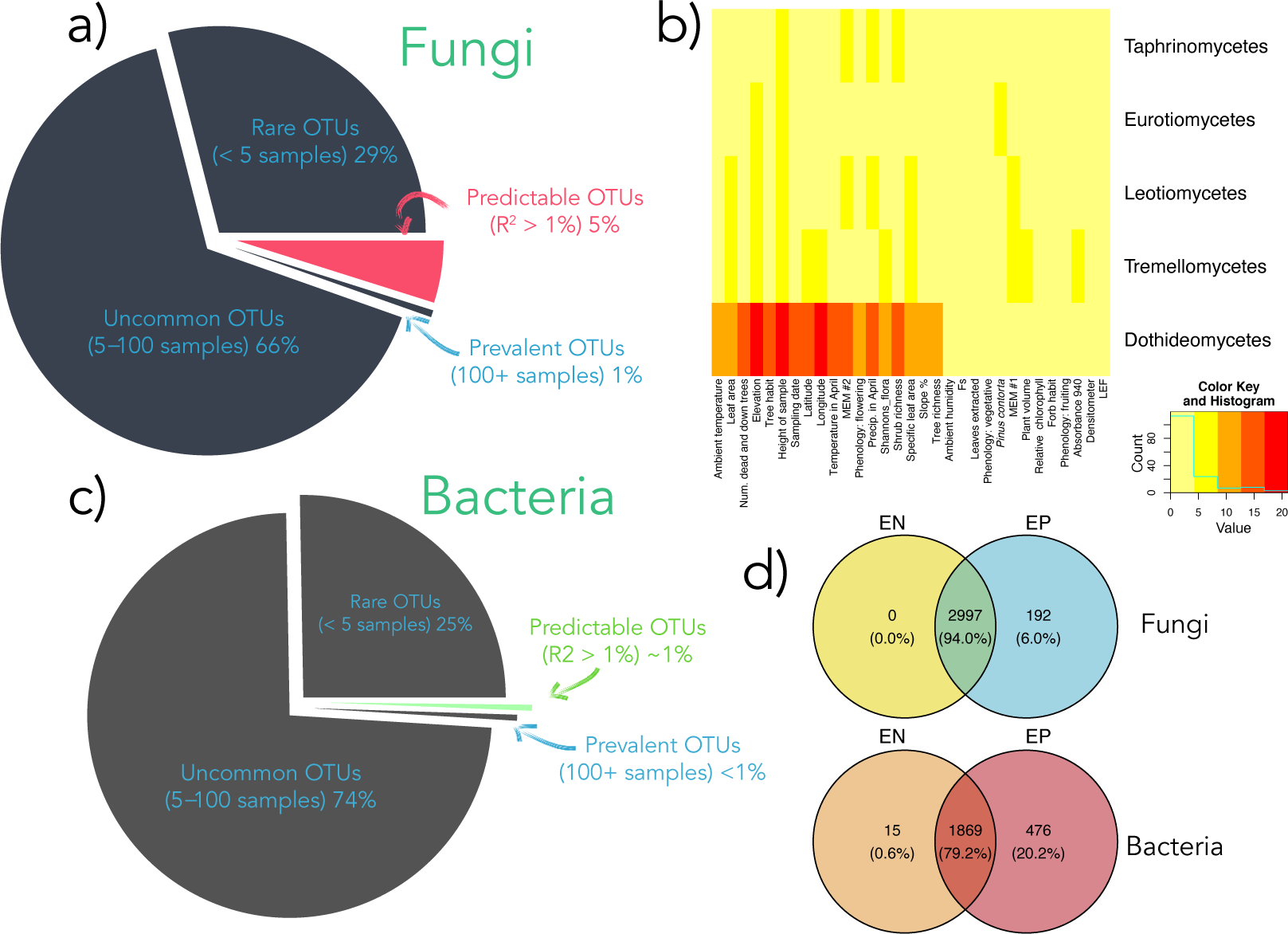
Most microbial taxa, be they fungal (a) or bacterial (c), had unpredictable abundances, in part because most taxa occurred in very few samples. Feature importances from models of fungal relative abundance are shown as a heatmap (b), with features that were more important for certain taxa shaded more darkly. Heatmaps were not generated for bacteria, because models for these taxa had poor performance. Features chosen were in the top ten most important for successful models (those models with an R sq over 1%) and were important for at least 20% of all successful models (thus those features that were important for isolated taxa are not shown here, for the sake of visualization). Only the fungal classes that each had at least five successful models are shown. For a similar figure, but for fungal occurrence models, see Fig. S15. Importantly, when models were rerun while including the abundances of certain co-occurring microbial taxa, model performance improved, though not beyond what is shown here. Microbial abundance features were typically some of the most influential in the model (see main text). d) Venn diagrams showing the number and proportion of microbial taxa that occurred as epiphytes (EP) or endophytes (EN) or both.

Ascomycota was by far the most abundant fungal phylum observed (Fig. 2) and Dothideomycetes the most abundant class. Proteobacteria and Actinobacteria were typically the most abundant bacterial phyla present, but there was more among-host heterogeneity in the abundances of bacterial phyla than for fungal phyla. For instance, Tenericutes were orders of magnitude more abundant in *Primula parryi* than in other plant taxa (Fig. 2). Hosts that were similar in terms of their fungal associates had similar bacterial assemblages, albeit the correlation was moderate (EN: r = 0.16, p < 0.01; EP: r = 0.18, p < 0.01). Patterns of host generalization shifted depending on microbial taxon and the plant compartment considered. For instance, for fungi, a similar degree of host generalization was observed for endophytes and epiphytes, but, in contrast, bacterial epiphytes were more generalized than bacterial endophytes (Fig. S7).

Ordinations and associated PERMANOVA analyses suggested greater divergence among samples in fungal assemblages compared to bacterial assemblages (Figs.S8– S11) and that the two groups of microbes responded to different dimensions of the phyllosphere habitat. Specifically, fungal samples weakly clustered by plant compartment (PERMANOVA; *R*^2^= 0.01, p < 0.01) and host life history (*R*^2^= 0.02, p < 0.01), and more strongly clustered by nominal taxon (*R*^2^= 0.17, p < 0.01). In comparison, plant compartment influenced bacteria much more than fungi (*R*^2^= 0.13, p < 0.01) as did host taxon (*R*^2^= 0.30, p < 0.01), but host life history was a poor predictor of assemblage dissimilarity (*R*^2^= 0.03, p < 0.01). The homogeneity of variances assumption of PERMANOVA was violated for these analyses, which can lead to less accurate p value determination.

### At the landscape level, most foliar microbial taxa have unpredictable abundances and occurrences

We attempted to predict the abundance and occurrence of microbial taxa that occurred within the leaves of Wyoming plants as a function of plant traits and abiotic conditions. We considered samples from 59 plant taxa growing at 20 sites spanning the mountains of Wyoming. During modeling, we considered microbial taxa that were present in ∼10% or more of our samples; very rare taxa were not considered because they were not observed enough to build informative models. Focal taxa were generalists, often occurring in 40 or more hosts (Fig. S7).

We used the random forest algorithm for predictive modeling and measured model performance using a nested resampling procedure. Models were deemed successful if they had an *R*^2^ of greater than 1%. Using this generous threshold, we could predict the absolute abundances of 57 fungal taxa (out of 172) and 4 bacterial taxa (out of 25). The median *R*^2^ for both fungi and bacteria was 0%. To check that these poor results were not due to model misspecification, data that were predictive of the response were simulated and included in a test model. Addition of simulated data dramatically increased *R*^2^, as expected.

When the abundances of co-occurring microbial taxa were included in models as features, model performance improved, with 114 fungal models and 18 bacterial models having positive *R*^2^ values. The median improvement in *R*^2^ for models that included other microbes as features was 0.07 for fungi and 0.08 for bacteria.

Relative abundances of fungi (Hellinger standardized count data) were easier to predict and models output *R*^2^ ≥ 0.01 for 155 taxa (the median *R*^2^ was 0.11 and the maximum was 0.40). But bacterial relative abundances were no easier to predict than their absolute abundances—only 12 taxa were predictable (median *R*^2^ = 0.03, max = 0.21). More abundant taxa were easier to predict, as expected; *R*^2^ was positively correlated with the relative abundance of fungi (r = 0.24, p < 0.01) but not significantly for bacteria (r = 0.32, p = 0.33; Fig. S12). Surprisingly, *R*^2^ was weakly negatively correlated with absolute abundances for both bacteria and fungi (Fig. S13).

Microbial presence within a sample (occupancy) was not easier to predict than abundance. To measure the performance of occupancy models, we used the Matthew’s correlation coefficient (MCC; Matthews 1975). The MCC takes into account true and false positives and negatives, deals well with imbalanced data, and ranges from negative one to one. Values of zero denote a model that performs no better than a guess. The median MCC for bacteria was zero and for fungi was 0.02. MCC for fungi was over 0.2 for 39 fungi and 1 bacterial taxon (an MCC of 0.2 represents modest predictive performance). Models typically did well when predicting absences, which was expected given that most microbes were infrequently observed. Prediction of presences was much more challenging—the median percentage of correctly predicted occurrences was 0% for bacteria and 2% for fungi. MCC was correlated with prevalence for both bacteria (R = 0.44, p = 0.03; Fig. S14) and fungi (R =0.29, p < 0.001).

For those fungal taxa that had model *R*^2^ of ≥ 1% or an MCC ≥ 0.2 we examined which model features (predictor variables) were the most important. To do this, we counted the number of times a feature was in the top 10 most important for 20% or more of models (Fig. 3, S15). Feature importance was determined via permuting each feature, rerunning the model, and calculating the decline in *R*^2^ or MCC. We did not consider models for bacterial taxa, due to their poor performance. The importance of specific features differed among taxonomic groups and data sets. For instance, the relative abundances of fungi tended to be influenced more by sampling height, elevation, leaf area, leaf density (SLA), and date and less by aspects of leaf productivity, such as linear electron flow and Fs, than were fungal absolute abundances and occupancy (Tables S2, S1, S3). There was no obvious pattern of the relative importance of abiotic versus biotic variables. Instead, most of the features we considered were identified as important for at least some taxa and not a single feature was in the top 10 most important for all successful models.

Despite these idiosyncrasies, we uncovered similarities among the best models for Ascomycetes (12 models of relative abundances that all had *R*^2^ *>* 0.25). For all of these models, elevation and shrub richness were important (Table S4), and sampling date, temperature, and latitude were often so. Even among these top models, when considering features that were often important, the relationship between feature variation and relative abundance shifted among taxa (Fig. S16). For instance, some taxa responded sharply negatively to increased shrub richness initially and then leveled off as shrub richness increased, others did not respond much at all to changes in shrub richness, and still other taxa were positively associated with increased shrub richness. Similar patterns were observed for other important features and non-linear relationships between features and response variables were common.

Because host taxa were included as one-hot encoded features in our model, their aggregate importance was not represented via the post-hoc perturbation approach we used to estimate feature importance. Therefore, modeling was repeated without host taxon and the decline in performance recorded. For absolute abundance models of fungi, *R*^2^ declined from 0.01-0.09 with an median decline of 0.01. Models for twenty fungal taxa had *R*^2^ values that dropped below 0.01 when host nominal taxon was removed. This pattern was mirrored for relative abundance data. For bacteria, the median and mean *R*^2^ was unaffected by the inclusion of nominal taxon in models.

When absolute abundance models included the abundances of co-occurring taxa as features, those taxa tended to hold great influence over model performance. Indeed, these features were selected as the most important in models, typically exceeding host traits and abiotic conditions. Influential taxa spanned multiple phyla (Tables S5 & S6).

### Intraspecific trait variation does a poor job of predicting microbial abundances

To better quantify the associations between microbial abundances and intraspecific variation in plant traits, we modeled microbiome variation within specific hosts. We did this because plant trait variation was confounded with nominal taxon. We only considered those combinations of host and microbe that were sampled at three or more locations, that were in 30% or more of hosts sampled, and that were present in at least 30 samples. 110 combinations of fungi and 15 geographically widespread host plants met these criteria, but only 9 combinations of host and bacterial taxa did. For absolute abundance data, 16 of the fungi-host combinations had *R*^2^ values between 0.01–0.08 but none of the models for the bacterial combinations met with any success. Phenology, relative chlorophyll, and compartment were influential features (Table S7). As we found in our models across hosts, model performance was greatest for relative abundance data—66 of the fungi-host combinations and 4 of the bacteria-host combinations had positive *R*^2^ values. For these models, SPAD 420, relative chlorophyll, compartment, phenology, and canopy cover were influential features (Table S8).

We repeated this analysis using occupancy data and noticed some improvement in model performance compared to the landscape-wide analysis: 81 combinations of fungal taxon and host and 3 bacterial combinations were modestly predictable (MCC ≥ 0.2). Median MCC for both bacteria-host combinations and fungi-host combinations was 0.12, as before models were challenged to predict microbial presences, not absences. About 4% of bacterial presences were correctly predicted and 13% of fungal presences were predicted (these are estimates of the median proportion predicted correctly across taxa). Compartment (either EP or EN), SPAD 420, tree richness, phenology, relative chlorophyll, and gh were the most important features for fungal models (Fig. S9; bacterial models were not considered because so few were successful).

### Microbial diversity was predictable but richness was not

Shannon’s diversity and estimated richness tended to be higher for fungi than bacteria (Fig. S17–S20). Patterns in richness and diversity among samples were few, though epiphyte diversity tended to be slightly *lower* than endophyte diversity for bacteria whereas the opposite was true for fungi (in many cases these comparisons were not significant, though the pattern is suggestive; Tables S10 & S11). No growth habit (i.e., tree, forb, shrub, graminoid) was much more diverse than any other, though we did find drastic differences among host taxa—particularly for fungi (Fig. S18, S19). The richest plant taxa in terms of fungi (but not bacteria) were trees and shrubs (this pattern held for both compartments; Tables S14, S15).

Neither fungal nor bacterial richness was predictable (*R*^2^ was near zero for both models). In contrast, models of Shannon’s diversity were somewhat successful (fungi *R*^2^ = 0.13; bacteria: *R*^2^ = 0.08). Important features for predicting fungal diversity included compartment (EP vs. EN), phenology, *Juniperus communis* (a widespread host plant taxon), and elevation (Fig. S21). Important features for bacterial diversity were different than those for fungal diversity and included densiometer measurements (canopy cover), date collected, and *Primula parryi* (an unusual plant taxon; Fig. S22). Diversity of both fungi and bacteria varied among host nominal taxa, and, indeed, certain taxa were strong predictors of shifts in diversity (Fig. S18, S19). When host taxon was omitted from models their performance declined slightly (dropping from 13% to 12% median explained variance for fungi and from 8% to 5% for bacteria). Inclusion of sampling site in our models, which was a proxy for soil variation and other unmeasured abiotic phenomena occurring at the sub-*km*^2^ spatial scale did not improve predictions.

We found drastic differences in the ability to predict epiphyte versus endophyte data. For example, a model of bacterial endophyte diversity had an *R*^2^ of ∼0.17 (Fig. S25), but the model of bacterial epiphyte diversity had near zero explained variance. Fungal endophyte and epiphyte diversity were both predictable (10–14% explained variance), but feature importance was shuffled between models (Figs. S23, S24). Elevation and latitude were the most important features for epiphytes whereas the number of leaves extracted and two host taxa (*Juniperus communis* and *Antennaria media*) were the most important for fungi.

## Discussion

Prediction of natural variation in phyllosphere microbial assemblages appears to be quite challenging. We found that only a few percent of foliar microbial taxa have predictable abundances or occurrences, even when using powerful machine learning to analyze a large dataset of over 1000 individual plants, 59 plant taxa, and 43 covariates that together characterized the foliar habitat. Our results are primarily due to the rarity of the majority of microbial life—most taxa were represented by a few reads in a few samples, making these taxa impractical to model. But predictive ability was low even for those few microbial taxa that were quite abundant and prevalent.

This has important implications for predictive ecology and biogeography—specifically, that we should not expect predictive modeling to be successful for most organisms, given that they are so infrequently observed. This observation almost seems trivial. However, we suggest that the literature does not reflect this inherent limitation within the data. Most publications assume that the biogeography of foliar microbiomes (and other microbiomes) is largely defined through deterministic, and thus predictable, causes. For example, our expectation for this project was that a large proportion of the variation in foliar microbiomes would be ascribable to the numerous aspects of the foliar habitat that we measured. However, the nature of ecological power laws (and sampling logistics) suggests that hard limits to prediction exist for most taxa, which should temper our expectations for the strength of associations between microbiome composition and habitat variation. Indeed, we are aware of no publications that demonstrate more than a moderate association between foliar microbiome composition and habitat variation. By moderate association, we refer to such results as estimated *R*^2^ values from PERMANOVA analysis of dissimilarity matrices less than 0.4. We suggest that stronger associations may not be observable, given that most taxa are rare and, as we show here, even more abundant taxa are often very challenging to quantitatively predict. We suggest that the limits to prediction may be estimated through mathematical means, given estimation of the parameters for the distributions underlying the data. Such a theory driven approach could precisely bound expectations for microbial biogeography.

While it seems that precise predictive modeling of rare microbial taxa is out of reach, at least for now, we do not suggest that these myriad taxa are unimportant. Indeed, a growing body of work demonstrates that low-biomass microbes can perform critical ecological functions (Jousset et al. 2017; Kalenitchenko et al. 2018). Moreover, it is unclear if rare taxa bloom into abundance given suitable conditions (Shade et al. 2014), just as the seed bank of annual plants germinates following spring rainfall. Thus, some of the taxa we deemed rare in our snapshot-style study could be more abundant at other times of year or immediately following shifts in abiotic conditions or host phenology. This speculation suggests that incorporating data from intensive temporal sampling, particularly including sampling across host phenology, could improve predictive modeling of microbiome composition, though the aforementioned limitations likely will still exist.

### Fungi and bacteria differed in abundances and predictability

When summarizing the results from our models, we found several interesting patterns in predictive performance. Most notably, fungi tended to be easier to predict than bacteria, regardless of data set and model choice. This may be due to the much better sequencing results we obtained for fungi than bacteria. Plant chloroplast sequences dominated the 16S data (Fig. S6), a problem noted elsewhere (Maignien et al. 2014), and that suggests that plant chloroplasts are much more common than bacterial endophytes within most leaves. Karasov et al. (2019) reported similar findings. They used shotgun metagenomics, controlled infections, and qPCR to quantify the ratio of host to bacterial genomes as 1–2.5 within the leaves of wild *Arabidopsis thaliana* and that foliar microbes may not become extremely abundant except during infection. Given that *A. thaliana* is an annual forb with small leaves, it seems reasonable that a longer-lived plant with larger leaves could accrue more microbial taxa. Still, when taken together, our results and those from Karasov et al. (2019) suggest that the ratio of bacteria to host cells within the leaves of healthy plants is probably low, though much more quantitatively rigorous work is needed. Unfortunately, we could not compare the ratio of fungal to plant reads because we recovered few of the latter due to the specificity of our ITS primers.

### Plant traits were only weakly associated with microbiome variation

We uncovered associations between many plant traits and specific abundant microbial taxa, but the associations were typically weak and idiosyncratic. Interestingly, we included 24 plant traits in models and yet nominal plant taxon remained modestly important. This suggests a weak effect of unmeasured host traits on microbial assemblages, possibly directly, because those traits could influence the ability for a microbe to encounter or live within a plant, and indirectly, since plant traits determine, at least in part, where a plant grows, and which other microbes might be present in the phyllosphere. While nominal taxon was an important feature in models, it is worth reiterating that removing this feature typically led to only a modest drop in *R*^2^. Thus, our results suggest that trait variation alone is unlikely to explain a large fraction of the variation in foliar microbiome assemblages—because nominal taxon is a proxy for all unmeasured host phenotypic and genetic variation of relevance to the foliar microbiome.

Perhaps many easily observable plant traits are too far removed from the spatial scale of relevance for microbes to be strongly predictive of microbiome variation. Indeed, for many microbes a single leaf is the equivalent of the whole state of Wyoming (the study area) for a human. Those traits that were important were proxies for microhabitat variation in leaves, including leaf productivity (e.g., relative chlorophyll, SPAD 420), specific leaf area, area of leaf sampled, and phenology (Tables S2–S9). For logistical reasons, we did not measure elemental concentrations in leaves or various phytochemicals, though previous work has shown weak associations between these traits and microbial assemblage composition and they could be useful to include in future studies (González-Teuber et al. 2020; Kembel 2009; Kembel et al. 2014).

The lack of strong association between variation in plant traits and the microbiome is puzzling because numerous experiments have demonstrated, beyond all doubt, that many microbes can mediate plant trait expression (Friesen et al. 2011; Hawkes et al. 2021)—in some cases, dramatically (e.g., Doty 2011; Doty 2008). This disparity could arise from the frequency with which plants encounter microbes in the wild. Indeed, from before the day a seed germinates to after its last leaf senesces, plants are continually interacting with microbes. Thus, without experimental removal or inoculation of specific taxa, the effects of microbes on many, easily-measured plant traits may be very challenging to observe (and thus predict), since host plants have never escaped from their microbial influencers. We note that these influencers need not be abundant within plants. For example, we have previously demonstrated that low biomass foliar microbes can affect plant traits (Harrison et al. 2021b), and, in that study, the low biomass microbes were so infrequent within the sequencing data as to preclude predictive modeling.

### Perhaps unmeasured ecology is behind poor model performance—including microbe-microbe interactions

When we repeated modeling while including the abundances of co-occurring microbes as features, model performance tended to increase. This was not solely due to the additional complexity of models, because the addition of randomized versions of these features did not increase model performance. This suggests that microbe-microbe interactions, be they antagonistic or mutualistic, direct or indirect, are important determinants of phyllosphere microbiomes. Recent work supports this hypothesis (Hassani et al. 2018; Harrison et al. 2021b), though mechanisms remain uncharacterized.

During modeling we focused on bottom-up forces, and including top-down pressures should improve model performance—after all, predicting the occurrence or abundance of herbivores (e.g., ungulates, insects) would likely fail without considering predators. Indeed, Morella et al. (2018) recently demonstrated, in a manipulative setting, that phages can shape bacterial assemblage composition within plant leaves.

We consider the possibility that the abundances and occurrences of many microbial taxa within individual leaves and plants is largely unknowable, as driven by the vagaries of dispersal and priority effects. Priority effects describe the situation whereby the sequence of encounters between microbial taxa and the leaf define the resulting assemblage. (Leopold and Busby 2020) recently demonstrated priority effects of fungi within the leaves of *Populus trichocarpa*, with order of immigration having effects on microbiome composition one month on.

Ecological drift is another stochastic force that shapes microbial assemblages via the aggregated influence of life history events (i.e., births and deaths; Vellend 2010). Ecological drift is notoriously difficult to study and so its influence on microbiome composition is debated (Zhou and Ning 2017), however theory from population genetics suggests that the influence of drift should decline with population size. Thus, we doubt that ecological drift plays a large role in our results, which focus on abundant microbial taxa, because these taxa should have high enough population densities when aggregated into samples that drift would have negligible influence assuming any differences among taxa in competitive ability or suitability to the habitat. However, ecological drift could be important for shaping the abundances of rarer taxa or ecologically equivalent taxa within an assemblage.

### Why are absolute abundances harder to predict than relative abundances?

For all of our analyses, we compared results using relative abundances and absolute abundances. Most microbial ecology studies rely on the former, because sequencing machines output relative abundance data (Tsilimigras and Fodor 2016). Incorporating internal standards (ISDs) during sequencing can allow absolute abundances to be approximated, though the approach is not without methodological challenges (Harrison et al. 2021c). We found that relative abundances were easier to predict than were absolute abundances, for both bacteria and fungi and regardless of scale, *except* when the abundances of co-occurring microbial taxa were included as features in models.

These results could be driven by biology: specifically, because the absolute abundances of microbes may shift in response to co-occurring microbial taxa or numerous unmeasured phenomena. Or, perhaps model *R*^2^ is higher for relative abundance models, because relative abundances are potentially more constrained then absolute abundances.

### The way forward

Over the past decade, phyllosphere microbial ecologists have explored the deterministic factors that shape foliar microbiomes. The numerous associations between plant traits, abiotic conditions, and microbial community composition that have been reported suggest that deterministic forces do matter. However, many of the associations described have been fairly weak, which led us to probe the limits of the approach through the use of a large dataset and an explicitly predictive paradigm. Our results were poor, partly because of the inherent challenge of predicting abundances and occurrences of rare taxa. However, even for the most well-observed microbial taxa in our survey, modeling was challenging. Our goal here was to explore if microbiome composition could be predicted, thus we did not build models at various taxonomic levels (e.g., at the phylum or genus level)—such models would be a potentially informative, next step.

The general failure of our models confirms that predictive phyllosphere ecology will require more than an encyclopedic characterization of the foliar habitat and its influence on microbial taxa. Instead, future work could profitably explore the influence of microbe-microbe interactions, top-down pressure (predation, e.g., by phages) and priority effects on foliar microbiome community composition. While such work will undoubtedly push the limits of prediction past what we present here, we suspect that the long-tailed rank-abundance curves typical of foliar microbiomes imply inherent constraints to predictive microbiology do exist.

## Acknowledgments

Funding was provided by the National Science Foundation EPSCoR grant 1655726. Computing was performed at the Advanced Research Computing Center, University of Wyoming, Laramie (https://doi.org/10.15786/M2FY47). Thanks to Ankita Arun Sawant, Jeff Waller, and William Herrick for laboratory assistance; Ernie Nelson and the Rocky Mountain Herbarium at the University of Wyoming for assistance identifying specimens; Gregg Randolph for guidance during library preparation; and W. John Calder and Matthew L. Forister for helpful advice.

## Author Contributions

JH and CAB designed the research; JH performed the research including field and lab work, data analysis and interpretation. JH and CAB both contributed to writing and editing the manuscript.

## Data availability

Voucher specimens of plant hosts are available at the Rocky Mountain Herbarium at the University of Wyoming. All scripts and plant trait data are available at the following DOI: 10.5281/zenodo.6112672. Raw and processed sequence data are available at: DOI for data is forthcoming and will be available prior to publication.

## Supplementary Material

### Field sampling details

Sampling locations were chosen to maximize geographical coverage within the state, facilitate reasonable access, and ensure specific vegetation assemblages were sampled. Climate data for each site from the month prior to sampling were downloaded from PRISM (PRISM Climate Group, Oregon State University, http://prism.oregonstate.edu). To control for the effect of aspect, all plots were located on west-facing slopes.

Plant taxa were chosen to ensure that approximately 80% or more of the biomass at each site was sampled (as visually estimated) and to maximize the phylogenetic and phenotypic breadth surveyed. Samples were collected using flame sterilized forceps, bagged, stored on ice, and frozen within approximately 12 hours of collection using a battery-powered freezer. At least three leaflets were sampled from most individuals, except for plant taxa with very large leaves, where a single leaf was removed for DNA extraction. Because of the necessity of sampling different numbers of leaves among taxa, the mass of leaf material from which DNA was extracted was included in models.

Within each plot, two transect tapes were placed along the cardinal axes with the intersection of the two tapes in the center of the plot. These tapes served as a Cartesian coordinate system. For focal plant taxa, random coordinates were generated and the nearest individual to those coordinates was sampled. If a plant taxon only occurred in a portion of the plot, individuals were sampled haphazardly. To characterize overall vegetation composition in each plot, all plant taxa within four randomly located 1m quadrats were assigned to one of the following cover classes: ≤0.1%, 0.1–5%, 5–25%, 26–50%, 51–75%, 76–95%. Data from each quadrat were aggregated by site and floral diversity (exponentiated Shannon’s) was calculated from the upper values of the ranges for each cover class.

For each focal plant, phenology, height, width at widest point, the width perpendicular to this point, and the product of these measurements, were collected. For trees, trunk diameter at ∼1.5m above the ground was also measured. A leaf proximal to those sampled for DNA extraction was removed for measurement of leaf area, specific leaf area (also known as density), leaf toughness (using a Chatillon 516-0500 force gauge), and water drop retention ability. The leaf was chosen such that it was the approximately equal in size and shape to those chosen for DNA extraction. It was not practical to perform DNA extraction on the same leaf used for these measurements due to contamination concerns. Water drop retention ability was the angle (in degrees) at which a droplet of water (4–5*µ*l) placed on the leaf surface began to move. Leaves were dried, digitally imaged, and leaf area was measured using imageJ (Schneider et al. 2012). The same leaves were weighed and specific leaf area (SLA) calculated. Additionally, for each focal individual, the Photosynq multispeQ (East Lansing Michigan, USA) portable flurometer and spectrophotomer (Kuhlgert et al. 2016) was used to measure relative chlorophyll content, leaf surface temperature, photosynthetically active solar radiation (PAR) in the vicinity of the leaf, various metrics of light absorbance and photosynthesis by the leaf, and ambient weather conditions (e.g., humidity; Table 1). In addition, to characterize and account for spatial relatedness among sampling locations during modeling, distance-based Moran’s eigenvector maps (MEM) were calculated at the site level using the dbmem function of the adespatial R package (Dray et al. 2016). The geodesic distance between sites was used generate a distance matrix, which was decomposed using dbmem, and the top two MEMs included in all models.

### Epiphyte removal

We removed epiphytes via washing leaves using detergent and water (see main text). Evidence for the success of this procedure includes the number of taxa that were observed within epiphyte samples that did not appear in endophyte samples (Fig. 3). Additionally, a subset of leaves (n = 84) were washed twice and the solution from the second rinse sequenced. On average, slightly fewer sequencing reads were obtained from secondary rinses then primary rinses and ordinations showed that rewashed samples were different than washed samples for bacteria (Fig. S1 –S4). Fungal ordinations showed a general overlap of rewashed samples with both epiphyte and endophyte samples, suggesting that fungi did not differ as much among compartments as did bacteria, possibly because fungi were harder to wash off or because they grew both within and without leaves (see main text).

### Library preparation details

DNA was extracted using Qiagen DNEasy plant kits (plate format). Libraries were prepared using a two-step procedure described in Harrison et al. (2021a). For bacteria, the 16S (V4) locus was amplified using the 515-806 primer pair (Wang and Qian 2009), while for fungi the ITS1 locus was amplified using the ITS1f-ITS2 primer pair (White et al. 1990). We used shortened variants of the molecules presented in (Tourlousse et al. 2017) as internal standards (see Harrison et al. 2021a, for details). To account for cross-contamination, short synthetic sequences were added to each well of the 96-well plates used during library preparation. By tracking these known sequences, we were able to identify instances of cross-contamination and remove those samples from analysis (see Harrison et al. 2021a). We added an equimolar amount (∼0.18 pg, which translates into ∼209–215 million molecules depending on the ISD; we had an ISD for each locus) of a synthetic DNA internal standard (ISD) to each sample prior to PCR. ISDs were inspired by (Tourlousse et al. 2017) and consisted of synthetic DNA (matching no known organisms) that was placed in between primer sequences.

Libraries were created using duplicate PCR reactions consisting of 6 µl of 0.01 pg µL*^−^*^1^ coligos and 0.03 pg µL*^−^*^1^ of the ISD and 30 ul of template. Template was normalized to 10 ng µL*^−^*^1^. 6 µl each of 0.25 µm forward and reverse primers, 0.3 µl Kapa HiFi HotStart DNA polymerase (New England BioLabs, Ipswich, MA, U.S.A.), 0.45 µl 10 M dNTPs, 3 µl 5x KAPA HiFi HotStart PCR buffer (New England BioLabs, Ipswich, MA, U.S.A.), 3.25 µl of water, and 2 µl of template were mixed and used for the first round of PCR. The PCR recipe was: denaturation at 95°C for 3 min, followed by 15 cycles of 98°C for 30 s, 62°C for 30 s, and 72°C for 30 s, and a final 5 min extension at 72°C. Amplicons were cleaned using Axygen AxyPrep MagBead (24 µl; Corning; Glendale, Arizona, U.S.A). Reaction volume for the second round of PCR was 15 µl. 0.5 µl each of 10 µm flowcell primers, 0.3 µl Kapa HiFi HotStart DNA polymerase, 0.45 µl 10 M dNTPs, 3 µl 5x KAPA HiFi HotStart PCR buffer, 0.75 µl of water, and 10 µl of cleaned product from the first round of PCR were mixed for the second round of PCR. Denaturation at 95°C for 3 min, was followed by 19 cycles of 98°C for 30 s, 55°C for 30 s, and 72°C for 30 s, and a final 5 min extension at 72°C. Products were again cleaned using magnetic beads, normalized, pooled, and sent off for sequencing by psomagen, Inc. (Rockville, MA, U.S.A.). Library success was determined via qPCR and by using a Bio-Analyzer (Agilent, Santa Clara, CA, U.S.A). Aside from sequencing, all laboratory work was conducted at the Genome Technologies Laboratory at the University of Wyoming (Laramie, WY, U.S.A.).

Since each PCR replicate was tagged with a unique molecular identifier, we were able to compare replicates. Sequence counts for PCR replicates were highly correlated and so were combined in downstream analyses.

### 0.1 Bioinformatics

Paired-end sequences were merged using vsearch v2.9.0 (Rognes et al. 2016). Some ITS reads did not overlap and thus were trimmed to a fixed length and concatenated. Reads with more than a single possible error were removed. Exact sequence variants (ESVs) were determined via vsearch and then these variants clustered by 97% similarity. We decided to combine ESVs in this way because we recovered many tens of thousands of ESVs and treating each of these ESVs individually was unwieldy. The large number of ESVs we recovered was at least partially due to the extreme sequence depth possible with the NovaSeq and the patterned flow cell of this machine may also have contributed (Singer et al. 2019). Taxonomic hypotheses were generated using the SINTAX algorithm (Edgar 2016) and the UNITE (for ITS; v7.2; Community 2017) and Greengenes (for 16S; v13.5; DeSantis et al. 2006) databases.

### Modeling estimated proportion data

Division by the ISD cannot estimate the probability that a zero is biological or due to low sequencing depth; therefore, a zero before normalization is a zero afterwards. To circumvent this issue we also attempted to model all data using a hierarchical Bayesian model that shares information among replicates within a sampling group (in our case, a sampling group was a host taxon and compartment at a site) to estimate the probability of observing a taxon in each sample; overall results were similar to those presented here for models of individual taxon abundances and diversity (see the Supplemental Material; Harrison et al. 2020b; Harrison et al. 2020a).

We also modeled count data using a hierarchical Bayesian approach implemented via the CNVRG R package (v. 1.0.0; Harrison et al. 2020b). This method estimates taxon proportions within a sample as parameters of a multinomial distribution with a Dirichlet prior, which is used to share information among samples within treatment groups (Harrison et al. 2020a). Treatment groups were the combination of host taxon and compartment at individual sites. For example, we shared information among the ten EN samples from a particular plant taxon at a location. The benefit of the CNVRG approach is the information sharing among samples within a treatment group and the ability to pass uncertainty in microbial abundances to downstream analyses. Our rationale was that sharing of information could be useful for estimating the probability of observing rare taxa when there was extreme variation in sequencing depth among samples. This style of modeling, since it relies on proportional data, avoids the issues imposed by rarefaction.

When using proportional abundances extracted from our hierarchical Bayesian approach in our random forest, model performance was generally poor for fungi, with *R*^2^ exceeding 1% for only 8 fungi. All 23 bacteria had model *R*^2^ of between 5–9%. The similarity among bacteria likely is due to the constraints imposed by prior structure of the hierarchical model, given the relatively low read count for many bacterial taxa in many samples.

### 0.2 Machine learning details

We used a rigorous approach to model fitting, tuning, and performance determination that was reliant upon the mlr3 R package (v 0.12.0; Lang et al. 2019). mlr3 provides an interface to assist with the considerable programmatic bookkeeping needed to run many thousands of models using different parameters and datasets. Our general approach was to split the data into many testing and training datasets, then train the model on those data, tune the hyper-parameters of the model and then assess model performance. Imputation was done within each sub-dataset, as was hyper-parameter tuning, thus avoiding information from the training data leaking into the testing data.

Feature engineering included conversion of categorical covariate data to numeric data via one-hot encoding, imputation of missing data, and scaling and centering data (conversion to z scores). Features were built to stratify data during splitting into testing and training sets to ensure similarity between both datasets. For example, a feature describing if a focal taxon was above its median abundance in a sample was used to ensure that neither the testing nor the training data were under-representative of the focal taxon’s variation in abundance. Plant compartment (either EP or EN) was also used during stratification, because *a priori* analysis suggested it was an important determinant of microbiome composition.

Correlation of features was examined and one of a group of highly correlated features were included in models (e.g., elevation was included but not atmospheric pressure; Fig. S5). The exception to this were several features that were correlated with elevation, such as mean temperature and tree and shrub species richness within sampling plots. These features were retained since they varied among lower elevation sites and because they were of general interest to us.

Hyper-parameter tuning was performed using the “AutoTuner” function of mlr3 using the hyperband tuning algorithm (Li et al. 2017), which adaptively allocates compute resources to better explore higher performing portions of parameter space. Hyper-parameters considered during tuning were the number of trees in the ensemble (50–400), the fraction of the samples used for each tree (0.01–0.3), the minimum node size (1–25; meaning how many samples are retained in the “leaves” of the trees), and the splitting rule used for designating splits in decision trees. Both the “extratrees” and “variance” split rules were considered. The former institutes the “extremely randomized trees” approach of Geurts et al. (2006), where splits in the trees are assigned randomly. The “variance” splitting rule determines splits such that variance in the response is minimized within each group delineated by the split and trees are built using subsets of the data. A nested resampling approach (using mlr3; three outer splits and four inner splits) was used to determine an unbiased estimate of model performance.

### 0.3 Supplemental results and discussion

Our poor results likely *overestimate* our ability to predict microbiome composition. This is because much of our sequencing data was composed of extremely rare genetic material that was removed during the bioinformatics prior to modeling. Between both loci surveyed, over 500,000 unique sequences were obtained, most of which were discarded during filtering (a process referred to as “denoising”, a standard step to remove possible technically-derived variants). While it is reasonable to assume a subset of these unique sequences were artifacts of PCR, it seems certain that some were biologically-derived. Unfortunately, current technology precludes accurate provenance determination of infrequent sequences, consequently the current standard is to discard these data.

Because we used the Novaseq platform, which improves upon older Illumina technology through the use of a patterned flow cell that can better distinguish between clusters during sequencing-by-synthesis, we may have obtained sequences from more rare taxa than is typical (a possibility suggested in Singer et al. 2019). However, the rank abundance curves that we observed for our data generally match those reported in the literature—with few abundant taxa and many rare taxa (Figs. S31 & S32). Indeed, this pattern is ubiquitous in ecology (Hubbell 2001), though perhaps microbial assemblages tend to have a longer tail of rare taxa than do macrobial assemblages (Locey and Lennon 2016).

There was an interesting disparity in overall abundance between bacteria and fungi when moving between plant compartments. Bacteria tended to be more abundant on the surfaces of leaves compared to their interiors, but the opposite was true for fungi (an inference that was made possible through the use of an internal standard). This result affords ample opportunity for speculation regarding mechanism. Perhaps fungi can better survive the plant’s immune system than bacteria or perhaps the result is because some fungi possess better tools, such as appressoria, to make their way into leaves. More simply, perhaps fungi have a harder time growing on the surface of the leaf than do bacteria. We suggest that culturing studies that simulate growth inside versus outside of a leaf (e.g., by varying ultraviolet light exposure, diel cycles in abiotic conditions, and nutrient availability) could provide insight into our results.

We also found that leaf area and Moran’s eigenvector maps were influential features in our models, as might be expected if dispersal was important (though, of course, these same features are confounded with potentially important variation in the foliar habitat), yet these features collectively explained only a small proportion of microbiome variation. If dispersal and priority effects are key to understanding patterns of variation among microbial taxa then this adds impetus to the study of dispersal rates and distances, taxon-specific propagule counts, and spore persistence among microbes, as without this basic natural history knowledge parameterizing theory will not be possible (i.e., island biogeographic theory).

We do not think that problems with the ISD itself caused the failure of our absolute abundance models because models were taxon-specific and samples that did not include the ISD (for instance, due to poor sequencing) were omitted from analysis. That is, variation in a single microbial taxon was modeled after dividing the sequence counts for that taxon by the counts for the ISD—thus, stated colloquially, models compared apples to apples. This approach was necessary because inter-taxon comparisons are not reliable due to copy-number variation (CNV) and likely differences among taxa in ISD commutability, which describes the similarity in behaviour between the ISD and a particular taxon during extraction and PCR (Harrison et al. 2021c). CNV could vary among samples for a single microbial taxon, but this should have influenced both relative and absolute abundances similarly and thus seems unlikely to be the primary reason absolute abundance models performed so badly when co-occurring microbes were not included as features. We note that without an ISD we would not have trusted the results from models that included co-occurring microbes as features, because of the spectre of compositionality.

---

**Table S1:**
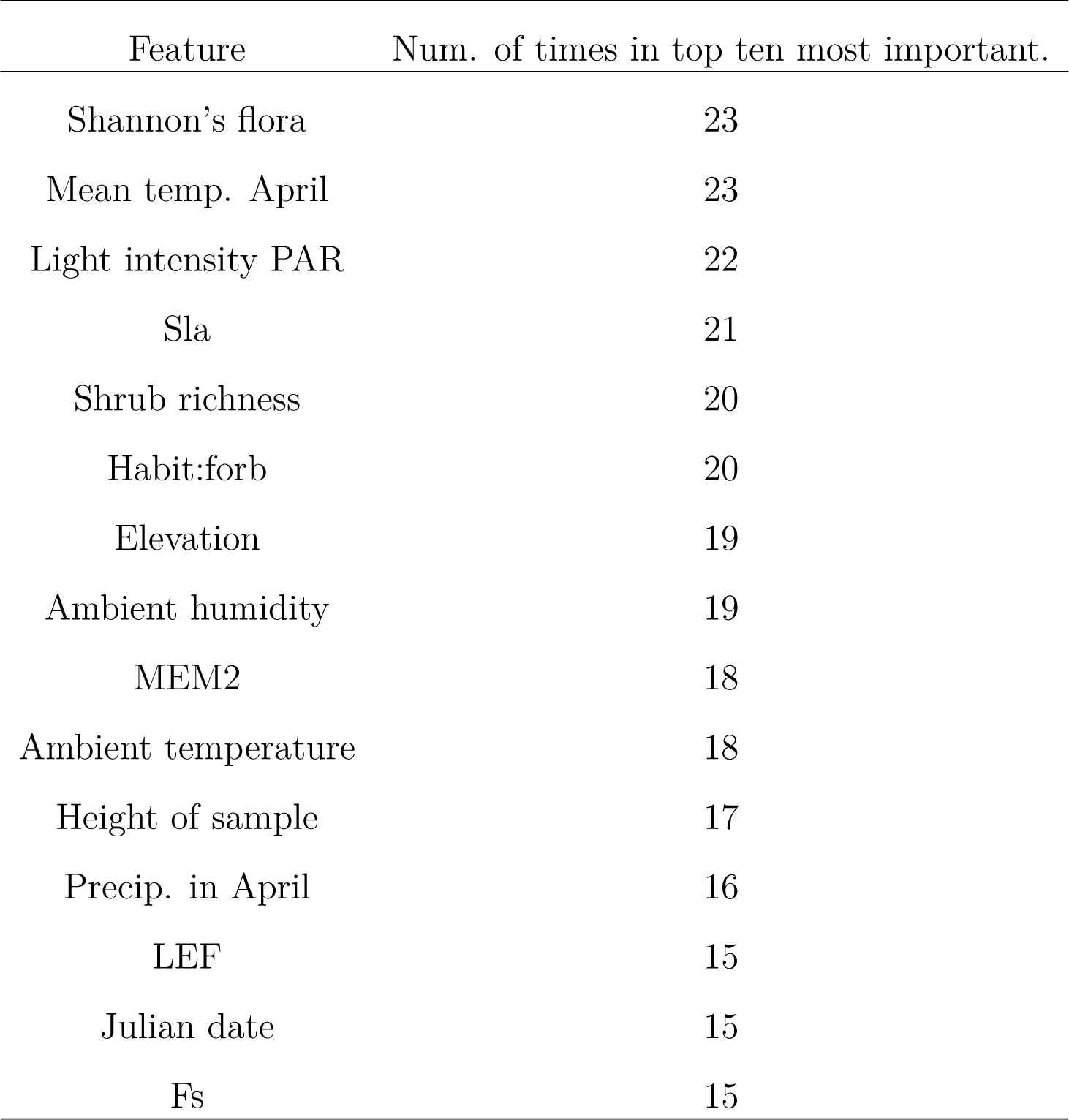
Number of times a feature was in the top ten most important for models of fungal abundance (ISD standardized) at the landscape level and which had an R^2^ > 0.01. Only the top fifteen most important features are shown. 57 taxa out of 172 that met our modeling requirements were predictable.

| Feature | Num. of times in top ten most important. |
| --- | --- |
| Shannon's flora | 23 |
| Mean temp. April | 23 |
| Light intensity PAR | 22 |
| Sla | 21 |
| Shrub richness | 20 |
| Habit:forb | 20 |
| Elevation | 19 |
| Ambient humidity | 19 |
| MEM2 | 18 |
| Ambient temperature | 18 |
| Height of sample | 17 |
| Precip. in April | 16 |
| LEF | 15 |
| Julian date | 15 |
| Fs | 15 |

**Table S2:** Number of times a feature was in the top ten most important for models of fungal *relative abundance* (Hellinger standardized) at the landscape level and which had an *R*^2^ *>* 0.01. Only the fifteen most important features are shown. 155 taxa out of 172 that met our modeling requirements were predictable.

| Feature | Num. of times in top ten most important. |
| --- | --- |
| Height of sample | 94 |
| Elevation | 81 |
| Leaf area | 73 |
| Sla | 72 |
| Julian date | 63 |
| Longitude | 60 |
| Latitude | 55 |
| Precip. in April | 53 |
| Habit:tree | 53 |
| Shannon's flora | 52 |
| Shrub richness | 51 |
| MEM2 | 51 |
| Mean temp. April | 50 |
| Plant volume | 49 |
| Ambient temperature | 45 |

**Table S3:** Number of times a feature was in the top ten most important for models of fungal occurrence with an MCC of 0.2 or greater. Only the fifteen most important features are shown. 39 taxa out of 172 that met our modeling requirements were predictable.

| Feature | Num. of times in top ten most important. |
| --- | --- |
| Height of sample | 41 |
| Leaf area | 40 |
| Sla | 37 |
| Elevation | 35 |
| Plant volume | 25 |
| Mean temp. April | 25 |
| Longitude | 25 |
| MEM2 | 20 |
| Latitude | 19 |
| Julian date | 19 |
| Shrub richness | 16 |
| Ambient humidity | 16 |
| Precip. in April | 15 |
| Dead and down | 15 |
| Shannon's flora | 14 |

**Table S4:** Features ranked by the number of times they were in the top ten most influential for only those models of relative abundances that had an *R*^2^ *>* 0.25. Models for fifteen fungal taxa had such high *R*^2^ values and 12 of those taxa were hypothesized to be within Ascomycota. The other three taxa were not placed within a phylum via a SINTAX query of the UNITE database (see main text). Only the top 20 features are shown. Notably, no single fungal taxon was predictable at an *R*^2^ *>* 0.25 when using ISD transformed data (absolute abundance) without including co-occurring microbes.

| Feature | Num. of times in top ten most important. |
| --- | --- |
| Shannon's flora | 3 |
| Ambient temperature | 4 |
| Taxon: *Pinus contorta* | 4 |
| Densitometer | 5 |
| Habit:tree | 5 |
| MEM1 | 5 |
| Slope perc. | 5 |
| Leaf area | 6 |
| Height of sample | 7 |
| Tree richness | 7 |
| Longitude | 8 |
| Sla | 8 |
| Dead and down | 9 |
| MEM2 | 9 |
| Precip. in April | 9 |
| Latitude | 10 |
| Mean temp. April | 10 |
| Julian date | 11 |
| elevation | 12 |
| Shrub richness | 12 |

**Table S5:**
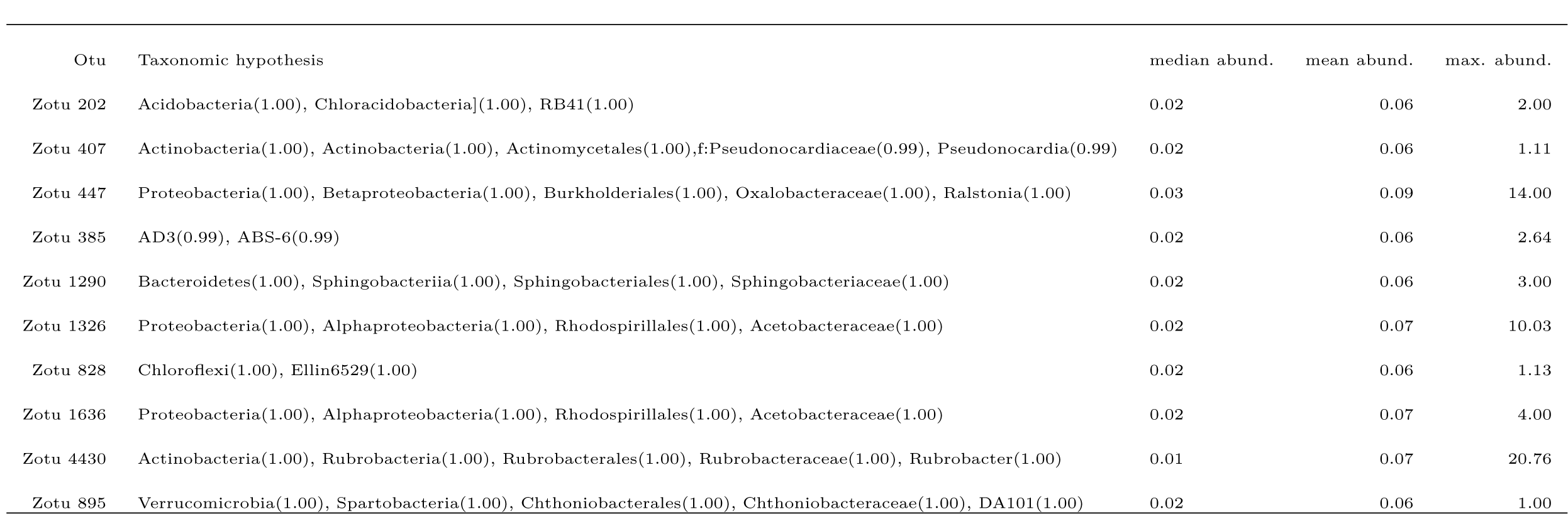
The most important bacterial taxa for models that included bacteria as features. Taxonomic hypotheses were generated using the Greengenes database. The numbers in parentheses are the proportion estimates for how certain the taxon was correctly identified, as output by SINTAX. Otu name is presented (i.e., Zotu 207) to aid in searching for the OTU in sequencing data. Abundances shown are ISD normalized data.

**Table S6:**
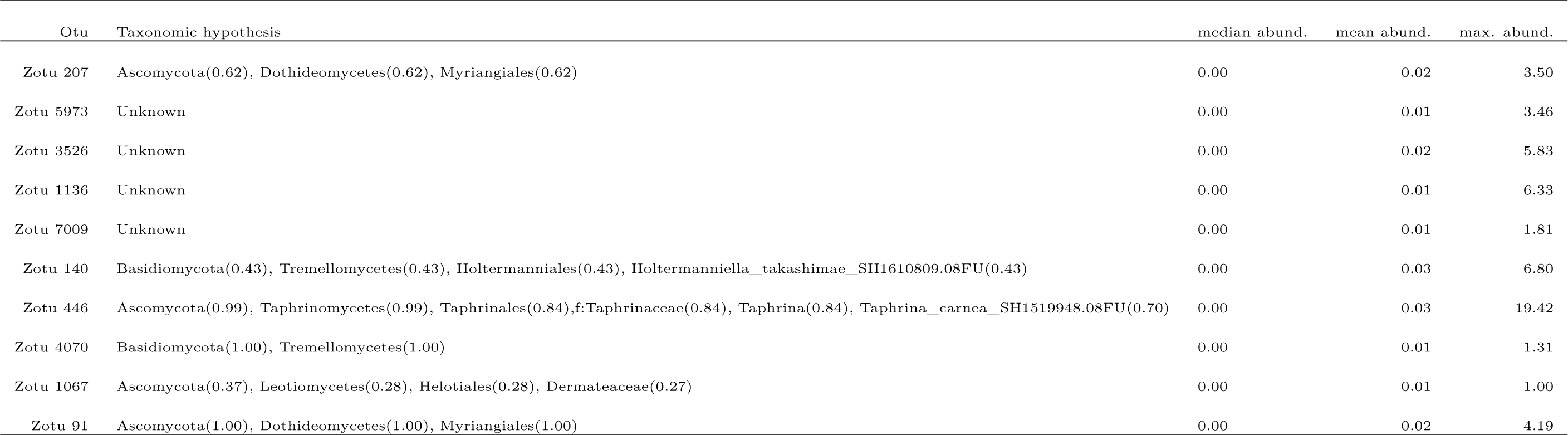
The most important fungal taxa for models that included fungi as features. Taxonomic hypotheses were generated using the UNITE database. The numbers in parentheses are the proportion estimates for how certain the taxon was correctly identified, as output by SINTAX. Otu name is presented (i.e., Zotu 207) to aid in searching for the OTU in sequencing data. Abundances shown are ISD normalized data.

**Table S7:** Number of times a feature was in the top ten most important for models of fungal abundance (ISD standardized) that were limited to a single host taxon and which had an R^2^ > 0.01. Only the fifteen most important features are shown. Models were successful for 16 out of 110 combinations of host and microbial taxon.

| Feature | Num. of times in top ten most important. |
| --- | --- |
| Rel. chlorophyll intensity | 13 |
| Compartment: EN | 13 |
| Phenology: fruiting | 12 |
| Tree richness | 11 |
| SPAD 420 intensity | 11 |
| Phenology: vegetative | 11 |
| Leaves extracted | 11 |
| Densitometer | 9 |
| Slope perc. | 7 |
| gH. | 7 |
| Absorbance 940 | 5 |
| Latitude | 4 |
| Height of sample | 4 |
| Shrub richness | 3 |
| Julian date | 3 |

**Table S8:** Number of times a feature was in the top ten most important for models of fungal *relative abundance* (Hellinger transformed) that were limited to a single host taxon and which had an *R*^2^ *>* 0.01. Only the fifteen most important features are shown. Models were successful for 66 out of 110 combinations of host and microbial taxon.

| Feature | Num. of times in top ten most important. |
| --- | --- |
| SPAD 420 intensity | 63 |
| Rel. chlorophyll intensity | 59 |
| Compartment: EN | 57 |
| Phenology: vegetative | 55 |
| Densitometer | 55 |
| Phenology: fruiting | 54 |
| gH. | 54 |
| Tree richness | 53 |
| Shrub richness | 53 |
| Leaves extracted | 41 |
| Slope perc. | 25 |
| MEM1 | 8 |
| Julian date | 8 |
| Dead and down | 8 |
| Phenology: flowering | 7 |

**Table S9:** Number of times a feature was in the top ten most important for models of fungal occupancy that were limited to a single host taxon and that had an MCC greater than 0.2. The fifteen most important features are shown. Models were successful for 81 out of 110 combinations of host and microbial taxon.

| Feature | Num. of times in top ten most important. |
| --- | --- |
| SPAD 420 intensity | 42 |
| Compartment: EN | 42 |
| Tree richness | 40 |
| Rel. chlorophyll intensity | 38 |
| gH. | 38 |
| Phenology: vegetative | 37 |
| Densitometer | 37 |
| Shrub richness | 36 |
| Phenology: fruiting | 33 |
| Leaves extracted | 26 |
| Slope perc. | 16 |
| Phenology: flowering | 6 |
| Julian date | 6 |
| Plant volume | 4 |
| Dead and down | 4 |

**Table S10:** Differences in estimated fungal Shannon’s diversity by compartment and host growth habit. EN refers to endophytes and EP refers to epiphytes. The estimated difference in diversity is shown with 95% confidence intervals and a p value adjusted for multiple comparisons.

| Comparison | diff | lwr | upr | p adj |
| --- | --- | --- | --- | --- |
| EN graminoid-EN forb | 17.43 | -371.38 | 406.25 | 1.00 |
| EN shrub-EN forb | -83.72 | -314.04 | 146.61 | 0.96 |
| EN tree-EN forb | -273.27 | -511.74 | -34.79 | 0.01 |
| EP forb-EN forb | 383.26 | 195.33 | 571.18 | 0.00 |
| EP graminoid-EN forb | 380.05 | -4.34 | 764.44 | 0.06 |
| EP shrub-EN forb | 150.86 | -78.93 | 380.66 | 0.49 |
| EP tree-EN forb | 116.34 | -128.20 | 360.87 | 0.84 |
| EN shrub-EN graminoid | -101.15 | -512.03 | 309.73 | 1.00 |
| EN tree-EN graminoid | -290.70 | -706.21 | 124.80 | 0.40 |
| EP forb-EN graminoid | 365.82 | -22.88 | 754.52 | 0.08 |
| EP graminoid-EN graminoid | 362.62 | -150.73 | 875.97 | 0.39 |
| EP shrub-EN graminoid | 133.43 | -277.15 | 544.01 | 0.98 |
| EP tree-EN graminoid | 98.90 | -320.11 | 517.91 | 1.00 |
| EN tree-EN shrub | -189.55 | -462.52 | 83.42 | 0.41 |
| EP forb-EN shrub | 466.97 | 236.85 | 697.10 | 0.00 |
| EP graminoid-EN shrub | 463.77 | 57.07 | 870.46 | 0.01 |
| EP shrub-EN shrub | 234.58 | -30.84 | 500.00 | 0.13 |
| EP tree-EN shrub | 200.05 | -78.23 | 478.33 | 0.36 |
| EP forb-EN tree | 656.53 | 418.24 | 894.81 | 0.00 |
| EP graminoid-EN tree | 653.32 | 241.96 | 1064.68 | 0.00 |
| EP shrub-EN tree | 424.13 | 151.61 | 696.65 | 0.00 |
| EP tree-EN tree | 389.60 | 104.54 | 674.66 | 0.00 |
| EP graminoid-EP forb | -3.21 | -387.48 | 381.07 | 1.00 |
| EP shrub-EP forb | -232.39 | -461.99 | -2.80 | 0.04 |
| EP tree-EP forb | -266.92 | -511.27 | -22.58 | 0.02 |
| EP shrub-EP<br>graminoid | -229.19 | -635.58 | 177.21 | 0.68 |
| EP tree-EP<br>graminoid | -263.72 | -678.62 | 151.19 | 0.53 |
| EP tree-EP shrub | -34.53 | -312.37 | 243.31 | 1.00 |

**Table S11:** Differences in estimated bacterial Shannon’s diversity by compartment and host growth habit. EN refers to endophytes and EP refers to epiphytes. The estimated difference in diversity is shown with 95% confidence intervals and a p value adjusted for multiple comparisons.

| Comparison | diff | lwr | upr | p adj |
| --- | --- | --- | --- | --- |
| EN graminoid-EN<br>forb | 38.43 | -60.14 | 137.00 | 0.94 |
| EN shrub-EN forb | 11.21 | -47.09 | 69.51 | 1.00 |
| EN tree-EN forb | -15.59 | -75.79 | 44.60 | 0.99 |
| EP forb-EN forb | 7.37 | -39.82 | 54.57 | 1.00 |
| EP graminoid-EN<br>forb | -15.29 | -112.12 | 81.55 | 1.00 |
| EP shrub-EN forb | -18.55 | -76.31 | 39.20 | 0.98 |
| EP tree-EN forb | -7.85 | -69.43 | 53.73 | 1.00 |
| EN shrub-EN<br>graminoid | -27.22 | -131.32 | 76.89 | 0.99 |
| EN tree-EN<br>graminoid | -54.02 | -159.20 | 51.16 | 0.78 |
| EP forb-EN<br>graminoid | -31.05 | -129.37 | 67.26 | 0.98 |
| EP graminoid-EN<br>graminoid | -53.72 | -183.39 | 75.96 | 0.91 |
| EP shrub-EN<br>graminoid | -56.98 | -160.78 | 46.82 | 0.71 |
| EP tree-EN<br>graminoid | -46.28 | -152.25 | 59.70 | 0.89 |
| EN tree-EN shrub | -26.80 | -95.69 | 42.08 | 0.94 |
| EP forb-EN shrub | -3.84 | -61.71 | 54.04 | 1.00 |
| EP graminoid-EN<br>shrub | -26.50 | -128.96 | 75.97 | 0.99 |
| EP shrub-EN<br>shrub | -29.76 | -96.53 | 37.00 | 0.88 |
| EP tree-EN shrub | -19.06 | -89.16 | 51.04 | 0.99 |
| EP forb-EN tree | 22.97 | -36.81 | 82.75 | 0.94 |
| EP graminoid-EN<br>tree | 0.30 | -103.25 | 103.86 | 1.00 |
| EP shrub-EN tree | -2.96 | -71.38 | 65.46 | 1.00 |
| EP tree-EN tree | 7.74 | -63.94 | 79.42 | 1.00 |
| EP graminoid-EP<br>forb | -22.66 | -119.24 | 73.92 | 1.00 |
| EP shrub-EP forb | -25.93 | -83.25 | 31.40 | 0.87 |
| EP tree-EP forb | -15.22 | -76.40 | 45.95 | 1.00 |
| EP shrub-EP<br>graminoid | -3.26 | -105.42 | 98.89 | 1.00 |
| EP tree-EP<br>graminoid | 7.44 | -96.93 | 111.80 | 1.00 |
| EP tree-EP shrub | 10.70 | -58.94 | 80.34 | 1.00 |

**Table S12:** Differences in estimated bacterial richness by compartment and host growth habit. EN refers to endophytes and EP refers to epiphytes. The estimated difference in richness is shown with 95% confidence intervals and a p value adjusted for multiple comparisons. The random forest model for 16s richness was not successful (*R*^2^∼=0).

| Comparison | Diff. | Lower bound | Upper bound | Adj. p |
| --- | --- | --- | --- | --- |
| forb EP-forb EN | 0.76 | 0.37 | 1.15 | 0.00 |
| graminoid<br>EN-forb EN | -0.46 | -1.31 | 0.40 | 0.74 |
| graminoid EP-forb<br>EN | 0.74 | -0.00 | 1.48 | 0.05 |
| shrub EN-forb EN | 0.14 | -0.36 | 0.63 | 0.99 |
| shrub EP-forb EN | 0.82 | 0.37 | 1.28 | 0.00 |
| tree EN-forb EN | 0.18 | -0.31 | 0.67 | 0.96 |
| tree EP-forb EN | 0.74 | 0.26 | 1.23 | 0.00 |
| graminoid<br>EN-forb EP | -1.22 | -2.06 | -0.38 | 0.00 |
| graminoid EP-forb<br>EP | -0.02 | -0.75 | 0.71 | 1.00 |
| shrub EN-forb EP | -0.62 | -1.10 | -0.15 | 0.00 |
| shrub EP-forb EP | 0.07 | -0.36 | 0.49 | 1.00 |
| tree EN-forb EP | -0.58 | -1.05 | -0.12 | 0.00 |
| tree EP-forb EP | -0.02 | -0.48 | 0.44 | 1.00 |
| graminoid<br>EP-graminoid EN | 1.20 | 0.15 | 2.25 | 0.01 |
| shrub<br>EN-graminoid EN | 0.59 | -0.30 | 1.48 | 0.48 |
| shrub<br>EP-graminoid EN | 1.28 | 0.41 | 2.15 | 0.00 |
| tree EN-graminoid<br>EN | 0.63 | -0.26 | 1.52 | 0.38 |
| tree EP-graminoid<br>EN | 1.20 | 0.31 | 2.09 | 0.00 |
| shrub<br>EN-graminoid EP | -0.61 | -1.40 | 0.18 | 0.28 |
| shrub<br>EP-graminoid EP | 0.08 | -0.68 | 0.85 | 1.00 |
| tree EN-graminoid<br>EP | -0.56 | -1.35 | 0.22 | 0.37 |
| tree EP-graminoid<br>EP | 0.00 | -0.78 | 0.79 | 1.00 |
| shrub EP-shrub<br>EN | 0.69 | 0.16 | 1.22 | 0.00 |
| tree EN-shrub EN | 0.04 | -0.52 | 0.60 | 1.00 |
| tree EP-shrub EN | 0.61 | 0.05 | 1.16 | 0.02 |
| tree EN-shrub EP | -0.65 | -1.17 | -0.12 | 0.00 |
| tree EP-shrub EP | -0.08 | -0.60 | 0.44 | 1.00 |
| tree EP-tree EN | 0.57 | 0.01 | 1.12 | 0.04 |

**Table S13:** Differences in estimated fungal richness by compartment and host growth habit. EN refers to endophytes and EP refers to epiphytes. The estimated difference in richness is shown with 95% confidence intervals and a p value adjusted for multiple comparisons. The random forest model for 16s richness was not successful (R^2^∼=0).

| Comparison | Diff. | Lower<br>bound | Upper<br>bound | Adj. p |
| --- | --- | --- | --- | --- |
| forb EP-forb EN | -0.36 | -0.65 | -0.07 | 0.00 |
| graminoid<br>EN-forb EN | -0.40 | -0.99 | 0.18 | 0.41 |
| graminoid EP-forb<br>EN | -0.23 | -0.81 | 0.36 | 0.94 |
| shrub EN-forb EN | 0.02 | -0.32 | 0.36 | 1.00 |
| shrub EP-forb EN | 0.02 | -0.32 | 0.36 | 1.00 |
| tree EN-forb EN | 0.07 | -0.29 | 0.43 | 1.00 |
| tree EP-forb EN | 0.19 | -0.18 | 0.56 | 0.75 |
| graminoid<br>EN-forb EP | -0.04 | -0.63 | 0.54 | 1.00 |
| graminoid EP-forb<br>EP | 0.13 | -0.45 | 0.72 | 1.00 |
| shrub EN-forb EP | 0.38 | 0.04 | 0.73 | 0.01 |
| shrub EP-forb EP | 0.38 | 0.04 | 0.72 | 0.02 |
| tree EN-forb EP | 0.43 | 0.07 | 0.79 | 0.01 |
| tree EP-forb EP | 0.55 | 0.18 | 0.93 | 0.00 |
| graminoid<br>EP-graminoid EN | 0.18 | -0.60 | 0.95 | 1.00 |
| shrub<br>EN-graminoid EN | 0.43 | -0.18 | 1.04 | 0.40 |
| shrub<br>EP-graminoid EN | 0.42 | -0.19 | 1.03 | 0.43 |
| tree EN-graminoid<br>EN | 0.47 | -0.15 | 1.09 | 0.29 |
| tree EP-graminoid<br>EN | 0.60 | -0.03 | 1.23 | 0.08 |
| shrub<br>EN-graminoid EP | 0.25 | -0.36 | 0.86 | 0.92 |
| shrub<br>EP-graminoid EP | 0.24 | -0.37 | 0.86 | 0.93 |
| tree EN-graminoid<br>EP | 0.30 | -0.33 | 0.92 | 0.83 |
| tree EP-graminoid<br>EP | 0.42 | -0.21 | 1.05 | 0.46 |
| shrub EP-shrub<br>EN | -0.01 | -0.40 | 0.38 | 1.00 |
| tree EN-shrub EN | 0.05 | -0.36 | 0.45 | 1.00 |
| tree EP-shrub EN | 0.17 | -0.24 | 0.58 | 0.92 |
| tree EN-shrub EP | 0.05 | -0.35 | 0.46 | 1.00 |
| tree EP-shrub EP | 0.18 | -0.24 | 0.59 | 0.90 |
| tree EP-tree EN | 0.12 | -0.30 | 0.55 | 0.99 |

**Table S14:** Median fungal epiphyte richness of host plants. Only the top 20 most rich are shown.

| Host taxon | Median richness | Habit |
| --- | --- | --- |
| Unknown fir | 40.48 | tree |
| Wyethia amplexicaulis | 41.71 | forb |
| Arnica cordifolia | 44.00 | forb |
| Potentilla pulcherrima | 44.59 | forb |
| Abies concolor | 47.95 | tree |
| Picea engelmannii | 48.80 | tree |
| Helianthella uniflora | 53.11 | forb |
| Symphoricarpos albus | 53.65 | shrub |
| Pseudotsuga menziesii | 54.34 | tree |
| Unknown Spruce | 56.14 | tree |
| Ribes montigenum | 61.44 | shrub |
| Sedum lanceolatum | 62.49 | forb |
| Eriogonum umbellatum | 62.95 | forb |
| Minuartia obtusiloba | 73.93 | forb |
| Abies grandis | 76.98 | tree |
| Juniperus communis | 77.00 | shrub |
| Poa wheeleri | 82.10 | graminoid |
| Vaccinium membranaceum | 88.62 | shrub |
| Pinus contorta | 123.15 | tree |
| Paxistima myrsinites | 212.51 | shrub |

**Table S15:** Median fungal endophyte richness of host plants. Only top 20 most rich host taxa shown.

| Host taxon | Median richness | Habit |
| --- | --- | --- |
| *Symphoricarpos albus* | 42.38 | shrub |
| *Vaccinium scoparium* | 42.51 | shrub |
| *Ribes montigenum* | 43.73 | shrub |
| *Sedum lanceolatum* | 46.32 | forb |
| *Heracleum maximum* | 46.55 | forb |
| Unknown fir | 48.45 | tree |
| *Fragaria virginiana* | 49.56 | forb |
| *Pinus contorta* | 56.62 | tree |
| *Pseudotsuga menziesii* | 60.44 | tree |
| *Eriogonum umbellatum* | 61.35 | forb |
| *Astragalus kentrophyta* | 63.22 | forb |
| *Juniperus communis* | 74.34 | shrub |
| *Minuartia obtusiloba* | 77.87 | forb |
| *Poa pratensis* | 80.95 | graminoid |
| *Abies grandis* | 83.27 | tree |
| *Osmorhiza depauperata* | 89.40 | forb |
| *Vaccinium membranaceum* | 93.71 | shrub |
| *Abies concolor* | 119.11 | tree |
| *Antennaria media* | 179.80 | forb |
| *Paxistima myrsinites* | 289.50 | shrub |

**Figure S1:**
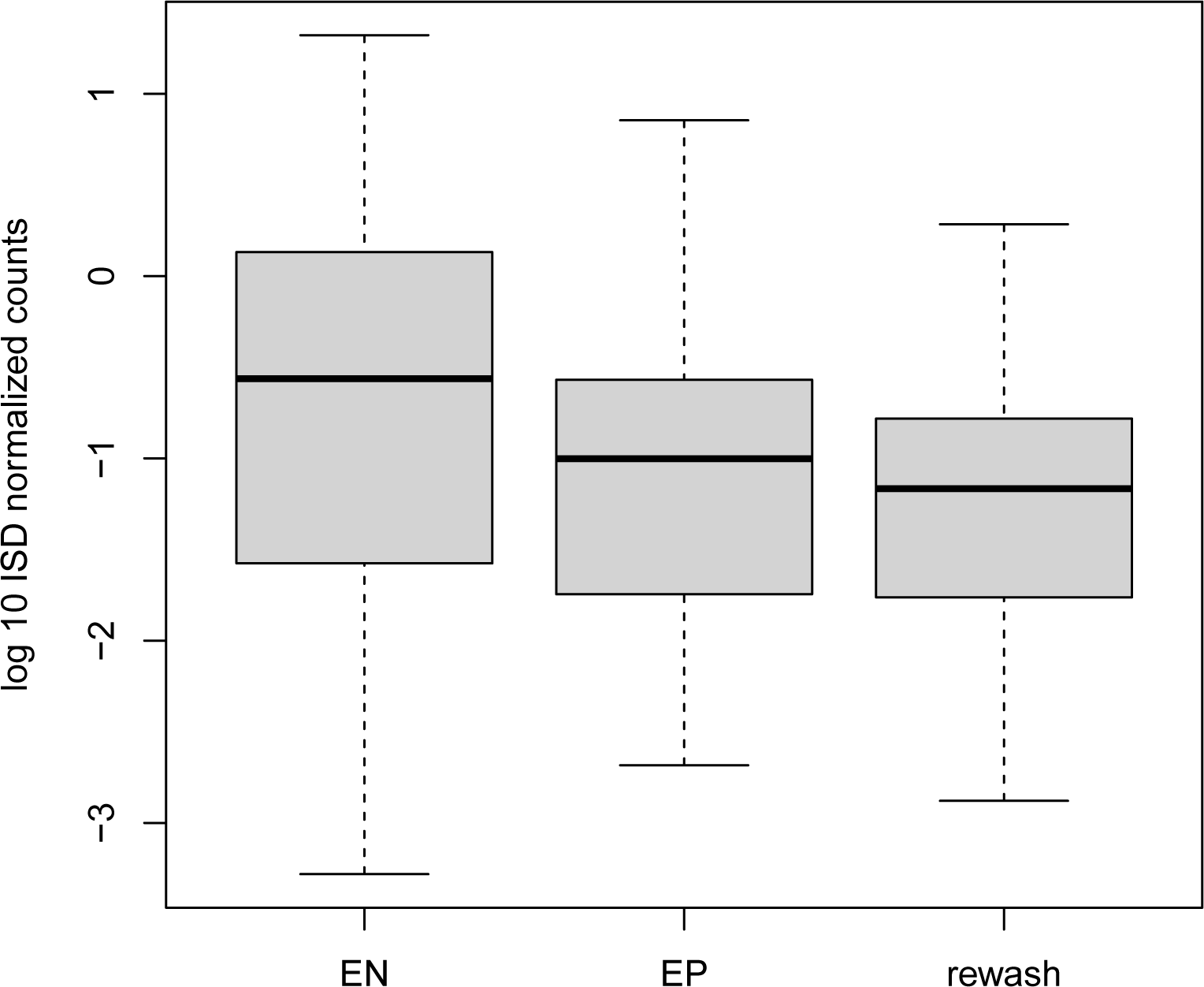
Boxplot showing read abundances from samples that were washed multiple times to determine the efficacy of our epiphyte removal technique. Data were divided by the internal standard to place them on a standard scale. For bacterial data, see Fig. S2

**Figure S2:**
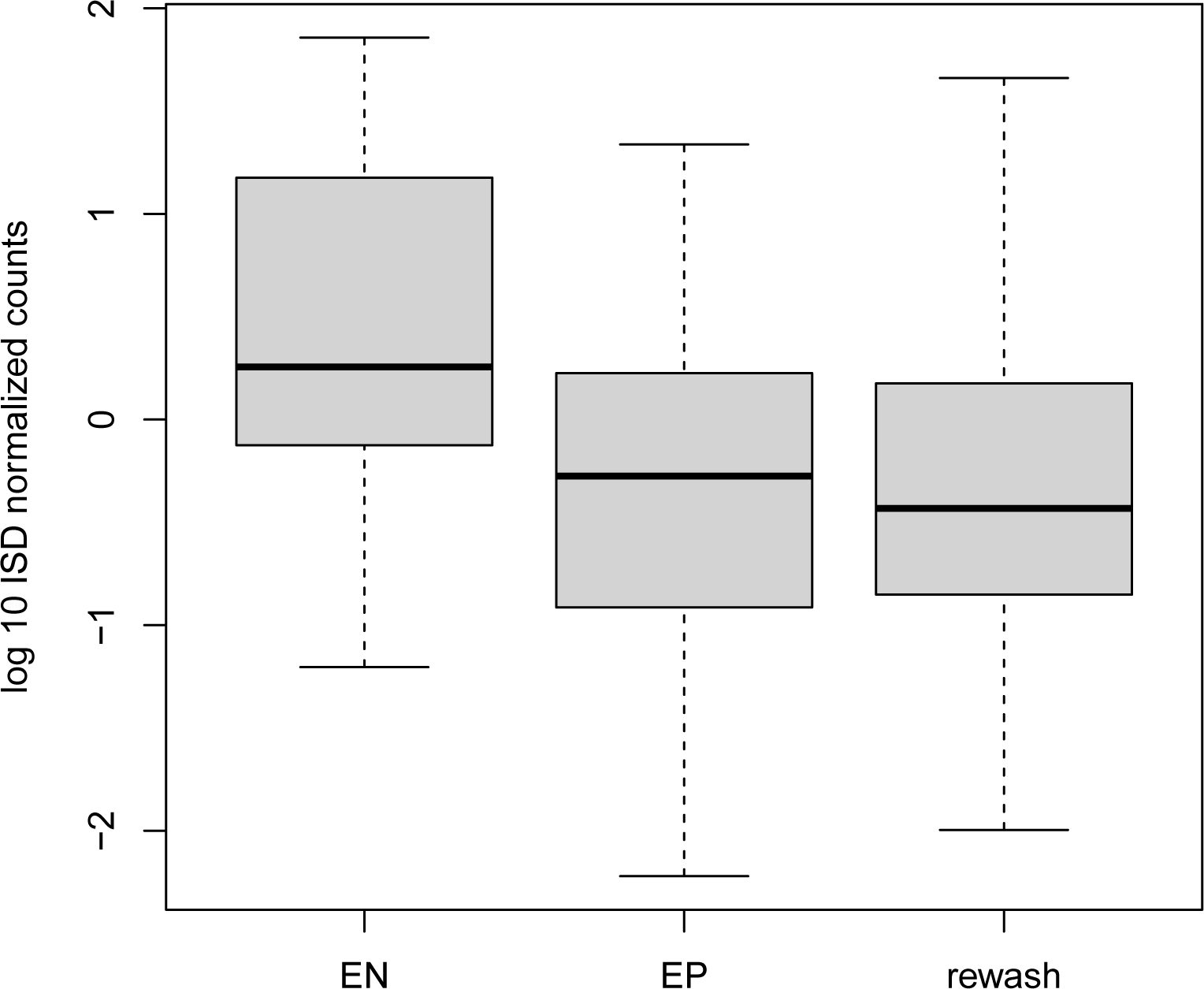
Boxplot showing read abundances from samples that were washed multiple times to determine the efficacy of our epiphyte removal technique. Data were divided by the internal standard to account for compositionality. For fungal data see, Fig. S1

**Figure S3:**
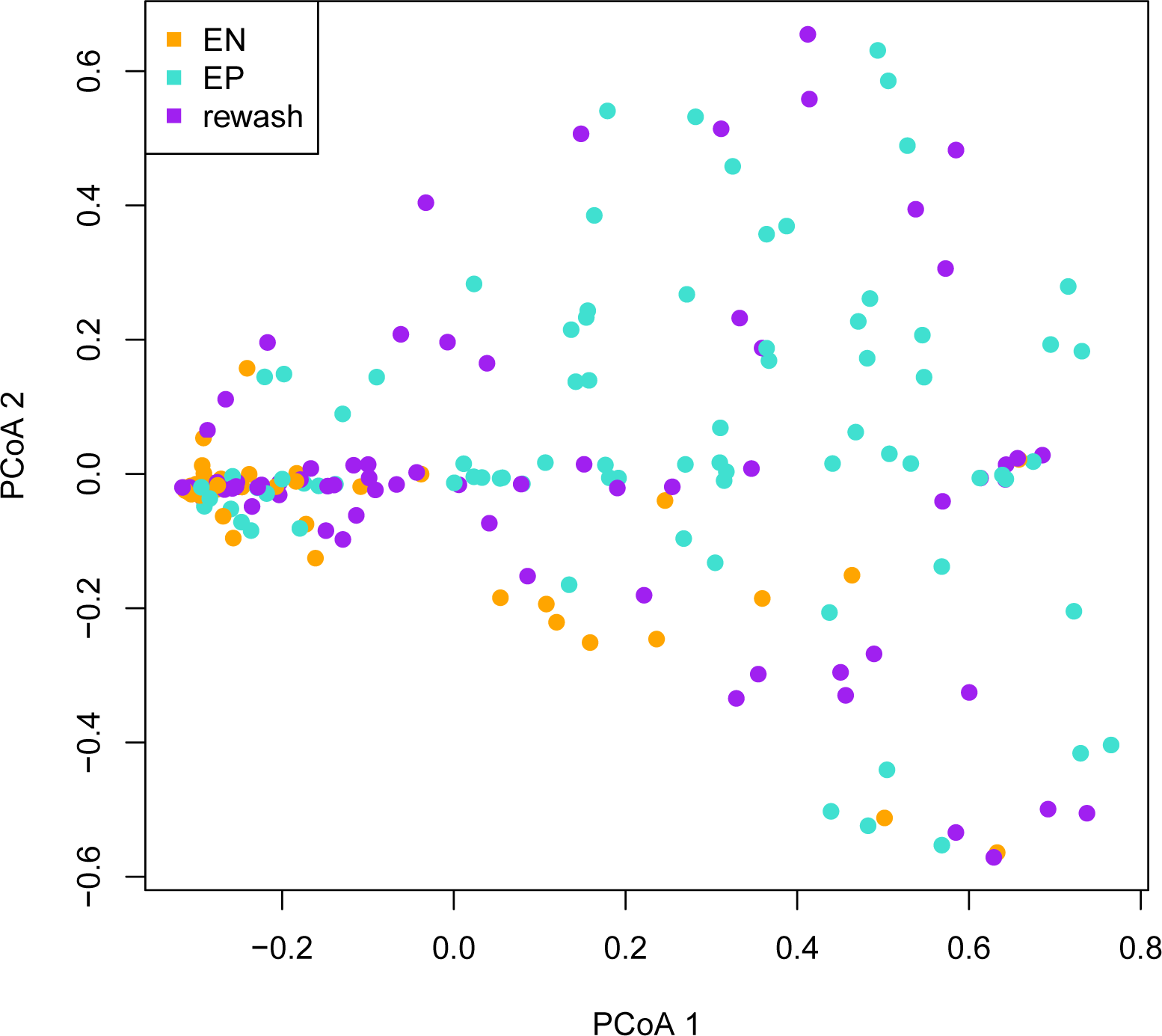
Principal coordinates ordination of samples that were washed multiple times to determine the efficacy of our epiphyte removal technique. Data were Hellinger transformed *bacterial* count data converted to a Euclidean distance matrix. A PERMANOVA by treatment (one of “EP”,“EN”,“rewash”) was significant.

**Figure S4:**
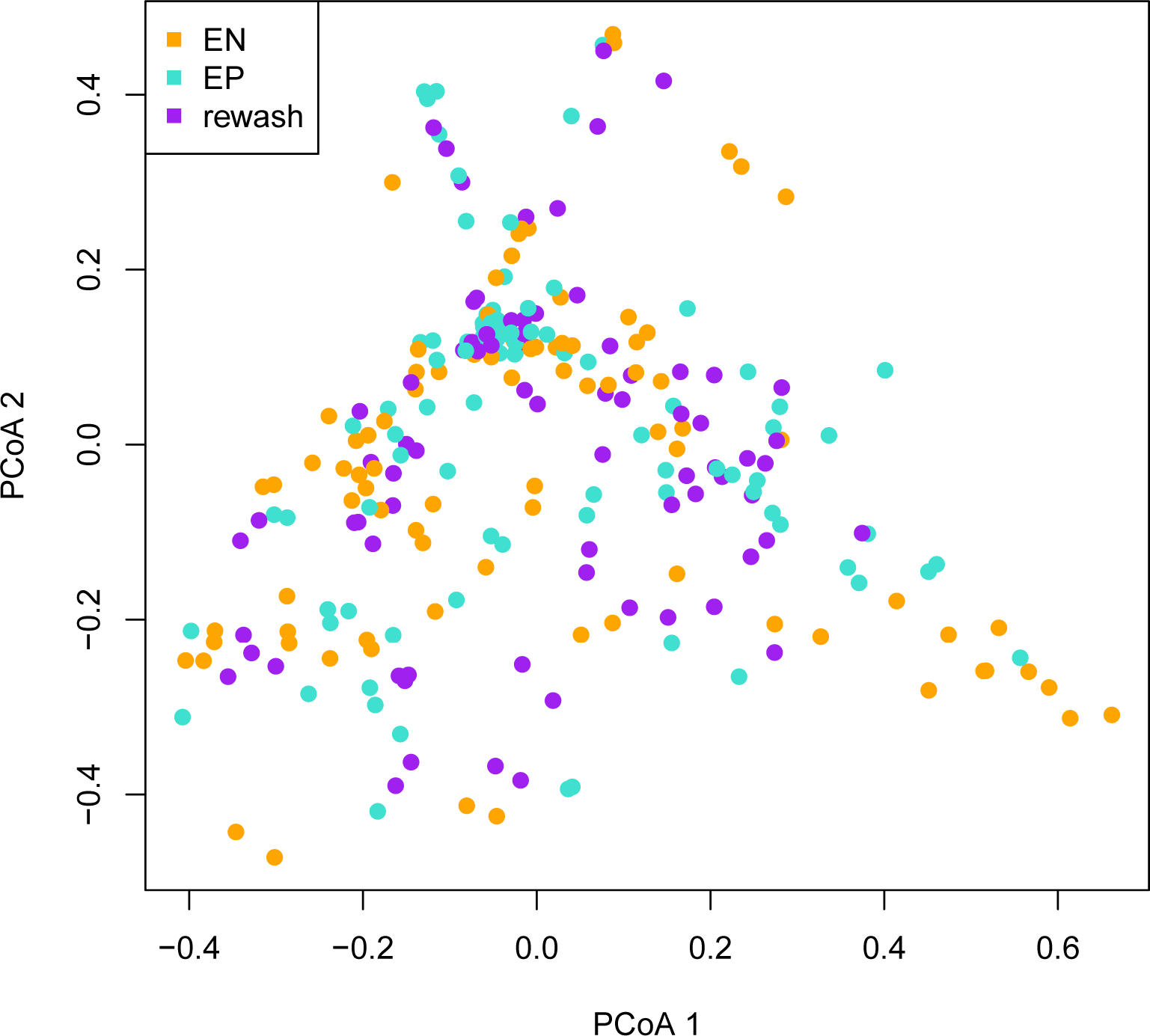
Principal coordinates ordination of samples that were washed multiple times to determine the efficacy of our epiphyte removal technique. Data were Hellinger transformed *fungal* count data converted to a Euclidean distance matrix. A PERMANOVA by treatment (one of “EP”,“EN”,“rewash”) was not significant.

**Figure S5:**
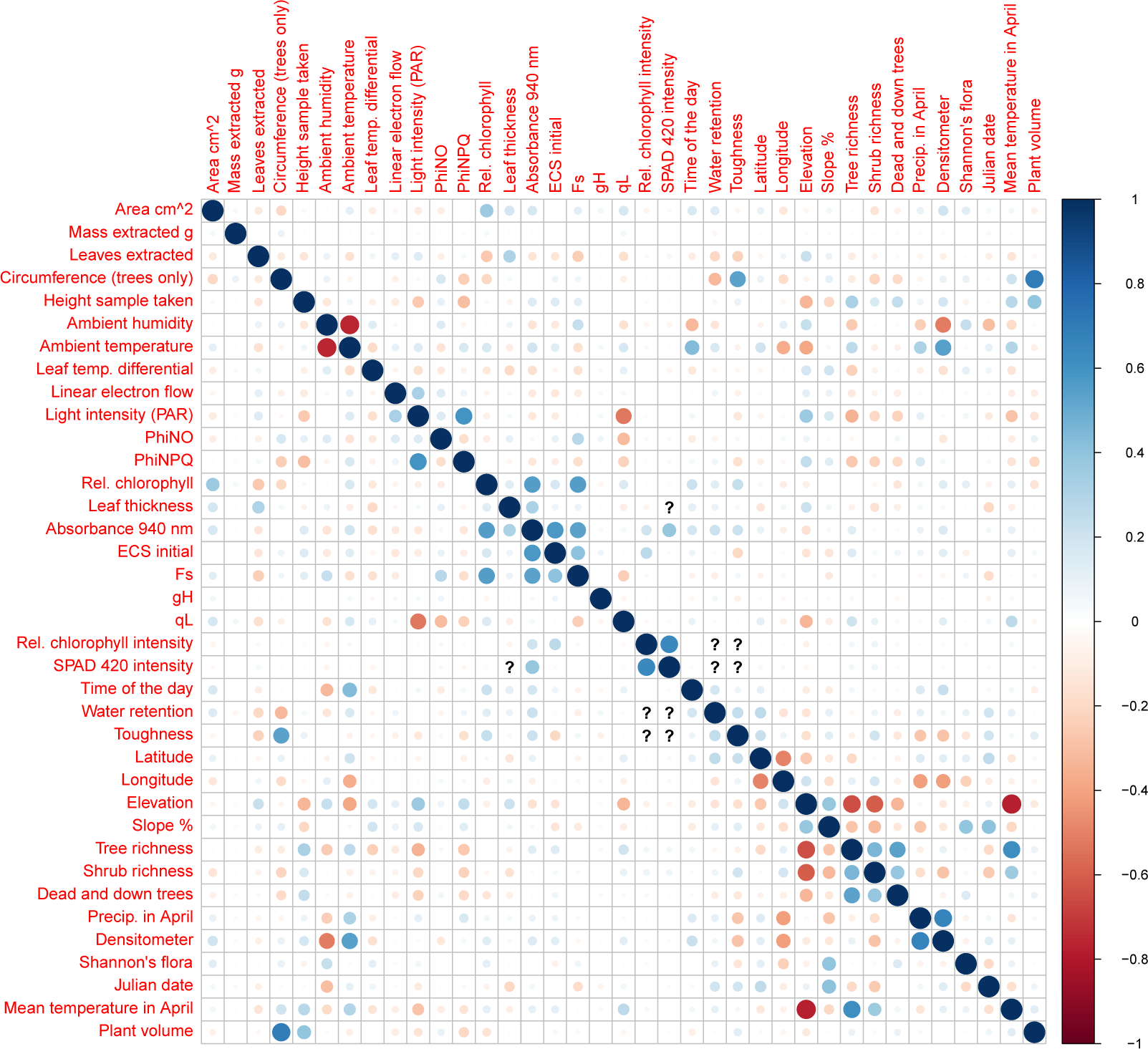
Pearson’s pairwise correlations among features used in random forest models of microbial relative abundances. Blue denotes positive correlation and red negative correlation. The strength of association is denoted via shading, as shown in the sidebar. Question marks denote instances where missing data in the features being associated coincided, thus preventing accurate correlation assessment. For a description of features, see Table 1.

**Figure S6:**
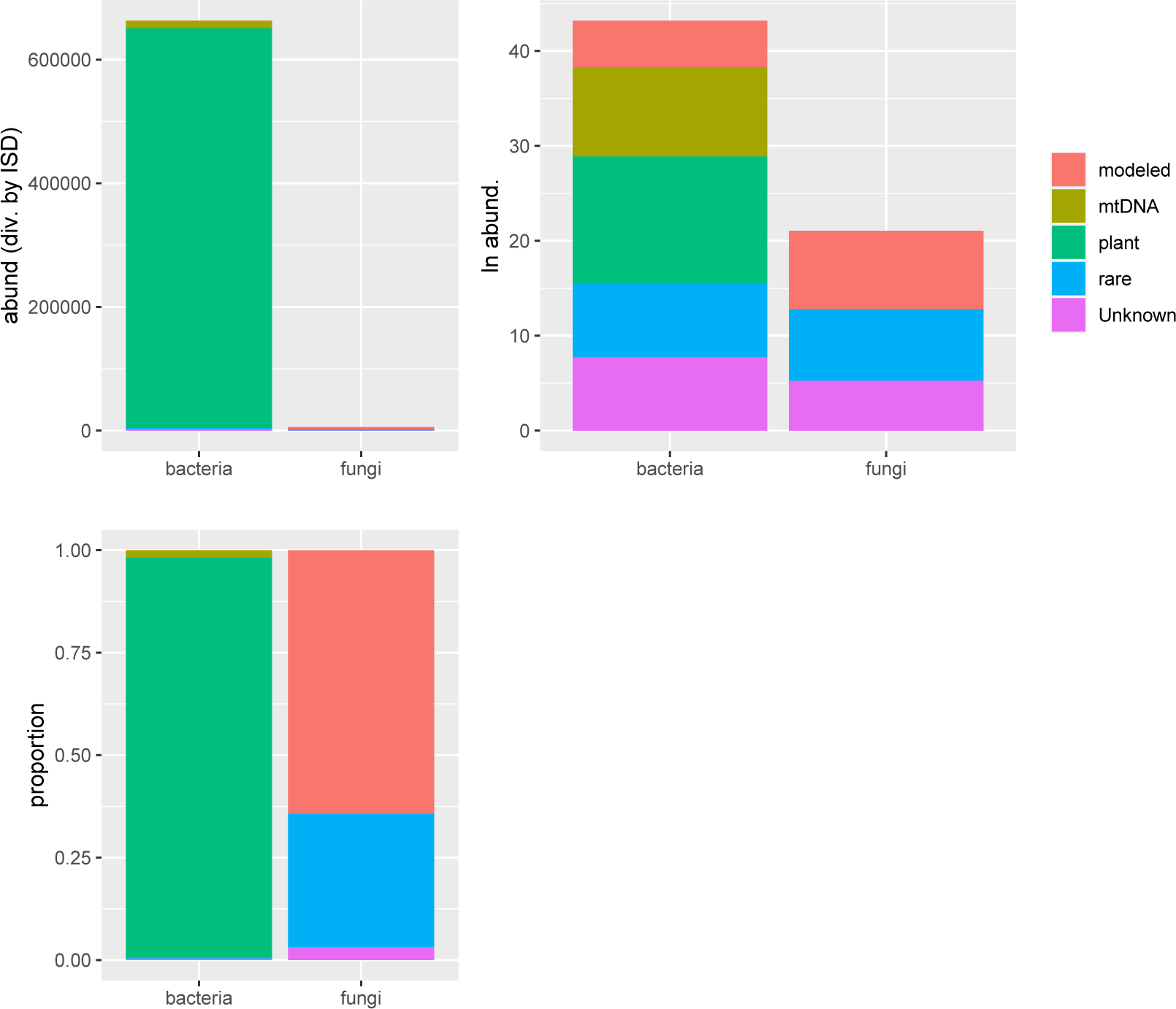
Read count data from various sources (e.g., plant, microbes) shown as abundances (divided by the ISD), logged abundances (natural log), and proportions. “modeled” refers to those taxa that were included in modeling efforts. “rare” refers to those taxa that were not modeled.

**Figure S7:**
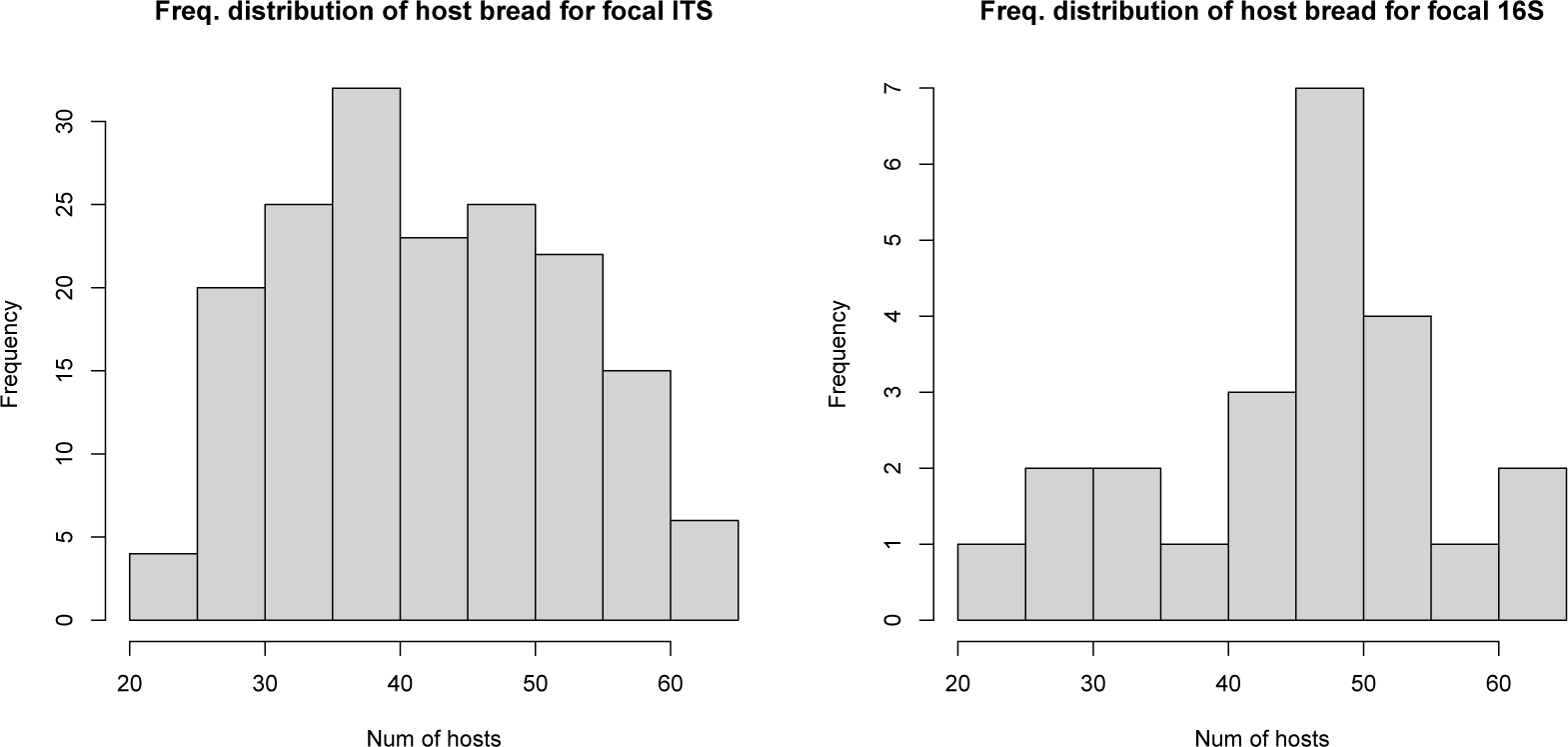
Frequency distribution of host use for fungal (left) and bacterial (right) taxa considered for our random forest analyses. These taxa were the most prevalent, and some of the most abundant, in our dataset.

**Figure S8:**
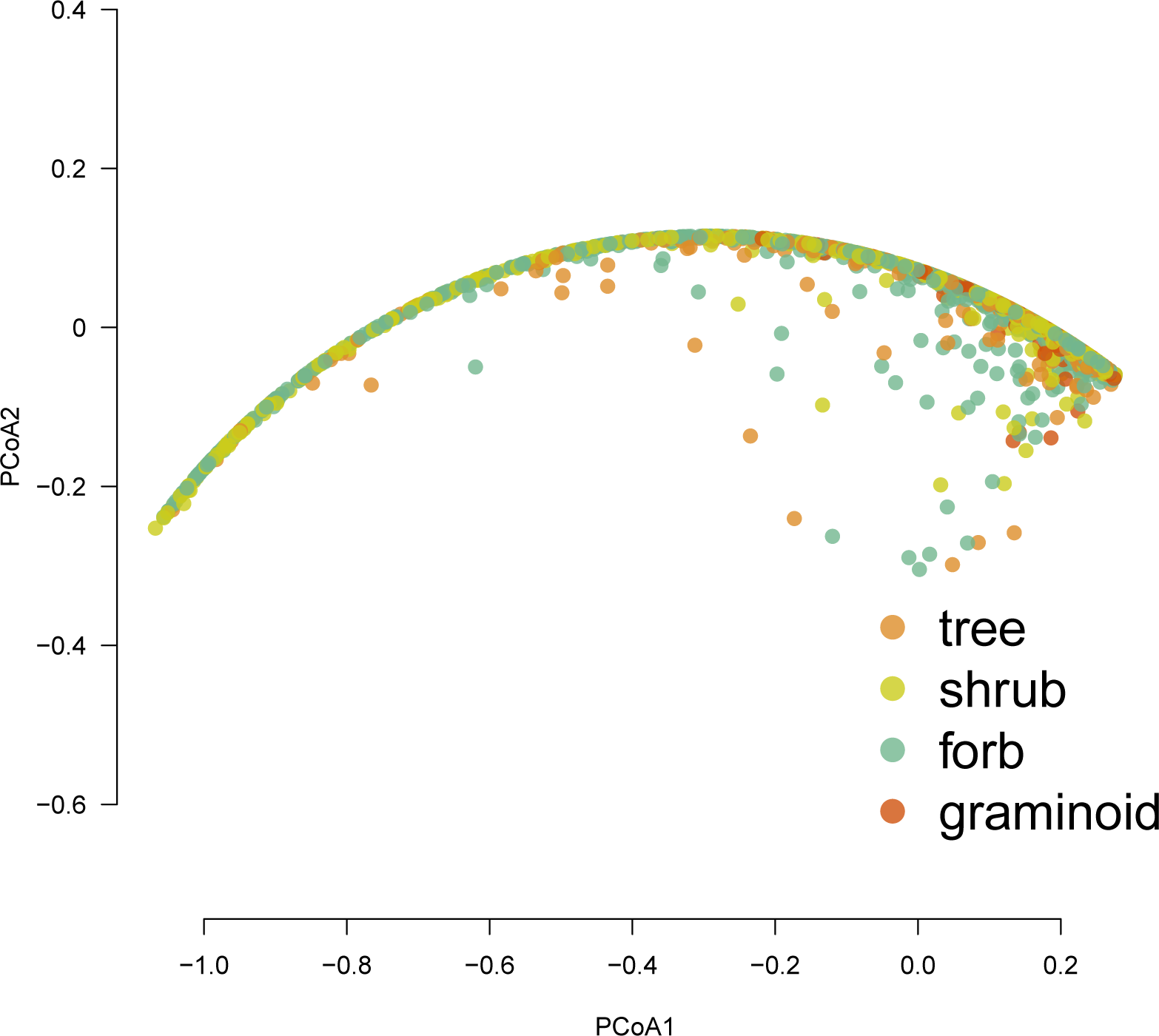
Principal coordinates analysis ordination of 16s (bacterial) data color-coded to reflect samples differing by plant life history. Data were normalized by the internal standard, Hellinger transformed, and converted to a Euclidean distance matrix. A PERMANOVA by treatment had an *R*^2^ =∼0.03 (*p* = 0.001), but the homogeneity of variances assumption of this test was violated.

**Figure S9:**
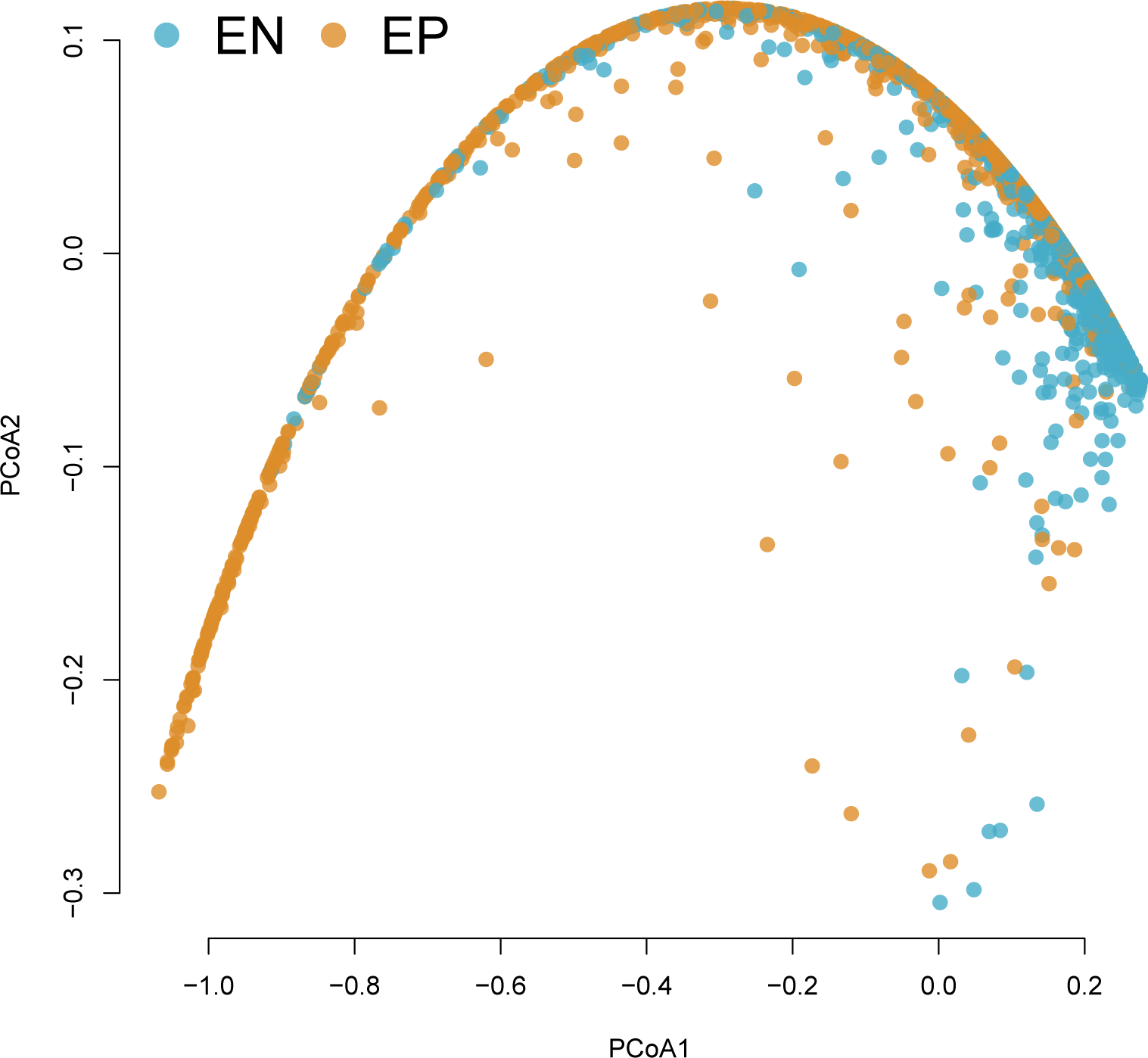
Principal coordinates analysis ordination of 16s (bacterial) data color-coded to reflect samples differing by compartment, either epiphyte (EP) or endophyte (EN). Data were normalized by the internal standard, Hellinger transformed, and converted to a Euclidean distance matrix. A PERMANOVA by treatment had an *R*^2^ =∼0.13 (*p* = 0.001), but the homogeneity of variances assumption of this test was violated.

**Figure S10:**
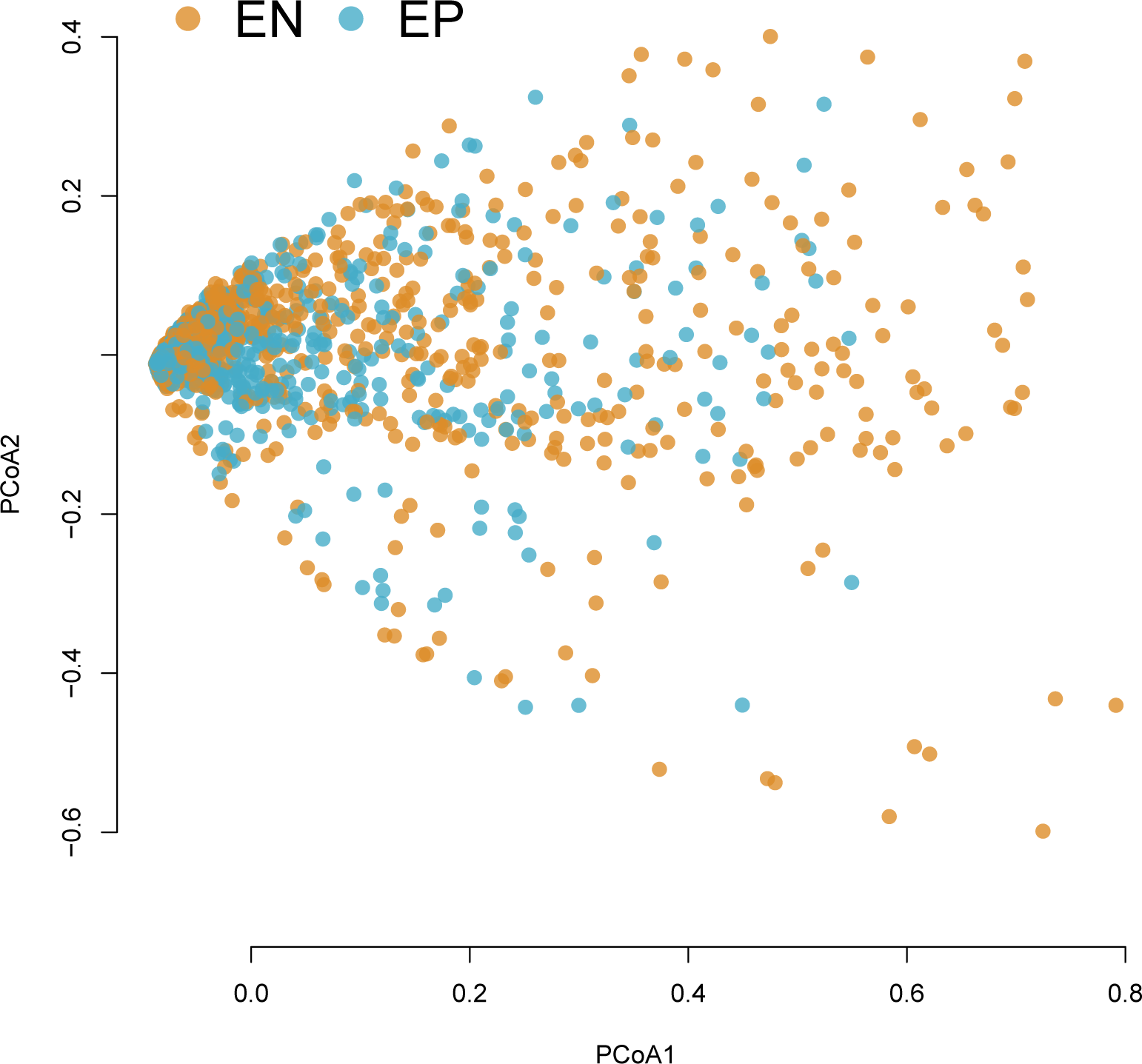
Principal coordinates analysis ordination of ITS (fungal) data color-coded to reflect samples differing by compartment, either epiphyte (EP) or endophyte (EN). Data were normalized by the internal standard, Hellinger transformed, and converted to a Euclidean distance matrix. A PERMANOVA by treatment had an *R*^2^ =∼0.01 (*p* = 0.001), but the homogeneity of variances assumption of this test was violated.

**Figure S11:**
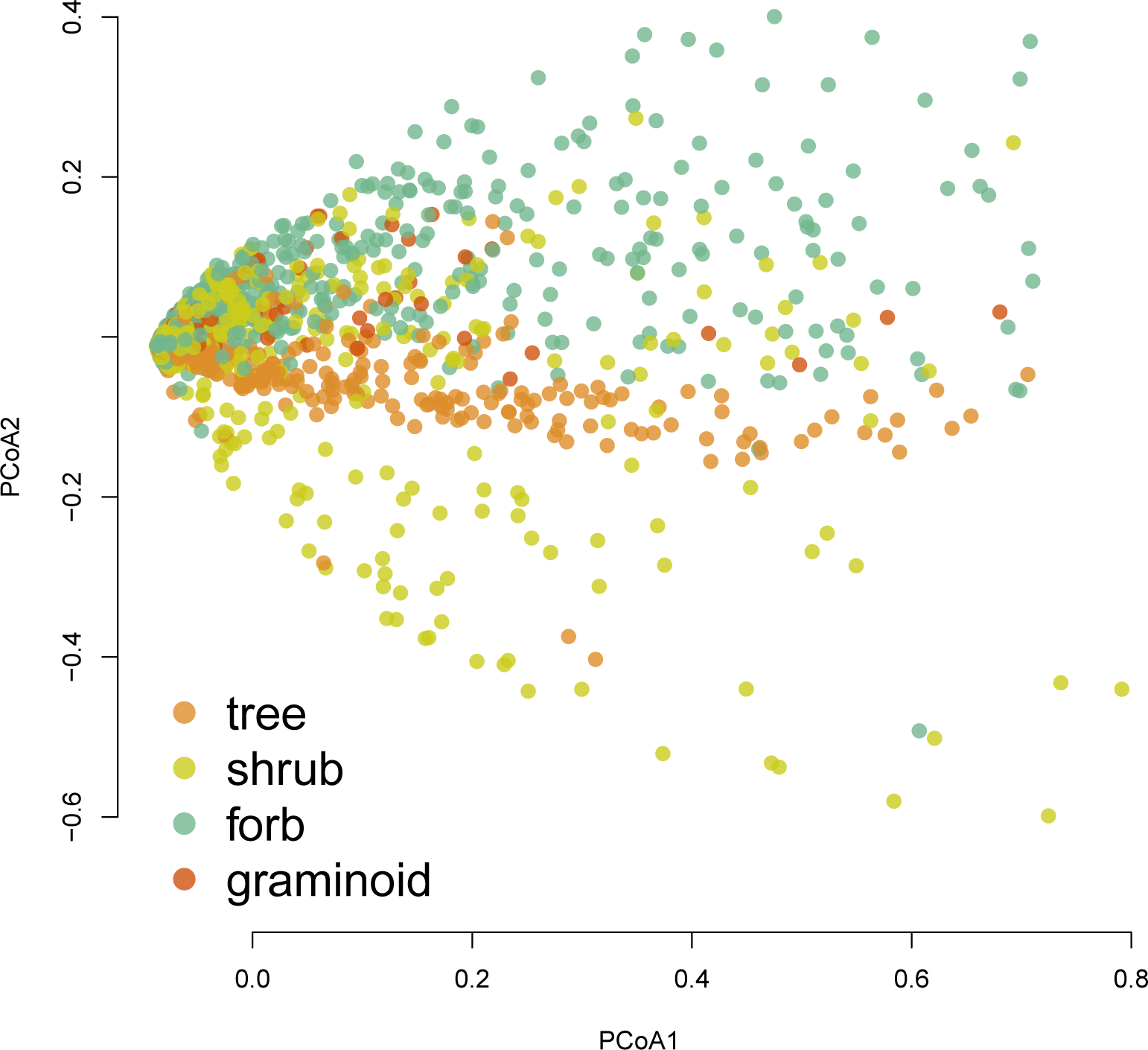
Principal coordinates analysis ordination of ITS (fungal) data color-coded to reflect samples differing by life history. Data were normalized by the internal standard, Hellinger transformed, and converted to a Euclidean distance matrix. A PERMANOVA by treatment had an *R*^2^ =∼0.02 (*p* = 0.001), but the homogeneity of variances assumption of this test was violated.

**Figure S12:**
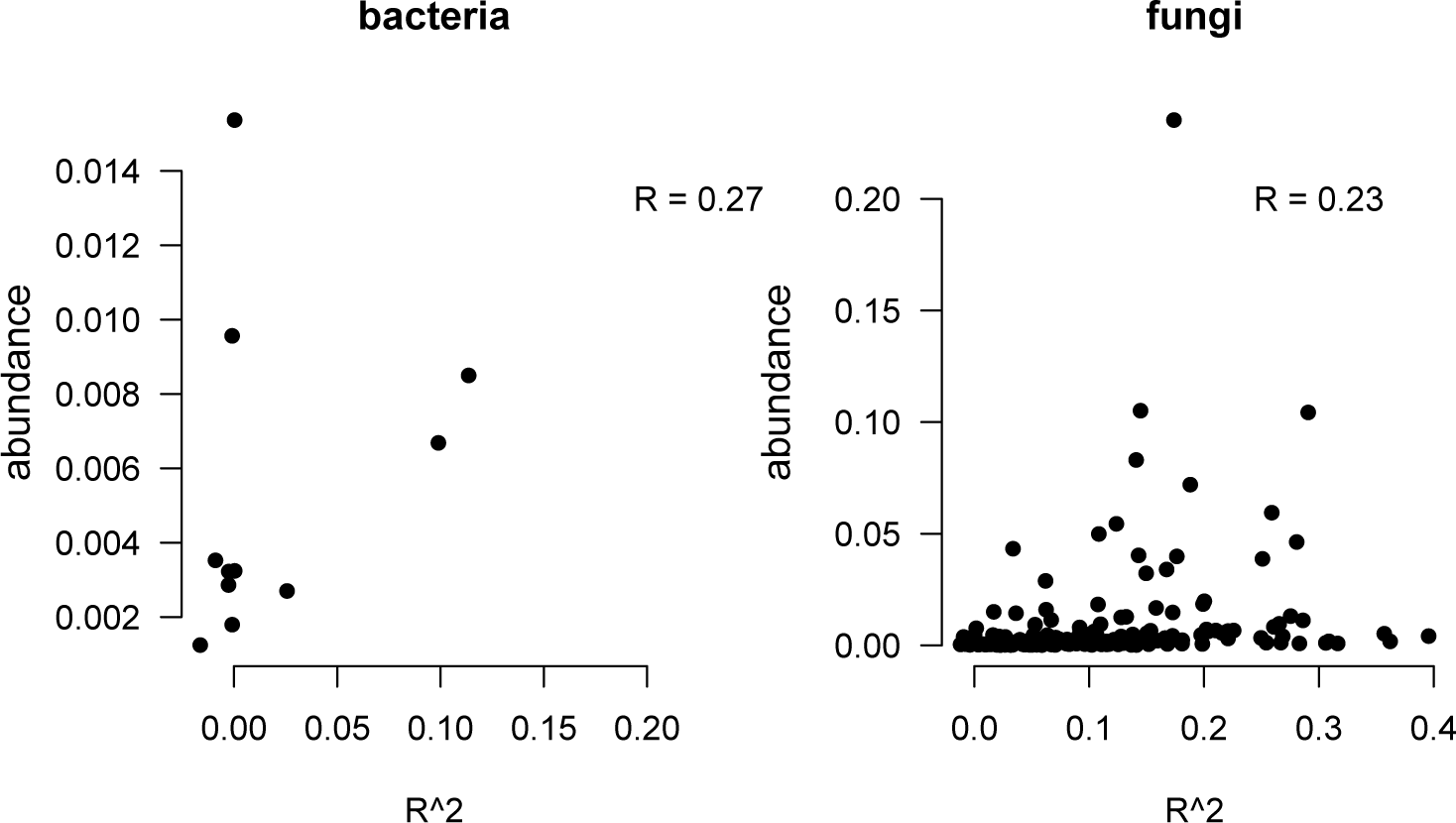
Relationship between *R*^2^ values of random forest models for the relative abundance of each microbial taxon (Hellinger standardized data) and mean relative abundance. This relationship shows if model performance was influenced by taxon relative abundance. Pearson’s correlation coefficients are shown. The correlation between fungal abundance and *R*^2^ was significant, but the correlation between bacterial abundance and *R*^2^ was not (sample size differed drastically between these comparisons, thus influencing p values, see main text).

**Figure S13:**
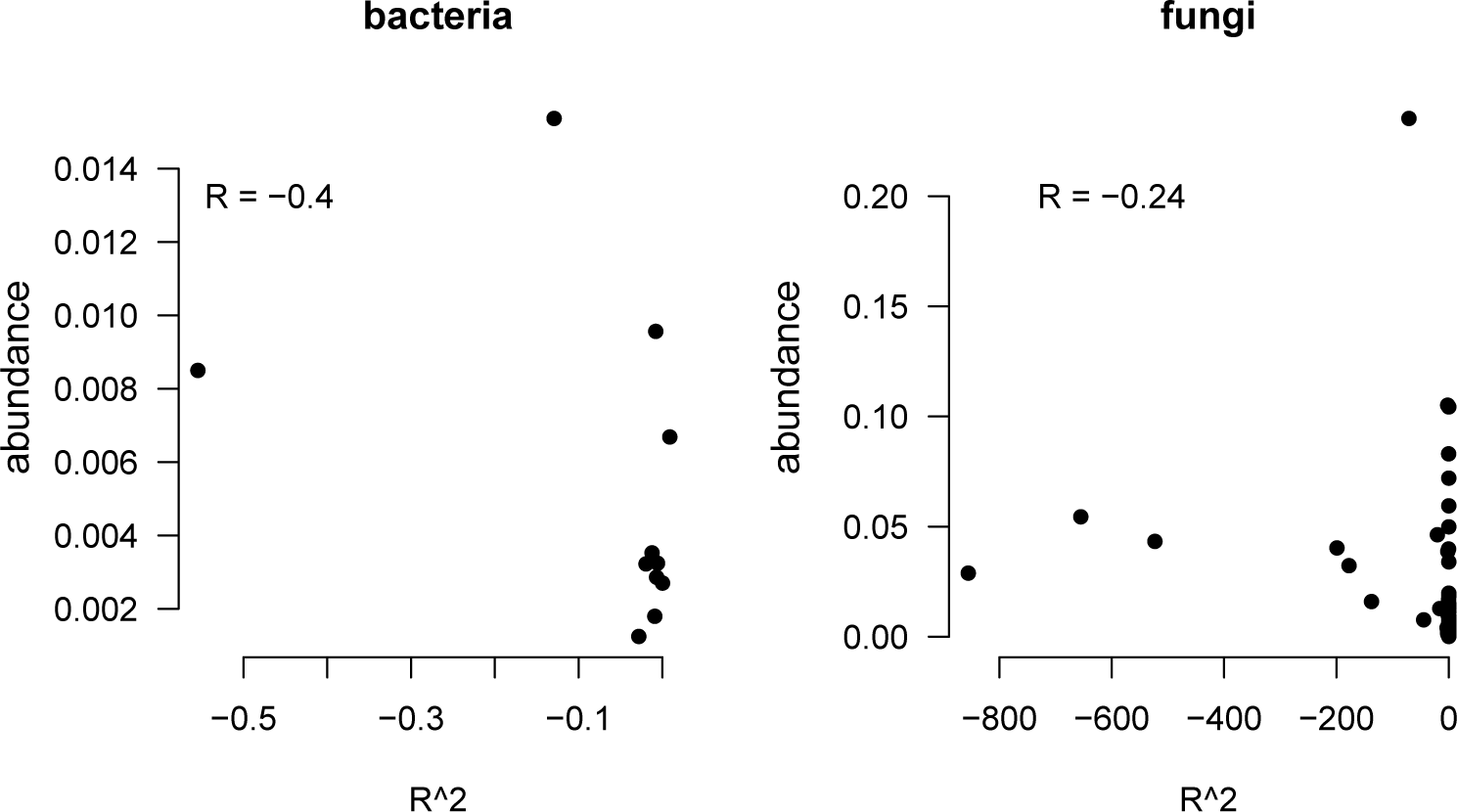
Relationship between *R*^2^ values of random forest models for the absolute abundance of each microbial taxon (abundances normalized by the ISD) and mean absolute abundance. This relationship shows if model performance was influenced by taxon abundance. Pearson’s correlation coefficients are shown. The correlation between fungal abundance and *R*^2^ was significant, but the correlation between bacterial abundance and *R*^2^ was not (sample size differed drastically between these comparisons, thus influencing p values, see main text).

**Figure S14:**
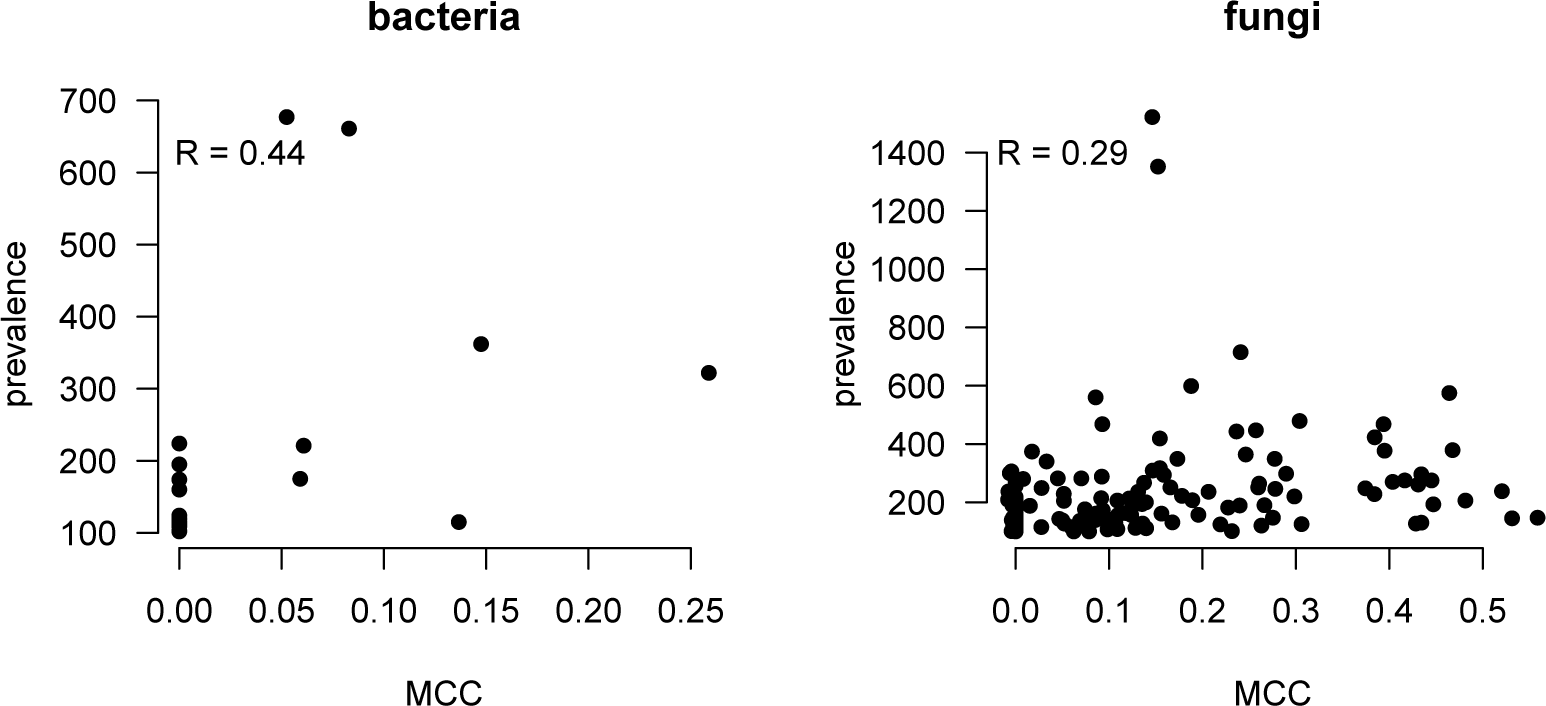
Relationship between Matthew’s Correlation Coefficient (MCC) values of random forest models for the occurrence of each microbial taxon and prevalence (how many samples the microbial taxon was observed within). Pearson’s correlation coefficients are shown. The correlation between fungal abundance and *R*^2^ was significant (p < 0.01), but the correlation between bacterial abundance and *R*^2^ was not (p = 0.11; sample size differed drastically between these comparisons, thus influencing p values, see main text).

**Figure S15:**
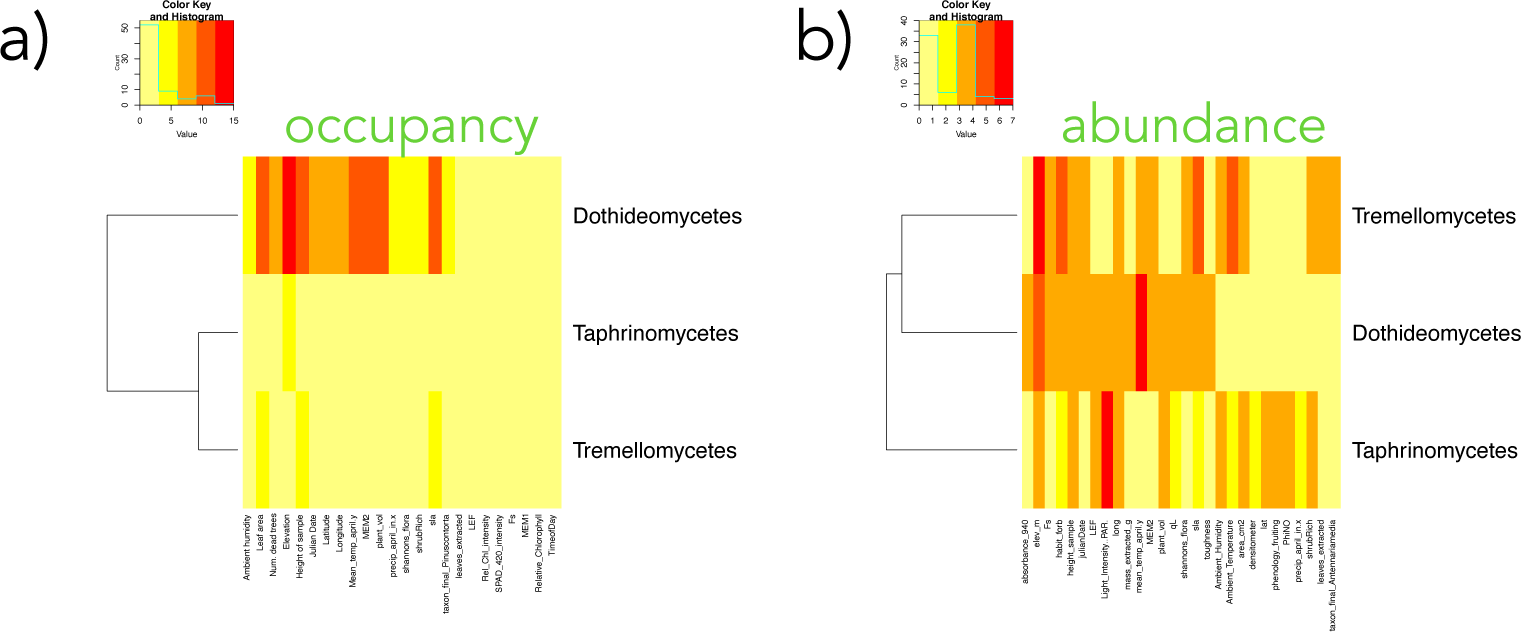
Heatmaps of feature importance for models of fungal occupancy (a) and absolute abundance (b). Heatmaps were not generated for bacteria, because bacterial assemblage variation was generally unpredictable. Features chosen were in the top ten most important for successful models (those models with an R sq over 1% or an MCC greater than 0.2) and were important for at least 20% of all successful models (thus those features that were important for isolated taxa are not shown here, for the sake of visualization).

**Figure S16:**
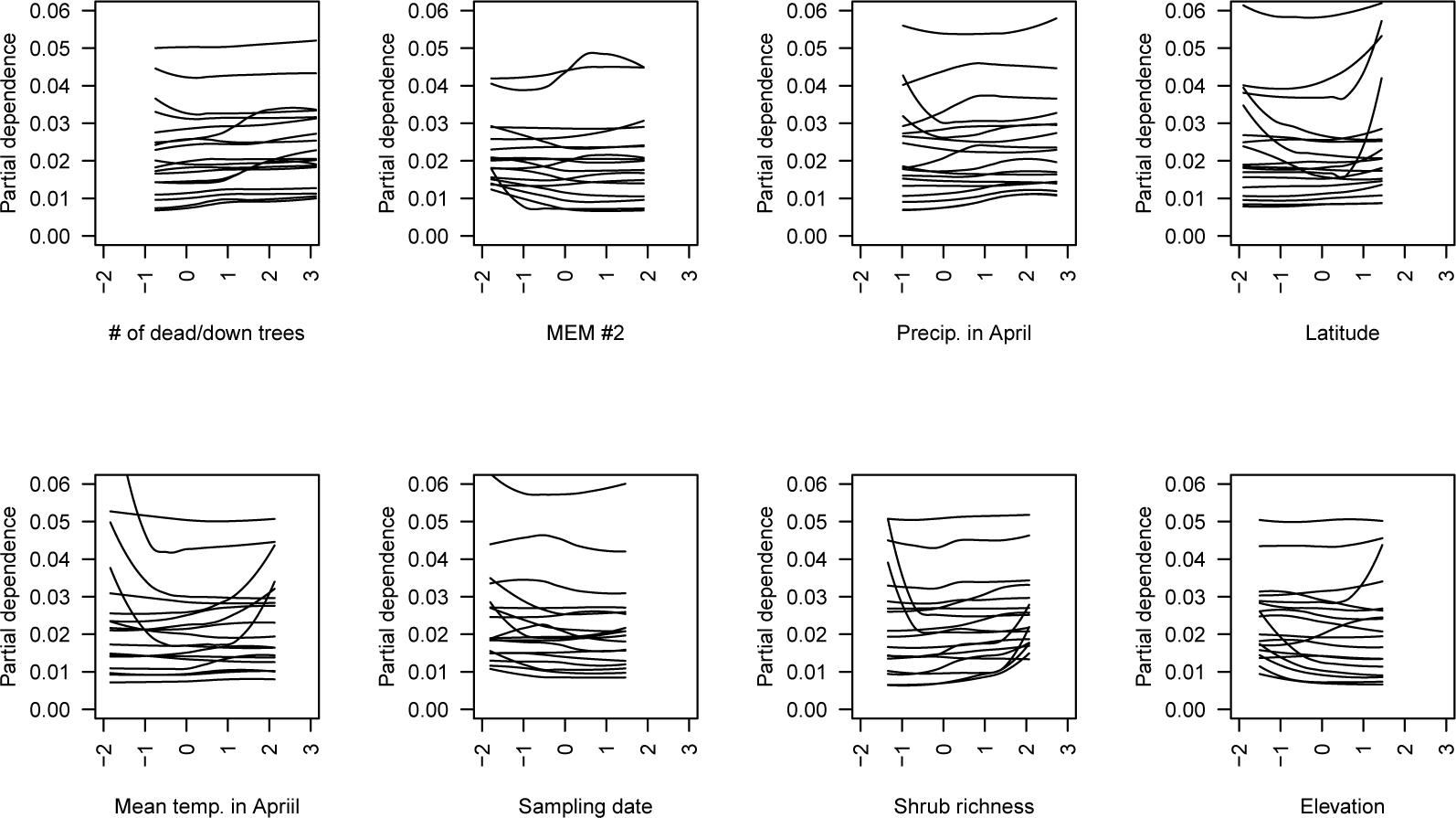
Partial dependence plots showing the modeled relationship between each feature (x axis; z score standardized) and the relative abundance of a fungal taxon (y axis). Each line shows the response curve for a different fungal taxon. Feature-taxon relationships shown are from the best performing models, each of which had an *R*^2^ *>* 0.25. Axis dimensions are standardized among plots to aid visual comparison. Because so few models were successful for fungal absolute abundances and bacteria we omit analogous figures for them.

**Figure S17:**
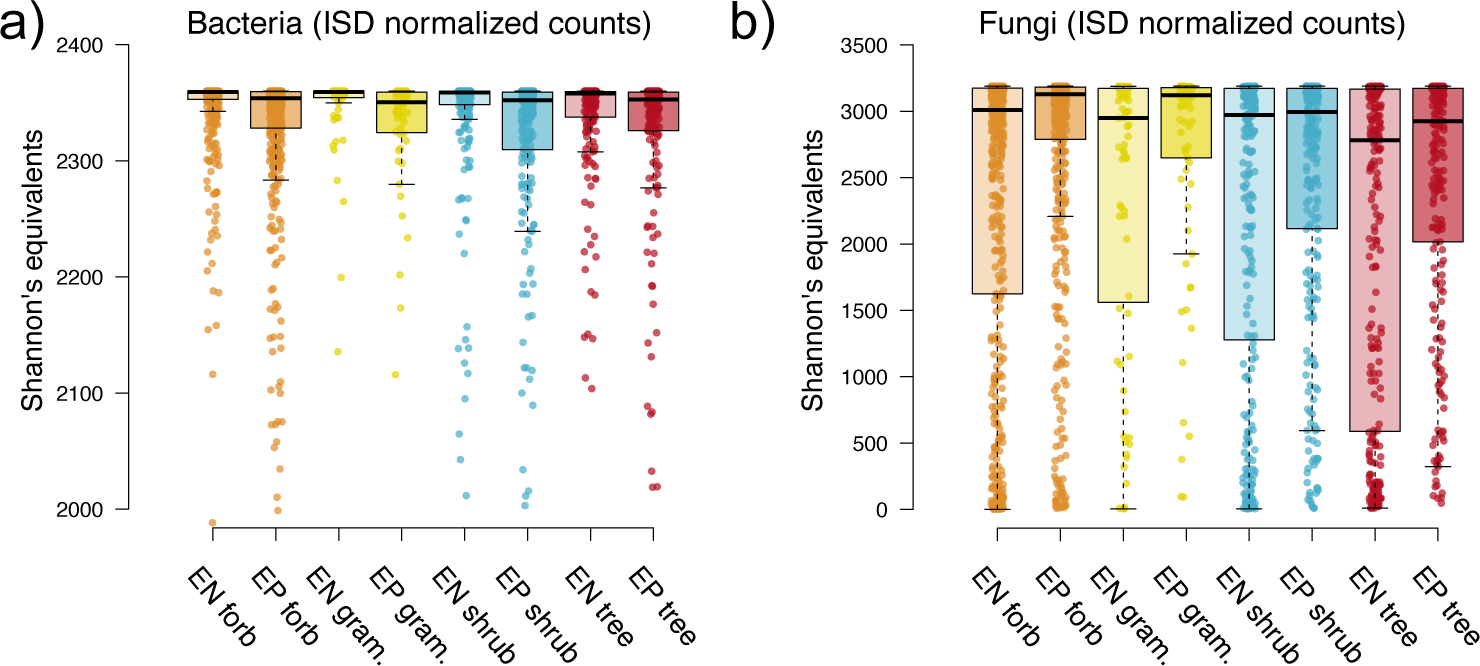
Shannon’s diversity of bacterial (a) and fungal (b) assemblages from epiphyte (EP) and endophyte (EN) samples of various plant taxa grouped by growth habit (forb, graminoid, shrub, or tree). Diversity estimates were exponentiated to convert them into species equivalents. Boxplots denote interquartile ranges, with a horizontal bar illustrating the median. Whiskers extend from the 10th to the 90th quantiles of the data. For Tukey’s HSD comparisons among groups see Tables S10 and S11.

**Figure S18:**
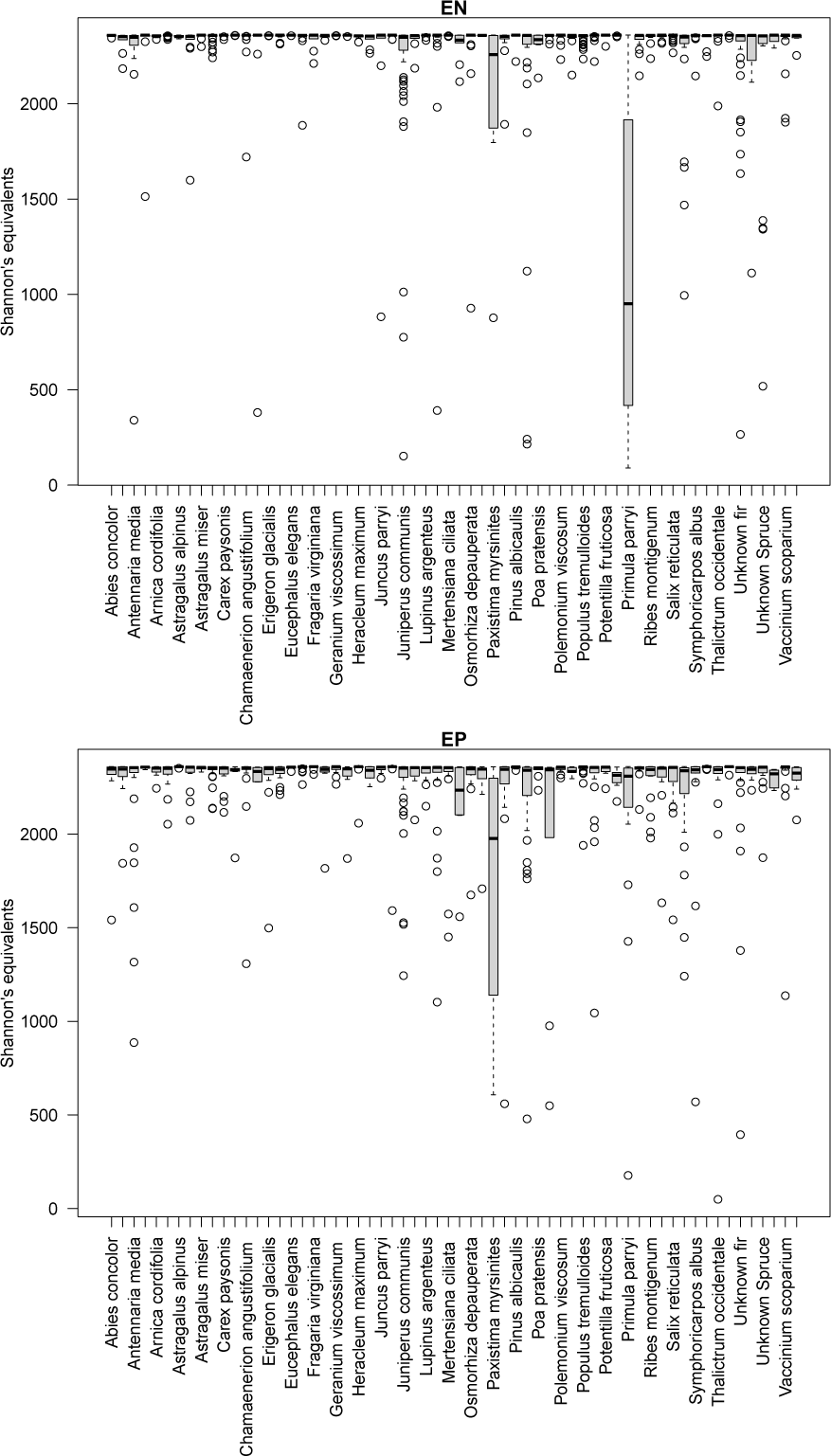
Shannon’s diversity of bacterial assemblages by host for epiphyte (EP) and endophyte (EN) samples. Diversity estimates were exponentiated to convert them into species equivalents and were calculated from ISD normalized count data. Boxplots denote interquartile ranges, with a horizontal bar illustrating the median. Whiskers extend from the 10th to the 90th quantiles of the data.

**Figure S19:**
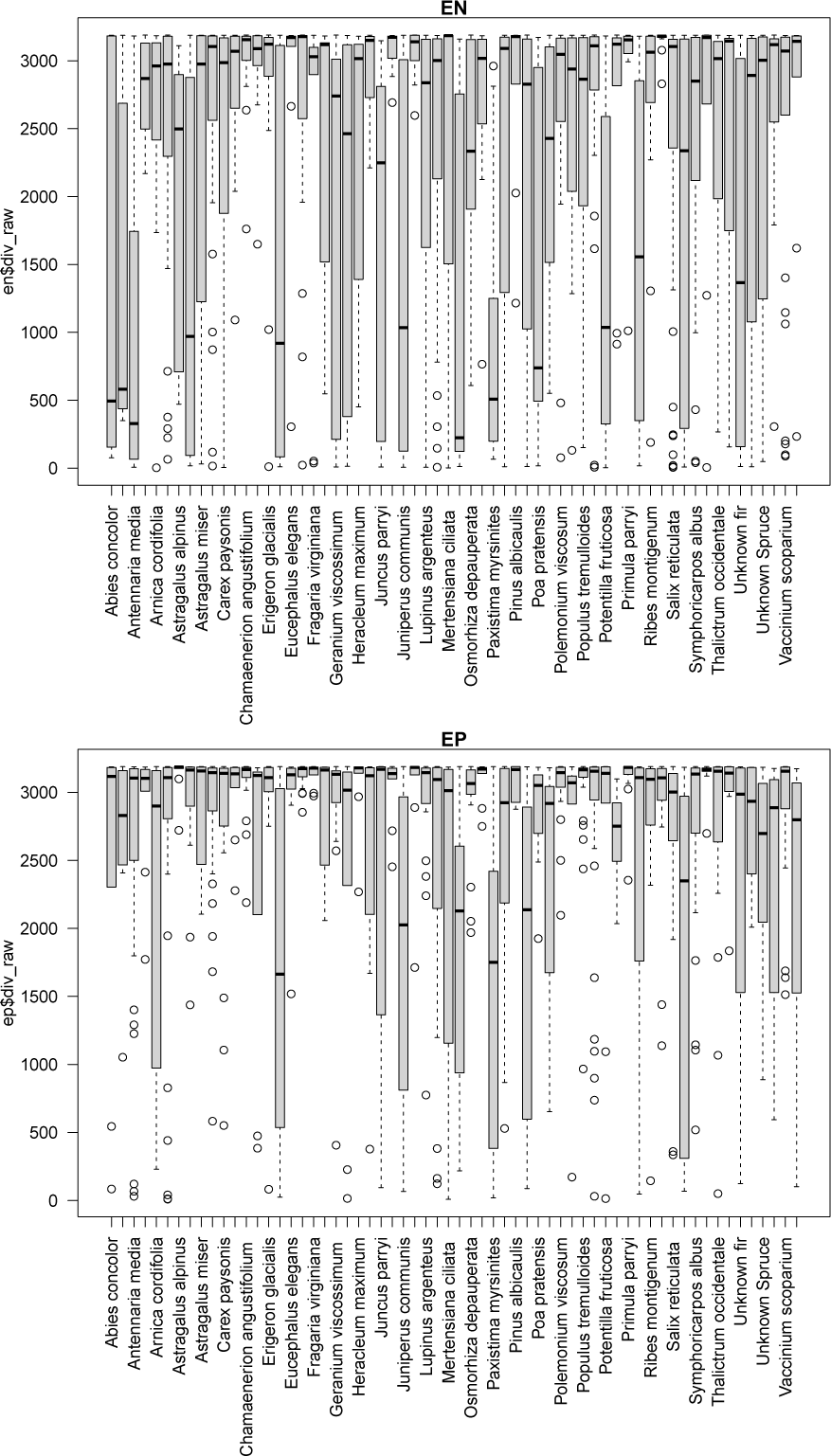
Shannon’s diversity of fungal assemblages by host for epiphyte (EP) and endophyte (EN) samples. Diversity estimates were exponentiated to convert them into species equivalents and were calculated from ISD normalized count data. Boxplots denote interquartile ranges, with a horizontal bar illustrating the median. Whiskers extend from the 10th to the 90th quantiles of the data.

**Figure S20:**
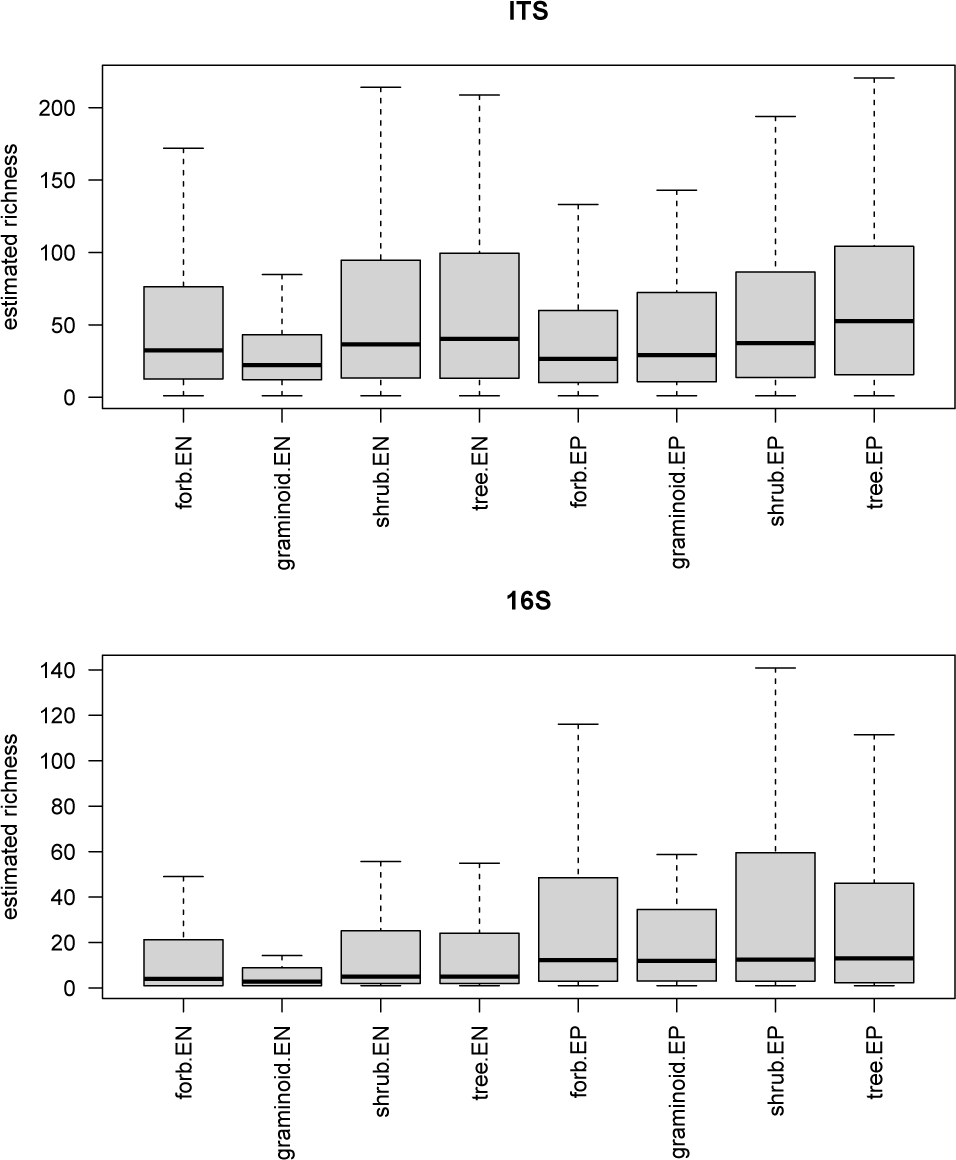
Estimated richness of ITS and 16S OTUs as a function of host compartment and life history. For details of richness estimation see main text. Results from a Tukey’s HSD test are shown in Tables S13 and S12.

**Figure S21:**
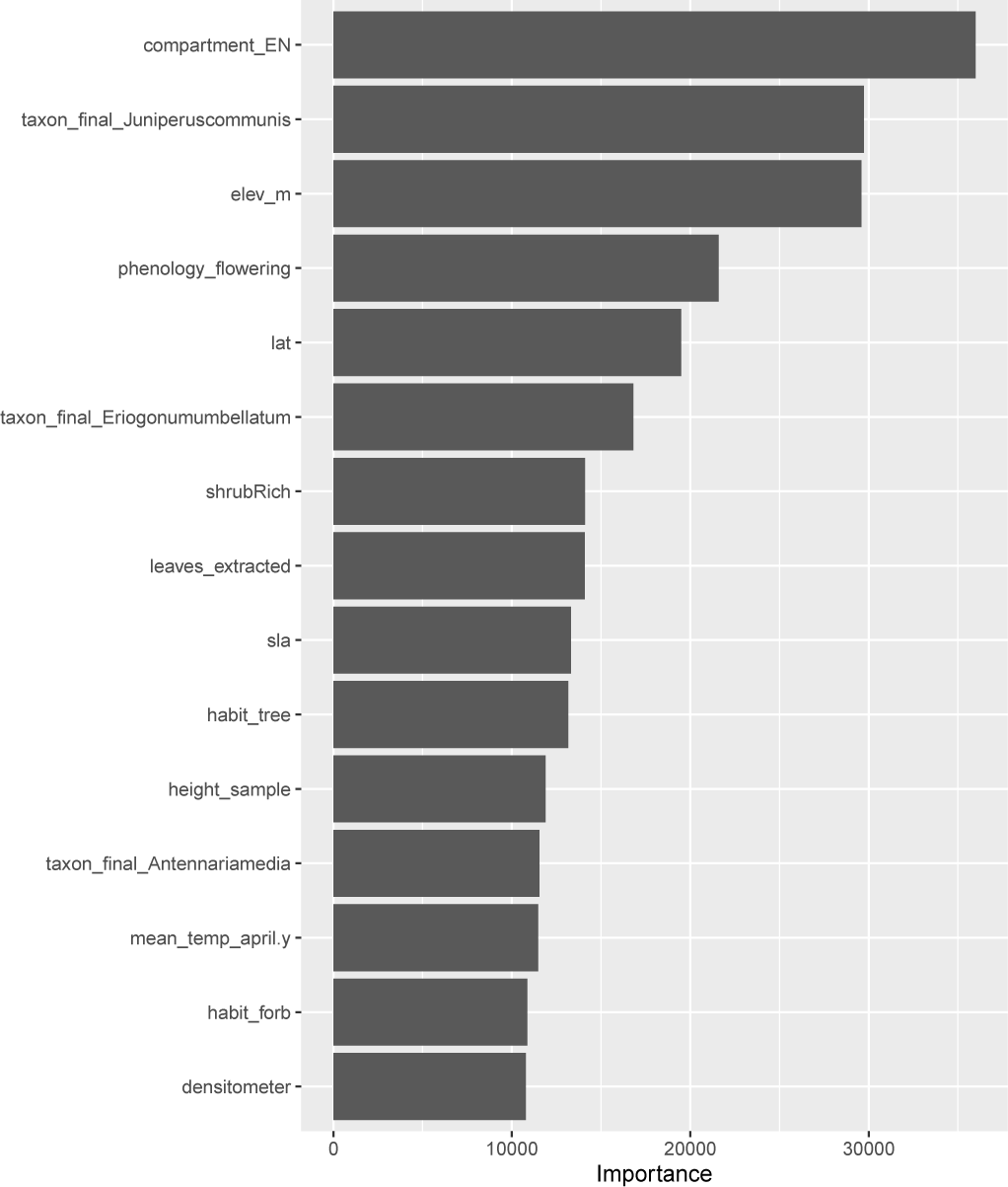
Feature importance plot for a random forest model of fungal Shannon’s diversity.

**Figure S22:**
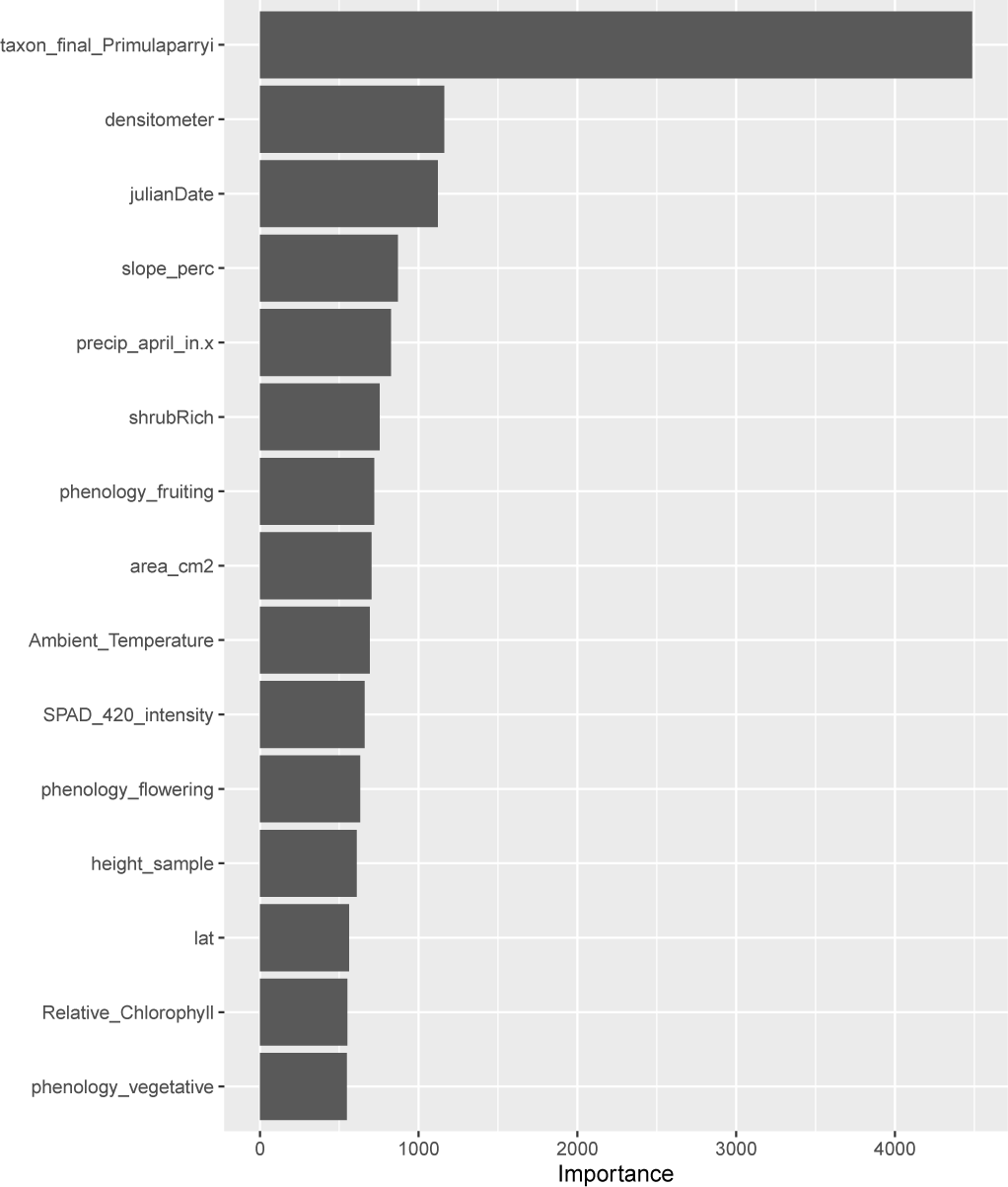
Feature importance plot for a random forest model of bacterial Shannon’s diversity.

**Figure S23:**
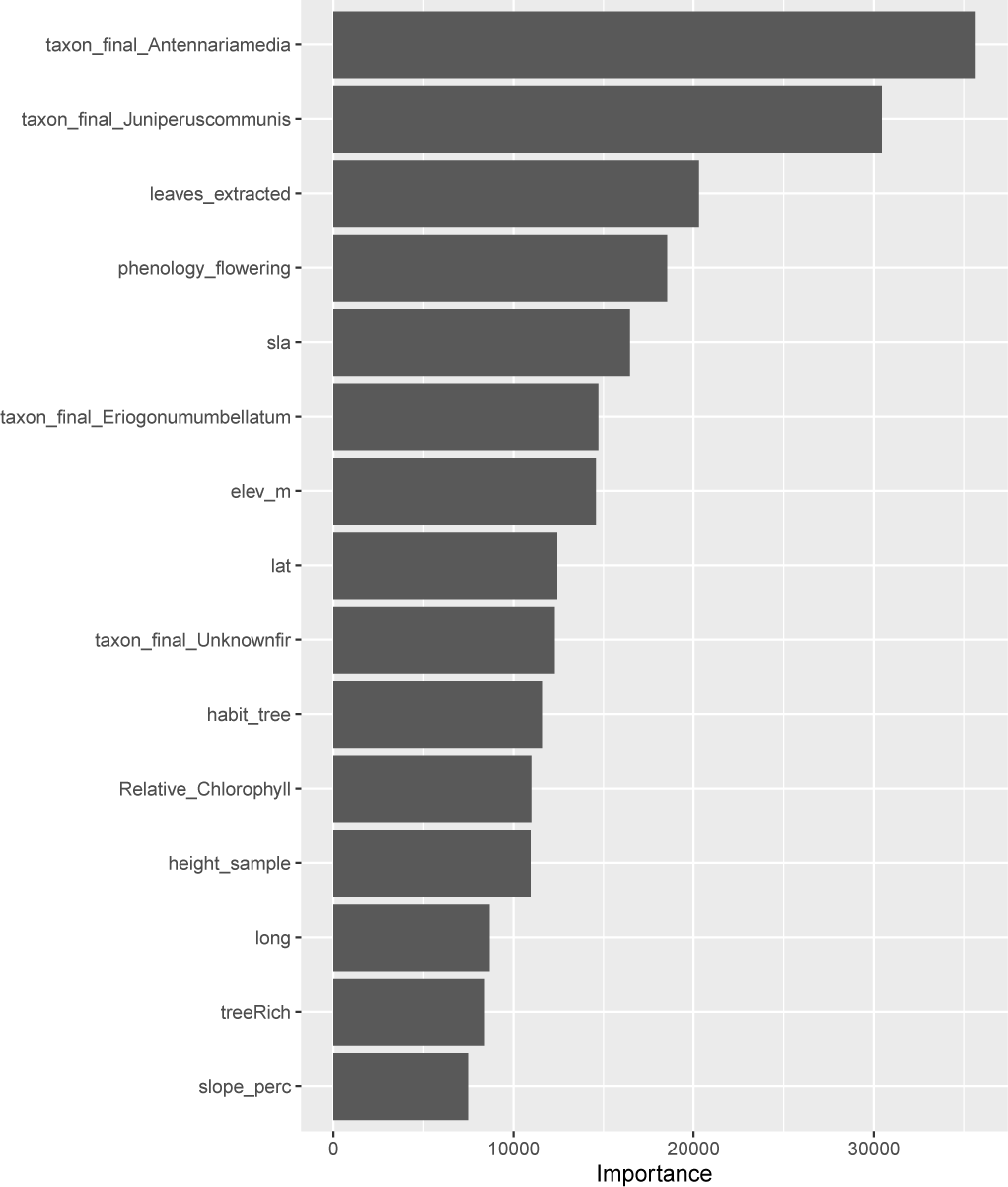
Feature importance plot for a random forest model of fungal endophyte Shannon’s diversity.

**Figure S24:**
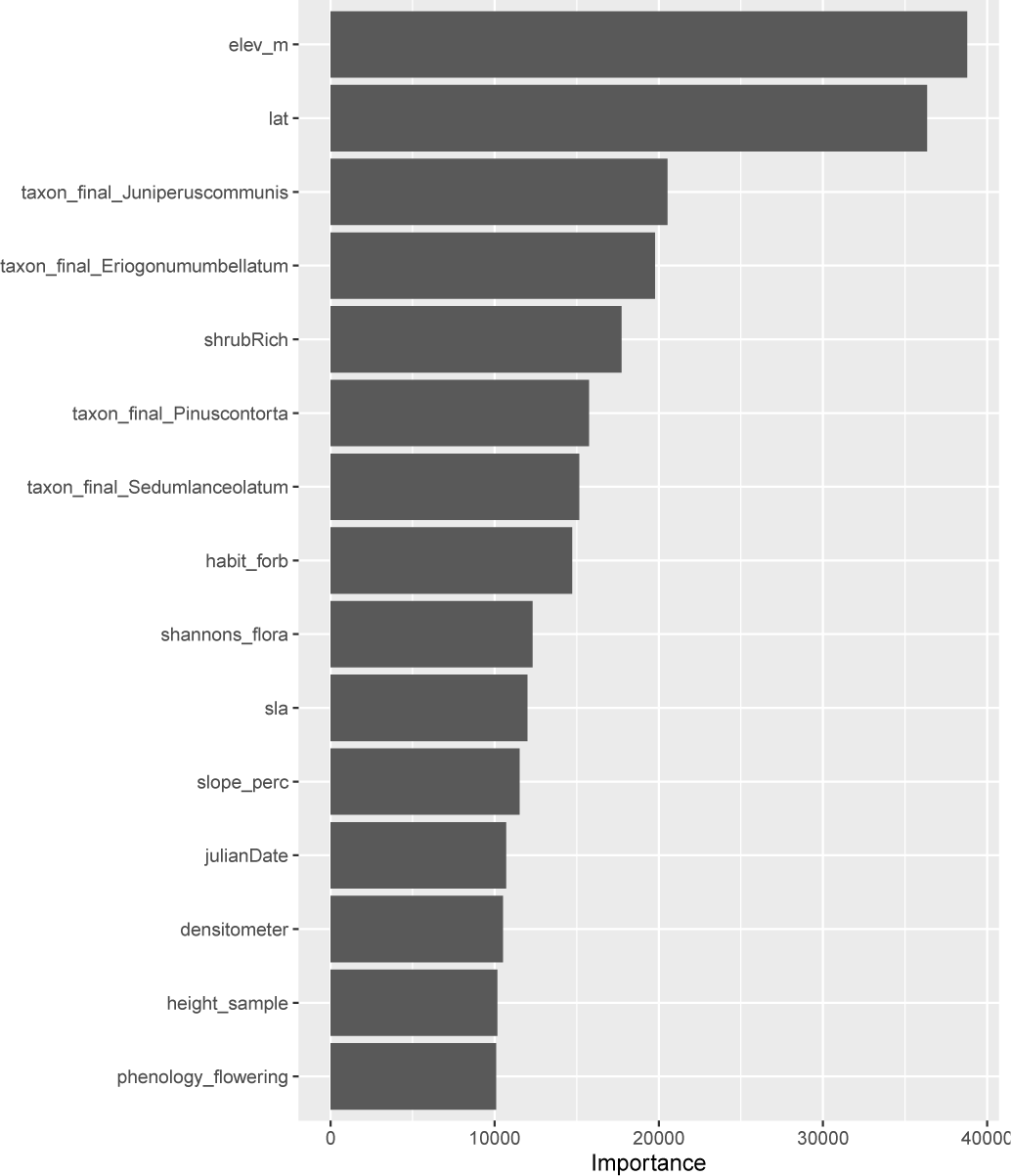
Feature importance plot for a random forest model of fungal epiphyte Shannon’s diversity.

**Figure S25:**
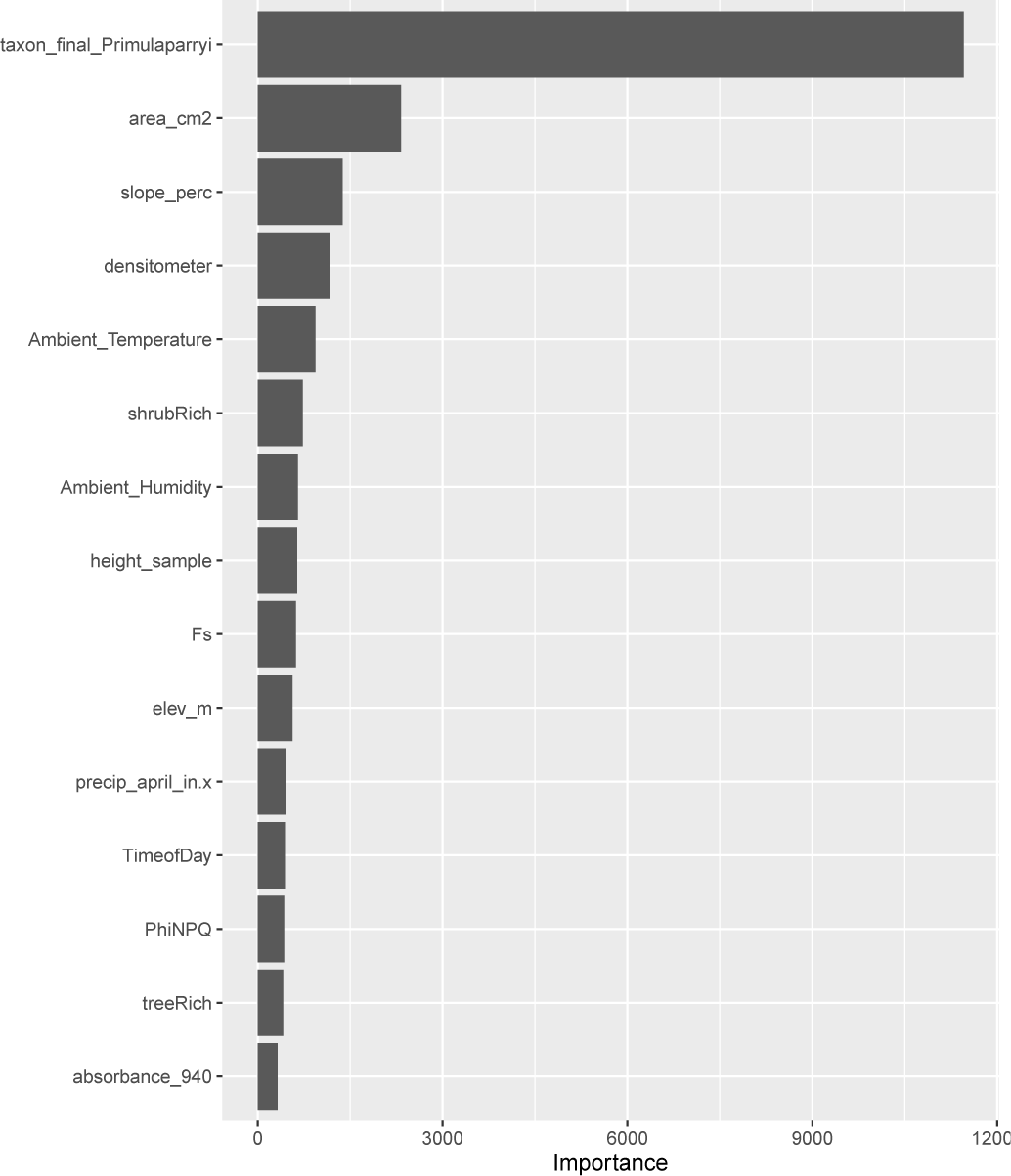
Feature importance plot for a random forest model of bacterial endophyte Shannon’s diversity.

**Figure S26:**
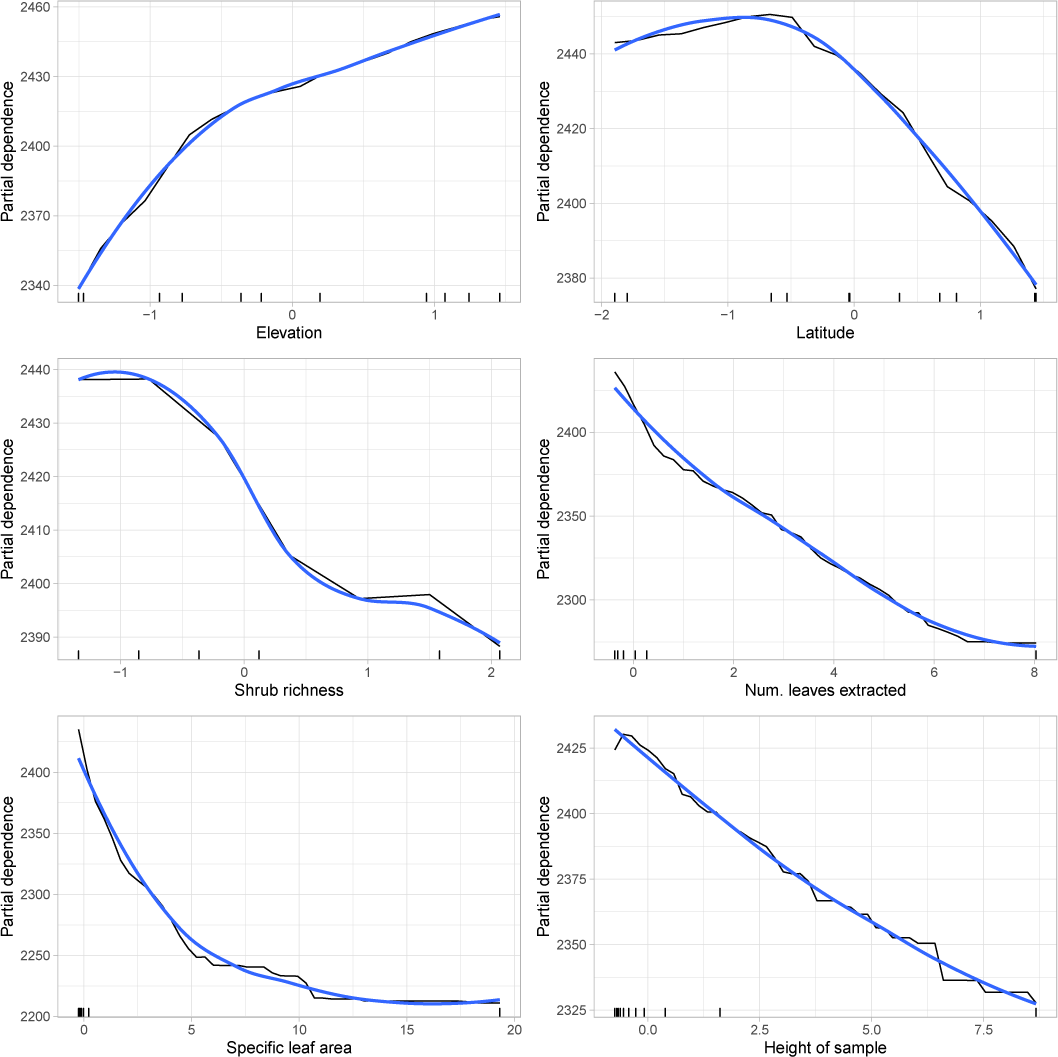
Partial dependence plot (PDP) of the most influential continuous features used by a random forest model of fungal Shannon’s diversity. PDPs show the relationship between a feature and the response across both of their ranges.

**Figure S27:**
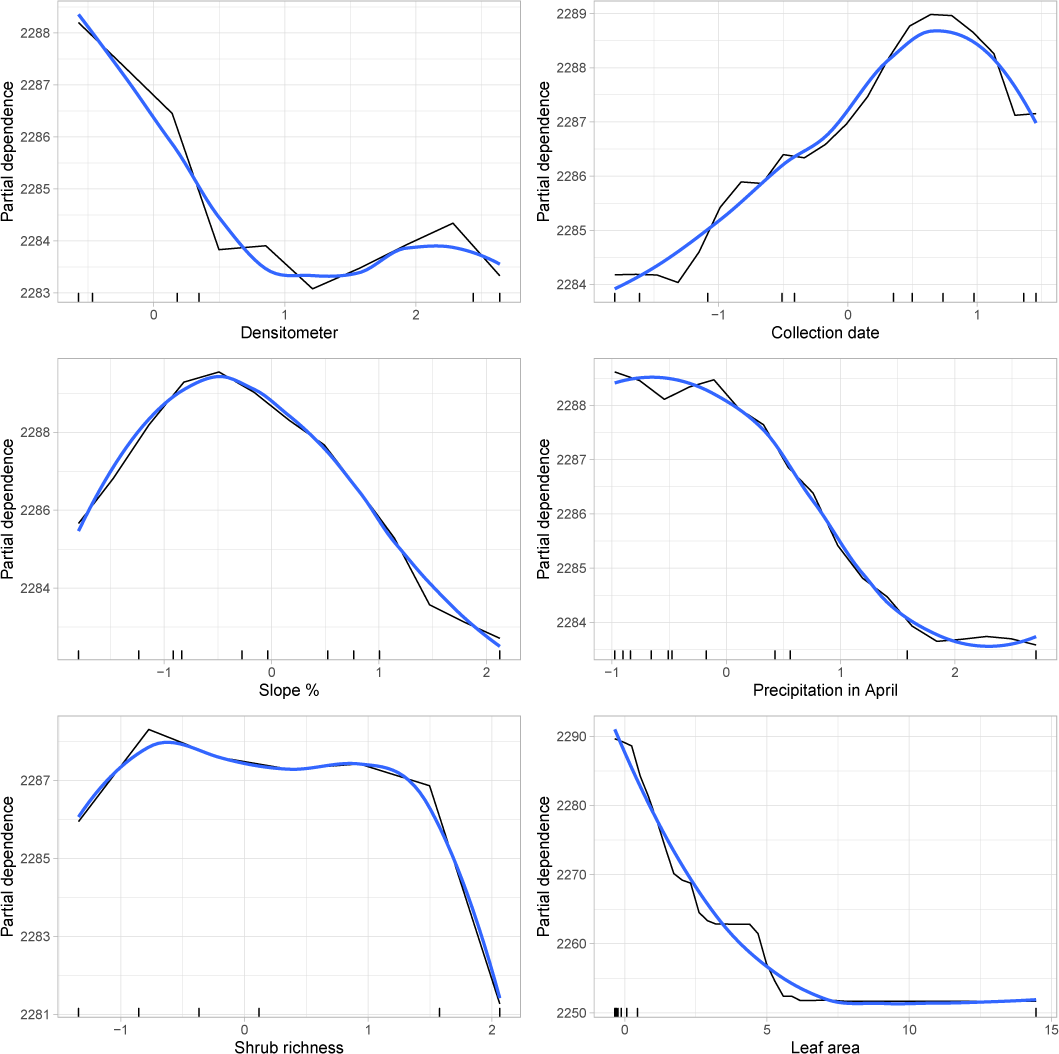
Partial dependence plot (PDP) of the most influential continuous features used by a random forest model of bacterial Shannon’s diversity. PDPs show the relationship between a feature and the response across both of their ranges.

**Figure S28:**
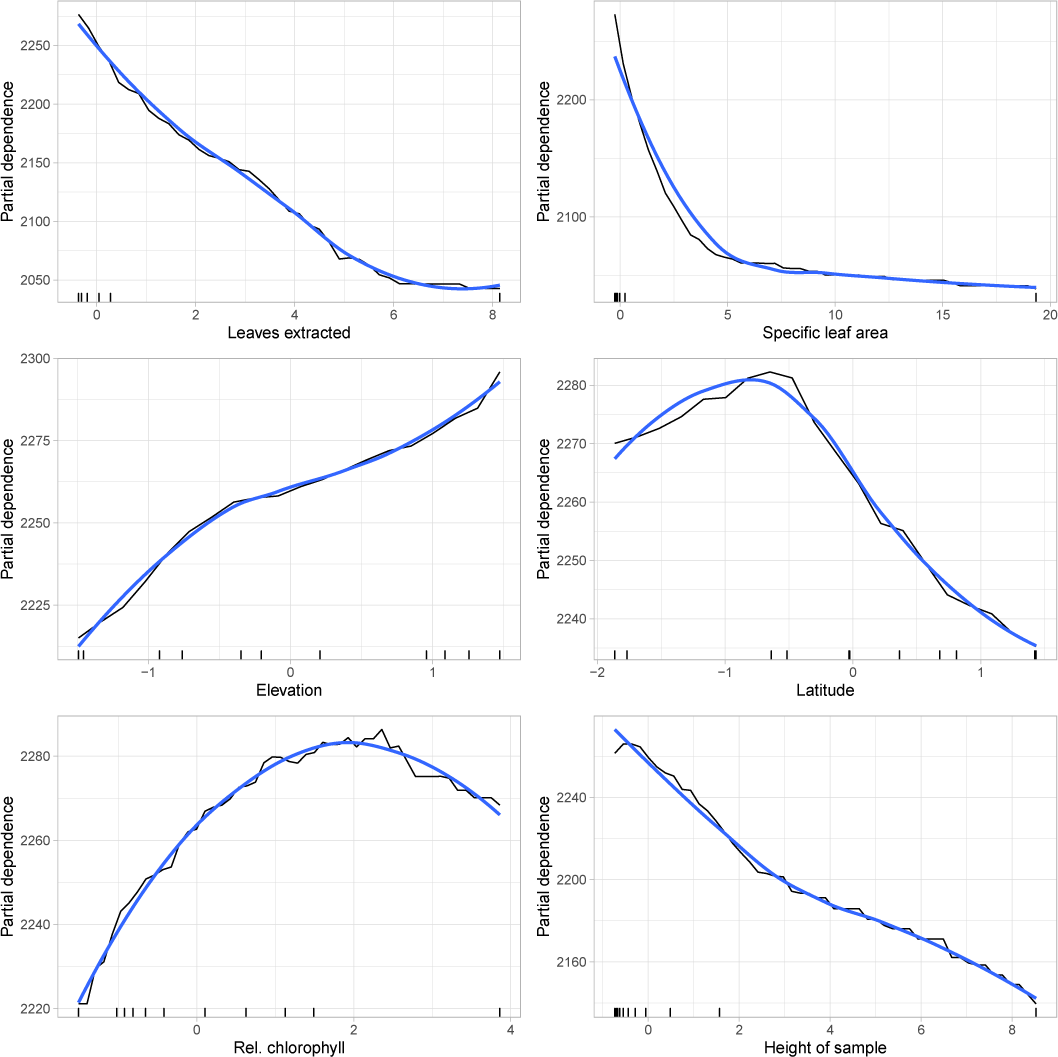
Partial dependence plot (PDP) of the most influential continuous features used by a random forest model of fungal endophyte Shannon’s diversity. PDPs show the relationship between a feature and the response across both of their ranges.

**Figure S29:**
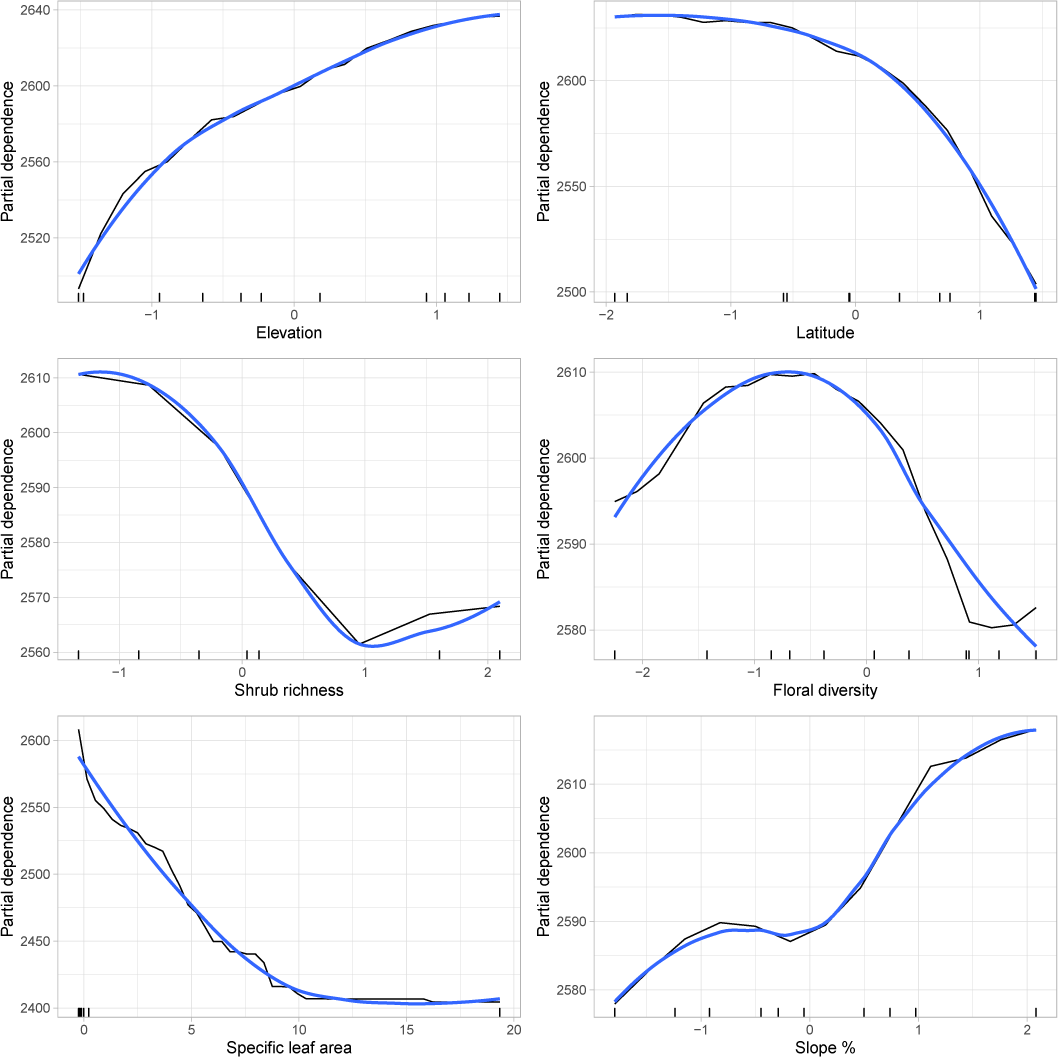
Partial dependence plot (PDP) of the most influential continuous features used by a random forest model of fungal epiphyte Shannon’s diversity. PDPs show the relationship between a feature and the response across both of their ranges.

**Figure S30:**
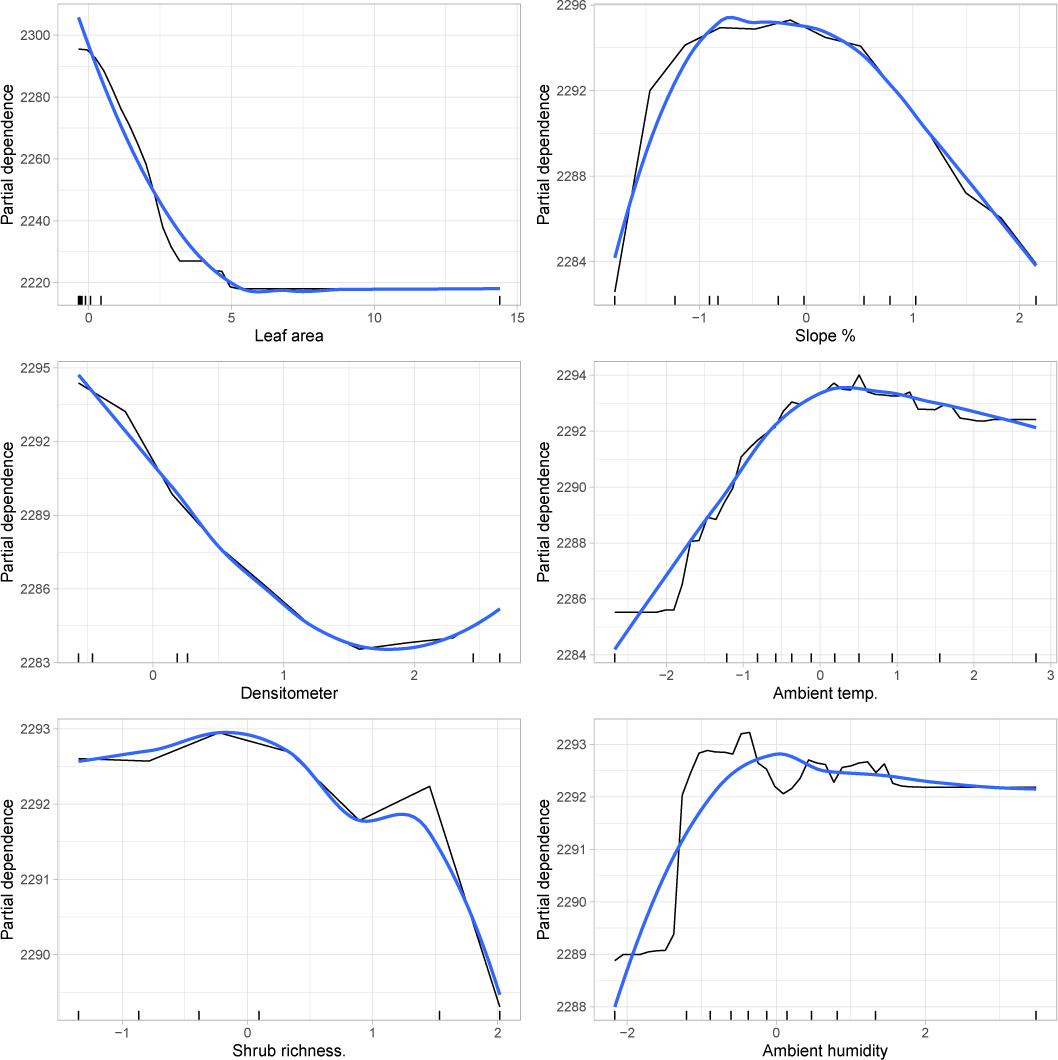
Partial dependence plot (PDP) of the most influential continuous features used by a random forest model of bacterial endophyte Shannon’s diversity. PDPs show the relationship between a feature and the response across both of their ranges. A PDP from the model of bacterial ephiphyte diversity is not provided because the model had poor predictive performance.

**Figure S31:**
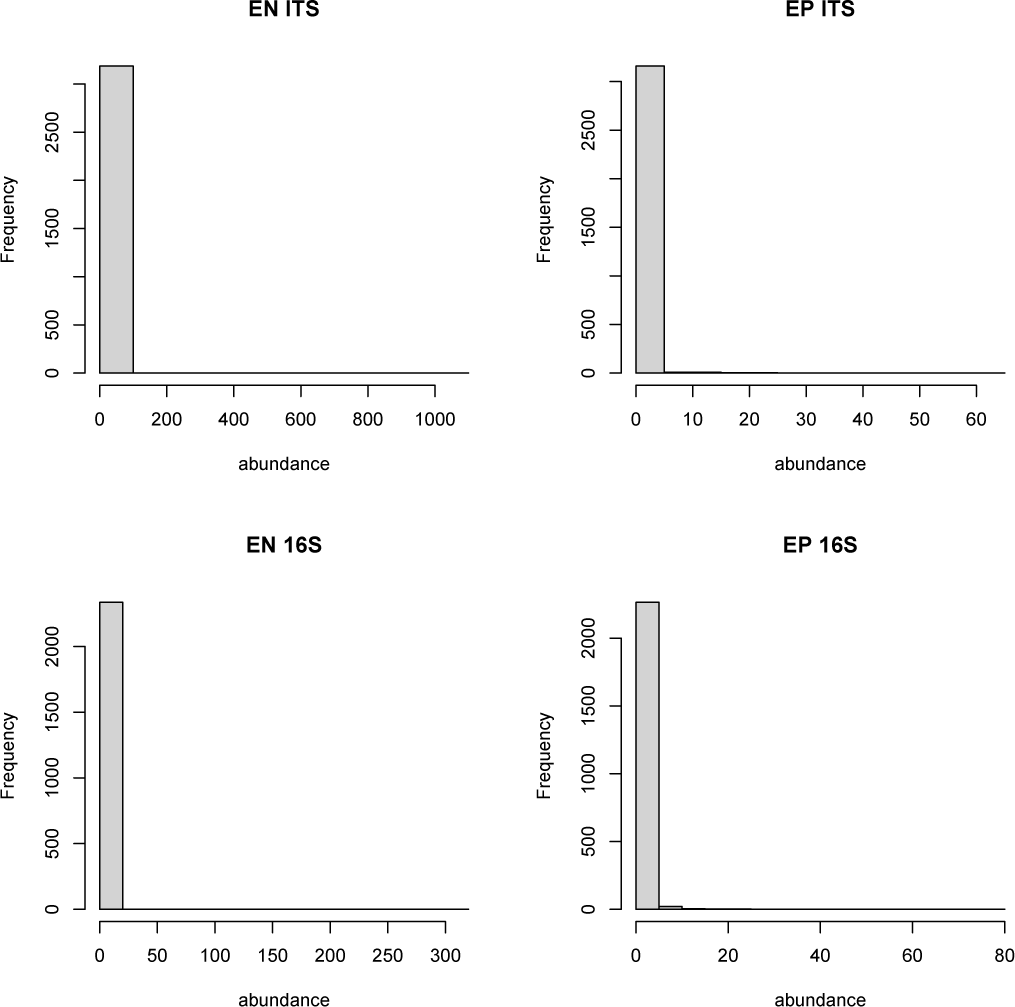
Rank abundance histograms for ISD transformed data. Locus and compartment are shown in each plot. Frequency refers to the number of taxa within a given abundance class. A single random sample was chosen from each sampling location and host, thus some hosts are represented more than once since they occurred at multiple sites.

**Figure S32:**
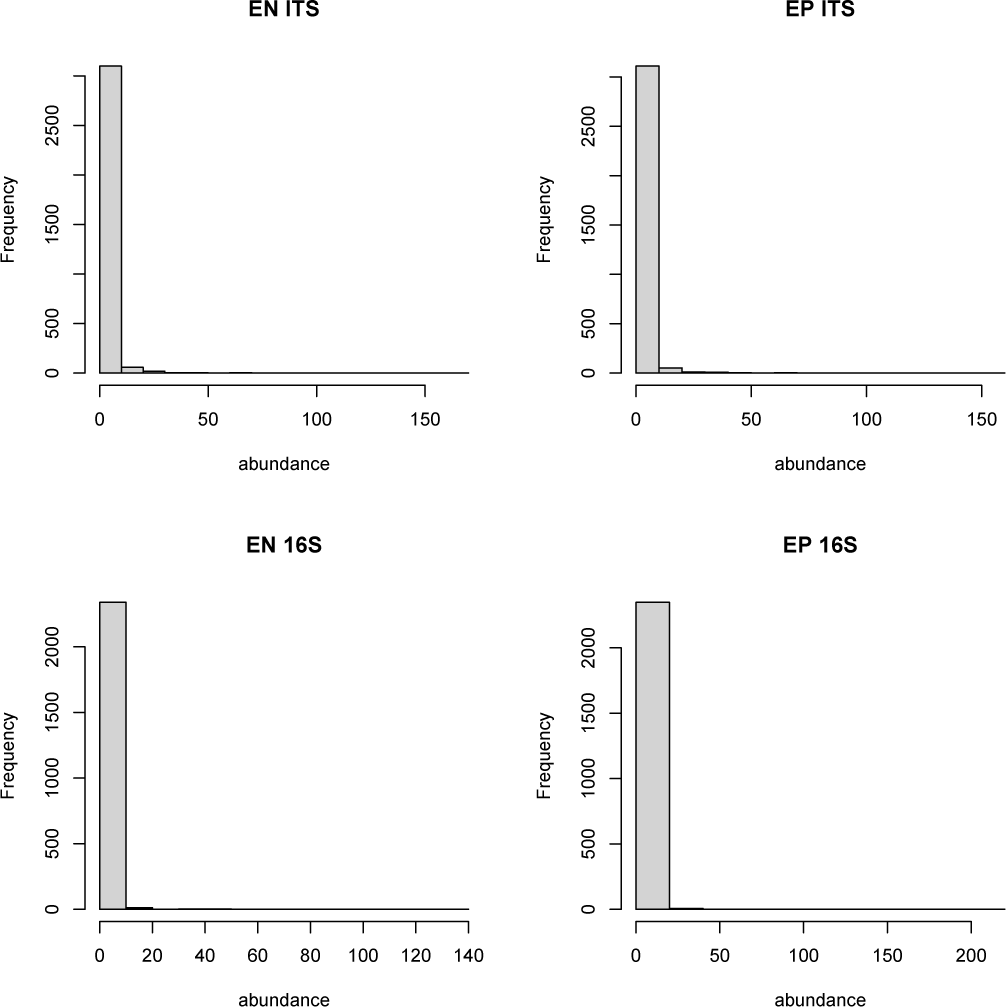
Rank abundance histograms for Hellinger transformed data. Locus and compartment are shown in each plot. Frequency refers to the number of taxa within a given abundance class. A single random sample was chosen from each sampling location and host, thus some hosts are represented more than once since they occurred at multiple sites.

